# CD24 Acts as an Evolutionarily Conserved Innate Immune Checkpoint in Colorectal Cancer

**DOI:** 10.64898/2026.08.25.746757

**Authors:** Ana B. Machado, Cátia Rebelo de Almeida, Catarina M. Azevedo, Nuno Viana, Inês Trancoso, Diogo R. Fernandes, Vanda Póvoa, Francisco Marques, Rita Zilhão, June Ereño-Orbea, Jesús Jiménez-Barbero, Ana Rita Carlos, Salomé S. Pinho, Rita Fior

## Abstract

Checkpoint immunotherapy has transformed cancer treatment, yet current approaches targeting adaptive immunity benefit only a subset of patients, leaving innate immunity as a largely untapped therapeutic frontier. Here, we identify CD24 as an innate immune checkpoint that protects colorectal tumors from macrophage-mediated clearance through an evolutionarily conserved recognition mechanism. Using zebrafish xenografts of isogenic colorectal cancer (CRC) cell lines, SW480 and SW620, we show that high CD24 expression in SW620 correlates with an immune-evasive, macrophage-resistant phenotype. Loss of human CD24 dramatically sensitizes tumors to clearance in zebrafish, while pharmacological macrophage depletion abolishes this effect. Mechanistically, CD24 suppresses innate immunity in a multilayered fashion, by limiting myeloid recruitment, dampening TNFα-driven macrophage inflammatory polarization, and blocking phagocytosis. Live imaging further revealed that CD24 constrains macrophages to a restrained, patrol-like state, and that its loss enables them to adopt a highly motile, tumor–directed, and functionally engaged state, characterized by increased fusion activity and myeloid intercellular interactions. We show that zebrafish macrophages respond to human CD24 despite extensive evolutionary divergence, and glycocalyx profiling revealed broad remodeling of the tumor cell surface upon CD24 loss, suggesting evolutionary conservation of sialic acid-dependent receptor recognition. Transcriptomic analyses identified the Siglec-like gene *si:dkey-24p1.7* as a candidate zebrafish macrophage-expressed receptor mediating this response. Finally, analysis of TCGA CRC cohorts revealed that CD24 expression is a stage-dependent prognostic marker, underscoring the clinical relevance of this axis. Together, these findings establish CD24 as a critical orchestrator of innate immune evasion in CRC, while further validating zebrafish xenografts as a powerful platform for dissecting innate immuno-oncobiology *in vivo*.

**Graphical abstract:** 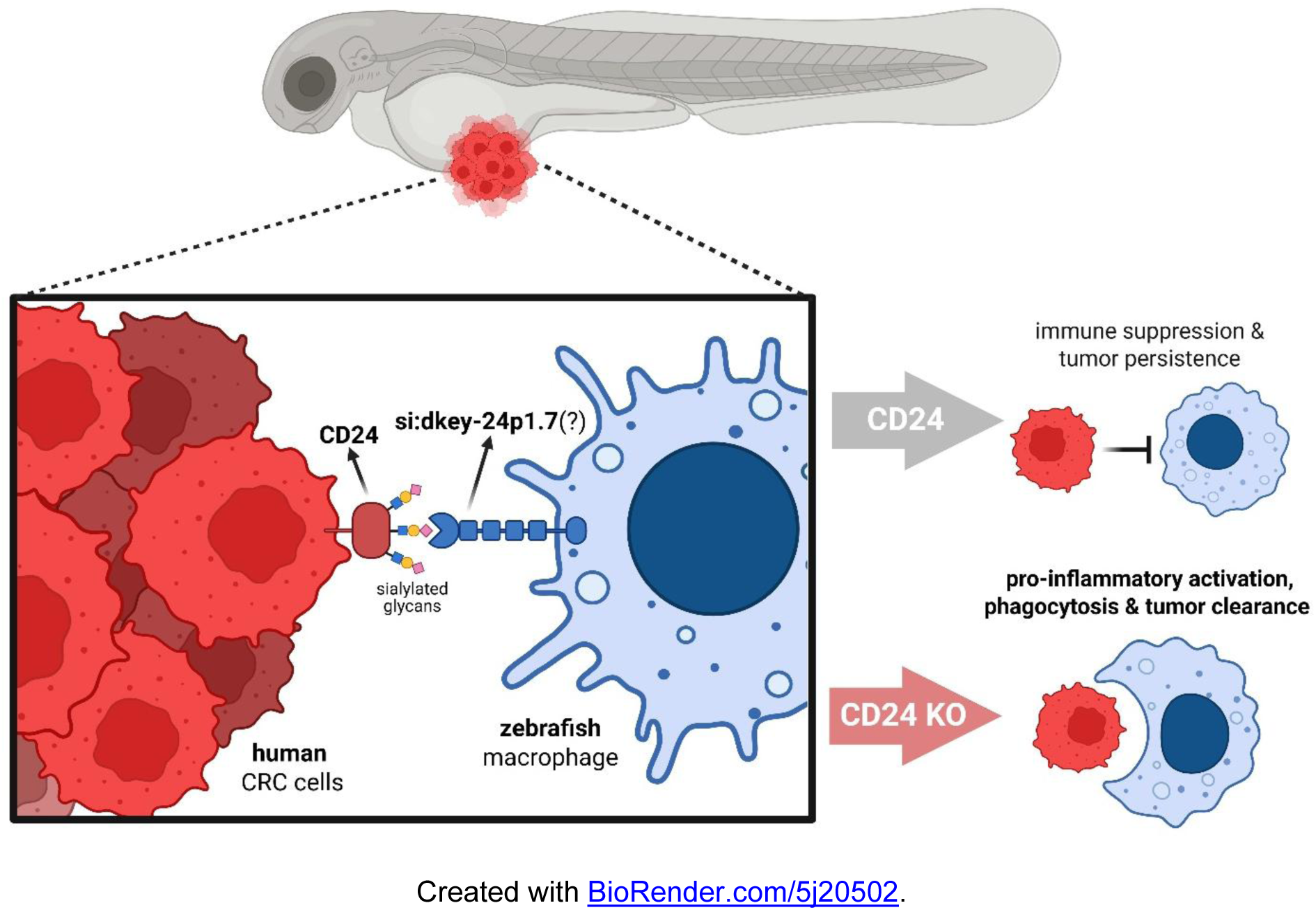

## Introduction

Although immune checkpoint blockade (ICB) has established that anti-tumor immunity can be therapeutically harnessed, many patients remain refractory or develop resistance^1,2^. The innate immune-myeloid compartment dominates the immune tumor microenvironment (TME) of most solid cancers, and plays a pivotal role in the earliest stages of tumor immune surveillance and in shaping subsequent adaptive responses^3,4^. Yet, while tumoral suppression of adaptive T cell immunity has been extensively characterized, cancer innate immune evasion remains comparatively underexplored. Overexpression of “don’t eat me” signals represents one of the first innate evasion strategies described in cancer, acting to suppress macrophage phagocytosis upon recognition by inhibitory receptors^5^. The CD47-signal-regulatory protein α (SIRPα) axis exemplifies this paradigm and has been defined as a canonical innate immune checkpoint^6^, as its blockade restores macrophage-mediated tumor clearance in preclinical models^7,8^. Clinical translation has been limited by dose-limiting toxicities arising from the broad expression of CD47 on nonmalignant cells^9,10^. These challenges underscore the need to identify and understand alternative mechanisms of innate immune evasion, unlocking therapeutic potential in patients who fail to respond to current immunotherapies.

The zebrafish xenograft model, in which human cancer cells are engrafted into 2-day post-fertilization (dpf) embryos^11^, is uniquely suited to identify and dissect innate immune mechanisms in cancer. Zebrafish adaptive immunity does not mature until 2–3 weeks post-fertilization^12–14^, creating a temporal window in which innate immune responses can be isolated without adaptive activity. Combined with optical transparency, genetic tractability, and conserved oncogenic and immune signaling pathways^15–18^, this system enables real-time *in vivo* imaging of tumor-immune dynamics at single-cell resolution. Exploiting this platform, we previously showed that isogenic colorectal cancer (CRC) cell lines, SW480 (primary tumor) and SW620 (lymph node metastasis)^19,20^, exhibit contrasting engraftment phenotypes in zebrafish xenografts: SW480 tumors are rapidly cleared (“regressors”), whereas SW620 tumors persist (“progressors”), in a manner critically dependent on host myeloid cells^21^. To identify molecular determinants underlying this innate immune evasion phenotype, we performed a comprehensive reanalysis of bulk RNA sequencing data of tumors isolated from progressor and regressor xenografts. Differential expression analysis revealed CD24 among the most significantly upregulated genes in SW620 progressors.

CD24 is a small, heavily glycosylated glycosylphosphatidylinositol (GPI)-anchored surface protein bearing up to 16 *O*- and *N*-linked glycosylation sites^22^, and has long been recognized as a cancer-associated antigen overexpressed across multiple malignancies^23^, including CRC^24^. Under physiological conditions, CD24 functions as a regulator of innate immune tolerance to tissue damage: upon cellular injury, damage-associated molecular patterns (DAMPs) – such as HMGB1, HSP70, and HSP90 – bind to CD24, which in turn recruits ITIM-bearing inhibitory receptors, including Siglec-10 and Siglec-5, expressed by immune cells^25,26^. Mechanistically, the Ig-like V-type domain of Siglecs specifically recognizes the sialylated glycans coating CD24, triggering SHP-1/SHP-2-dependent inhibitory signaling that dampens Toll-like receptor (TLR)-mediated inflammation^27^.

In cancer, this pathway is hijacked to evade immune surveillance, such that CD24 acts as a “don’t eat me” signal: engagement of tumor cell-expressed CD24 by Siglec-10 on tumor- associated macrophages establishes an innate immune checkpoint that suppresses phagocytosis^28^. Consistently, CD24 blockade in preclinical models of breast and ovarian cancer was shown to reduce tumor burden and prolong survival in a macrophage-dependent manner^28^. In CRC liver metastasis, neutrophil extracellular traps were shown to promote macrophage phagocytosis by downregulating CD24 in cancer cells^29^ and, in pancreatic cancer, CD24 has been identified as a resistance mechanism to KRAS inhibition^30^. In addition, CD24 can function as a sialyl-Lewis x–dependent ligand for P- and E-selectin on activated platelets and endothelium, thereby promoting leukocyte and tumor cell rolling that facilitates hematogenous metastasis^31–33^. It similarly engages L1CAM, which in cis forms a trimolecular complex with NCAM to support tumor cell migration^34,35^. CD24 further partitions into membrane lipid rafts^36^, where it may engage Src-family kinases such as Lyn, modulating CXCR4 signaling to reinforce invasive programs^37,38^.

The CD24-Siglec-10 immune checkpoint axis is critically dependent on sialylated glycans expressed by CD24, highlighting the importance of altered cellular glycosylation as a hallmark of cancer^39^ and as an active driver of immune suppression and evasion^40,41^. In CRC, increased cell surface sialylation has been shown to modulate macrophage function via the Siglec signaling pathways^42^. Accordingly, several therapeutic strategies targeting glycan-immune networks are emerging, with multiple approaches currently under clinical evaluation^40,43–47^, including combinations with PD-1/PD-L1 blockade^48–50^. In parallel, anti-CD24 antibodies have recently entered early-phase clinical trials^51–53^.

Here, we use zebrafish CRC xenografts to investigate whether CD24 functionally drives the immune-evasive progressor phenotype of SW620 tumors and to interrogate whether this innate immune checkpoint axis is conserved across vertebrate evolution. Building on the observation that CD24 is strongly upregulated in SW620 progressors, we combine CRISPR–mediated loss of function with *in vivo* clearance assays and live imaging of tumor–myeloid interactions to dissect how CD24 shapes innate immunity to suppress tumor clearance. We then leverage glycocalyx profiling and transcriptomic analysis to explore whether zebrafish macrophage recognition of human CD24 is mediated through conserved sialic acid–binding Siglec–like receptors. Finally, we make use of TCGA CRC cohorts to examine how CD24 expression relates to disease prognosis and outcome. Together, our findings indicate that Siglec-mediated recognition of sialylated glycans likely represents an ancient, evolutionarily conserved mechanism of innate immune regulation, validate CD24 as a therapeutic target in CRC, and establish zebrafish xenografts as a powerful platform to interrogate innate immune-tumor interactions *in vivo*.

## Results

### Transcriptomic analysis identifies CD24 in CRC progressor xenografts

Using the zebrafish xenograft model^11^, we previously established that the SW480 cell line, derived from a CRC primary tumor, presents a regressor behavior, whereas the SW620 cell line, isolated from a lymph node metastasis of the same patient^19,20^, displays a progressor phenotype^21^. This phenotype is defined by engraftment capacity, described as the frequency of xenografts that present a persistent tumor at the end of the assay, at 4 days post-injection (dpi). Importantly, the observed persistence versus clearance phenotype was shown to be critically dependent on the modulation of the zebrafish innate immune population, namely macrophages and neutrophils^21^.

To identify determinants of innate immune suppression and evasion in CRC, we performed a re-analysis of bulk RNA sequencing (RNAseq) data^21^ from SW480 regressor and SW620 progressor tumors isolated from zebrafish xenografts at 2 dpi and compared their transcriptional profiles (**Figure 1A**, **Supplementary Data 1**). Quality control metrics confirmed high-quality sequencing data across all samples (**Supplementary Figure 1A**) and principal component analysis (PCA) of human gene expression revealed clear separation between SW480 and SW620 tumor samples, indicating distinct transcriptional programs (**Supplementary Figure 1B**).

**Figure 1.**
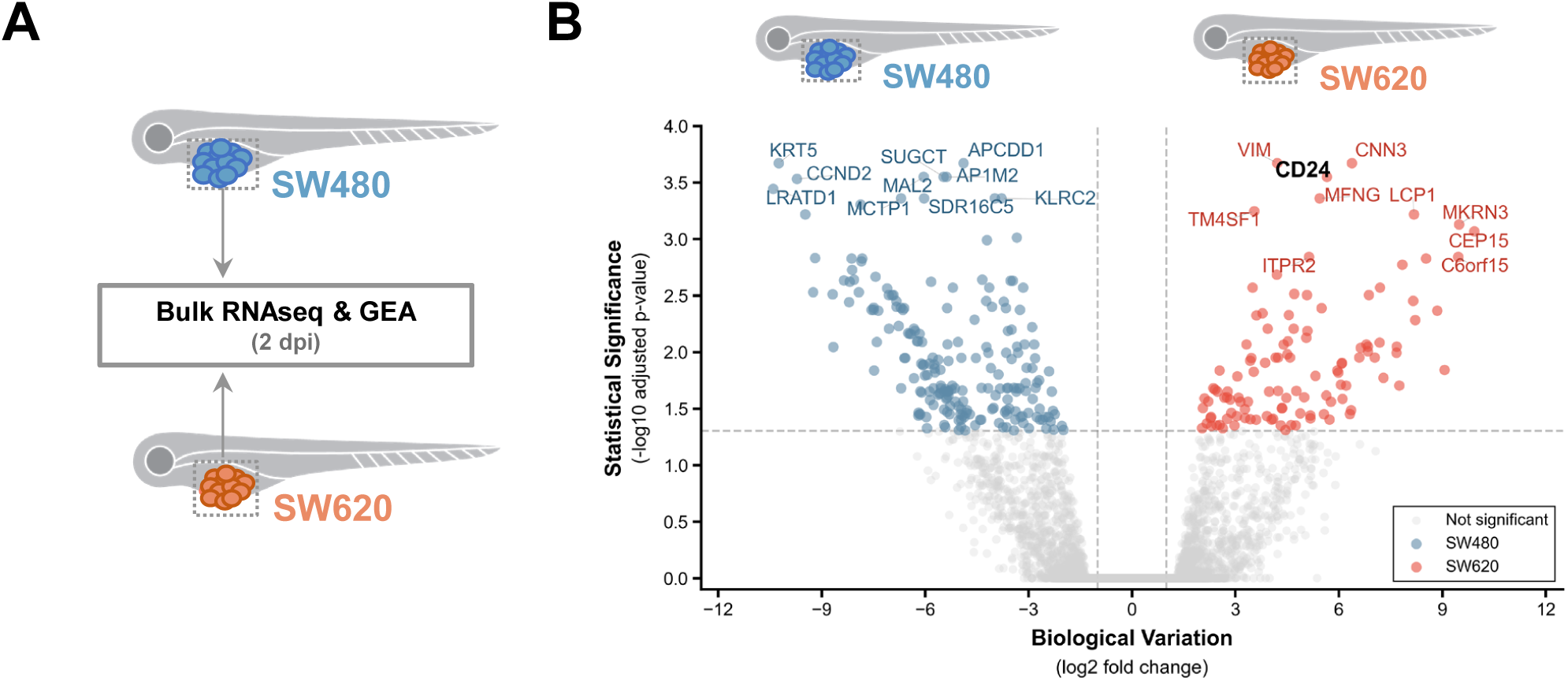
CD24 is overexpressed in progressor CRC tumors. **(A)** Schematic representation of bulk RNA sequencing of SW480 and SW620 tumors dissected from zebrafish xenografts at 2 days post-injection (dpi). GEA - gene expression analysis. **(B)** Volcano plot of differentially expressed human genes between SW480 (blue, 208 DEGs) and SW620 (red, 114 DEGs) tumors. Significantly differentially expressed genes (FDR < 0.05; |log2 fold-change| > 1) are highlighted and CD24 is indicated. Data correspond to 3 and 4 replicates from SW480 and SW620 xenografts, respectively.

Differential gene expression analysis identified 322 significantly differentially expressed human genes between regressor and progressor tumors, with 208 genes enriched in SW480 and 114 genes enriched in SW620 (**Figure 1B**, **Supplementary Figure 2A**). Overall, the transcriptional signature of SW620 progressors was consistent with mesenchymal transition and immune evasion, including upregulation of mesenchymal markers (*VIM*), extracellular matrix remodeling genes (*TNC*), and cell migration-associated factors (*LCP1*). SW620 progressors also showed elevated expression of glycosyltransferases (*GALNT5*, *MFNG*), consistent with the well-established role of aberrant glycosylation in immune evasion^40^. In contrast, SW480 regressors displayed a transcriptional program indicative of epithelial differentiation (*KRT5*, *CLDN7*, *MAL2*) and immunogenicity (NKG2 family receptors, *IL32*, *HLA-DQB1*), consistent with their susceptibility to host-mediated clearance. Gene set enrichment analysis (GSEA) revealed the expected transcriptional polarities of these cell lines (**Supplementary Figure 2B**) and recapitulated our prior work^21^.

To examine host responses to these divergent tumor phenotypes, we next analyzed the zebrafish transcriptome of the samples. GSEA of ranked zebrafish genes (**Supplementary Figure 3**) revealed that SW480 tumors elicited robust activation of innate immune pathways, including the cytosolic DNA-sensing pathway, Toll-like receptor signaling, NOD-like receptor signaling, and myeloid leukocyte differentiation. Accordingly, regressor tumors were also associated with elevated inflammatory metabolism, including electron transport chain and oxidative phosphorylation signatures. In striking contrast, SW620 progressor tumors were associated with enrichment of metabolic insulation programs (nuclear receptors in lipid metabolism, PPAR signaling, sphingolipid metabolism), cell adhesion pathways (neuroactive ligand-receptor interaction, cell adhesion molecules), and developmental plasticity programs (BMP signaling, cell migration regulation).

Among the top upregulated genes in progressors, *CD24* emerged as one of the most significantly upregulated transcripts (log₂FC = 5.65, FDR = 2.82×10⁻⁴, **Figure 1B**, **Supplementary Figure 2C**). CD24 has recently been identified as a novel “don’t eat me” signal that inhibits phagocytosis^28^, making it a compelling candidate underlying the immune-evasive phenotype of SW620 tumors. Together, these transcriptomic data reveal that regressor tumors express an immune-engaged transcriptional program that elicits robust host inflammatory responses, while progressor tumors display features of mesenchymal transition and create an immunologically quiescent microenvironment. These findings also identify CD24 as a potential driver of the progressor phenotype of SW620 tumors and provide the rationale for the functional studies that follow.

### CD24 protects human CRC cells from clearance in zebrafish xenografts

To directly assess the innate immune suppressive role of CD24 in CRC, as well as the conservation of the CD24-Siglec-10 axis in zebrafish, we generated and validated single-cell-derived *CD24* knockout (KO) SW620 clones via CRISPR/Cas9 technology (**Supplementary Figure 4**, **Supplementary Figure 5**). Importantly, CD24 loss did not alter cell viability or proliferation *in vitro* (**Supplementary Figure 6**), indicating that observed phenotypic differences in the xenograft model are not a result of intrinsic growth effects.

To test whether CD24 contributes to the progressor phenotype, we generated SW620 mock and CD24 KO zebrafish xenografts (**Figure 2A**), as previously described^54^. At 1 dpi, all xenografts were screened to select for the successfully injected, which were then monitored until 4 dpi. Tumor clearance was quantified as the ratio between the number of zebrafish xenografts whose tumors disappeared and the total number of xenografts (**Figure 2B**). Loss of CD24 dramatically increased tumor clearance compared to mock controls (**Figure 2C**, ∼70-80% vs ∼20% clearance, 4-fold increase, *p*<0.001). Time-course analysis revealed that the clearance of CD24-deficient tumors begins at a very early onset, with about 50% eliminated by 2 dpi, and continues progressively, whereas mock tumors largely persist over time (**Figure 2D**). These findings demonstrate that CD24 expression protects human CRC cells from clearance in the zebrafish xenograft model.

**Figure 2.**
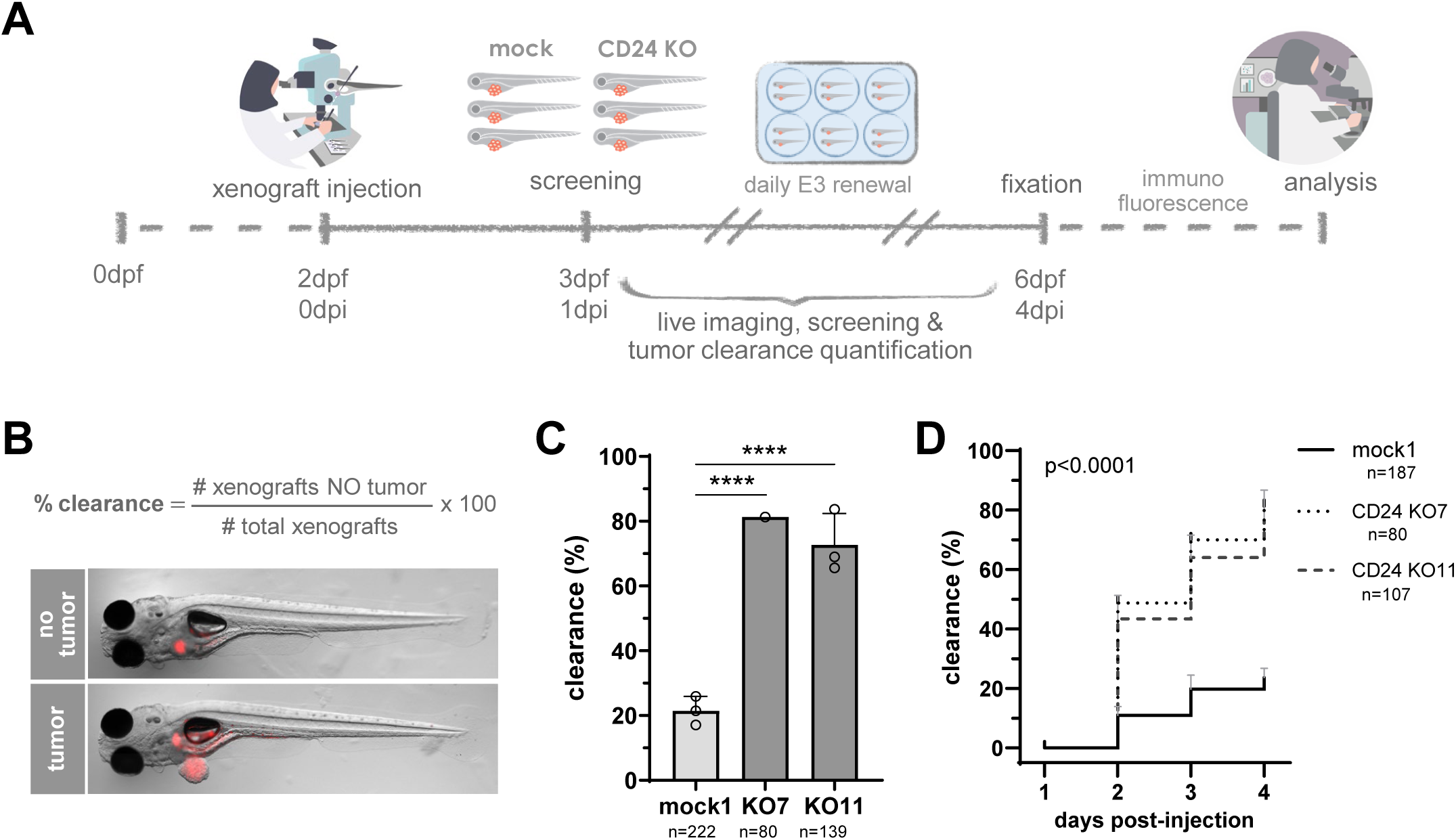
CD24 protects SW620 tumor cells from clearance in the zebrafish xenograft model. **(A)** Schematic representation of the zebrafish xenograft assay methodology. SW620 mock and CD24 KO tumor cells were labeled and injected into the perivitelline space (PVS) of 2 days post-fertilization (dpf) zebrafish larvae. Tumor clearance was screened and quantified from 1 to 4 days post-injection (dpi), when xenografts were fixed for immunofluorescence and further analysis. **(B)** Clearance is calculated as the ratio between the number of zebrafish xenografts whose tumors were cleared and the total number of xenografts at a given time point. Representative brightfield images of xenografts with and without a tumor (in red) at 4 dpi. **(C)** Clearance percentage of SW620 mock, CD24 KO clone 7, and CD24 KO clone 11 xenografts at 4 dpi. The graph shows mean ± S.D. and each dot represents one independent experiment. The total number of xenografts analyzed per condition (n) is depicted. Data were analyzed using Fisher’s exact test (****P<0.0001). **(D)** Tumor clearance evolution of each condition is depicted per day post-injection. The graph shows mean ± S.D. and the total number of xenografts analyzed per condition (n) is depicted. Data corresponds to 2 independent experiments for mock1 and CD24 KO11 and 1 experiment for CD24 KO7. Data was analyzed using the Mantel-Cox test (****P<0.0001).

### CD24 suppresses zebrafish innate immune response

Transgenic lines with fluorescently labeled myeloid cells revealed an increased abundance of both neutrophils and macrophages on CD24 KO tumors relative to mock controls, at both 2 and 4 dpi (**Figure 3A-D**). Furthermore, the proportion of TNFα-expressing macrophages infiltrating CD24-deficient tumors displayed a consistent increase across time, in contrast to mock tumors (**Figure 3B,E,F**; **Supplementary Figure 7**). These results are consistent with a more pro-inflammatory, anti-tumor immune environment in CD24 KO xenografts.

**Figure 3.**
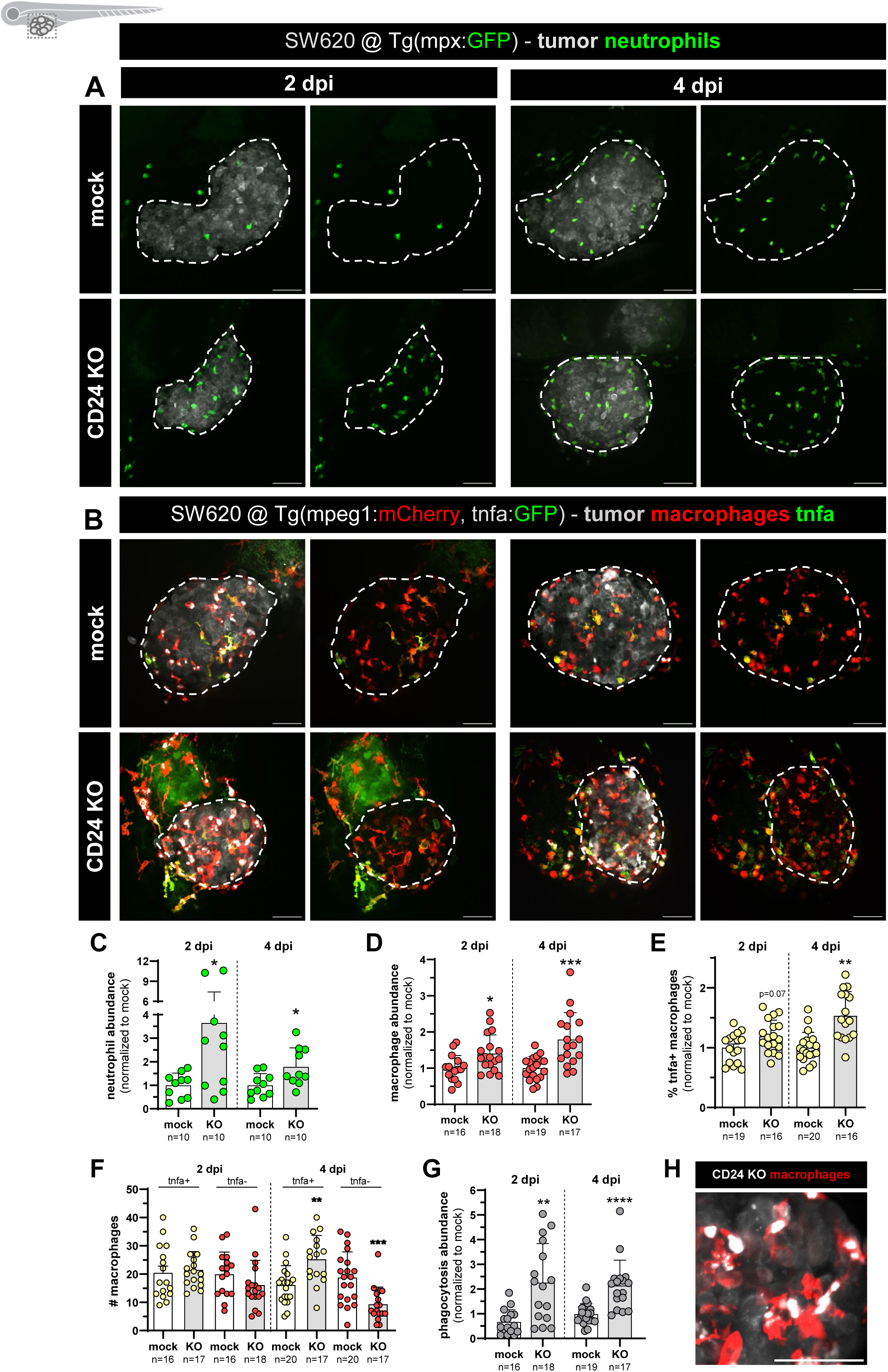
CD24 decreases macrophage and neutrophil abundance and protects tumor cells from phagocytosis. (A,B) Representative confocal images of neutrophils (A, green) and macrophages (B, red) in SW620 mock 1 and CD24 KO11 tumors (in white, outlined by dashed lines), at 2 and 4 dpi. In (B), Tnfa-expressing cells are depicted in green. All images are anterior to left, posterior to right, dorsal up, and ventral down. Scale bars: 50 μm. **(C,D)** Quantification of neutrophil (C) and macrophage (D) abundance at 2 and 4 dpi, calculated as the ratio between the total number of innate cells and the tumor area of each xenograft, normalized to the mock condition. **(E)** Quantification of the percentage of Tnfa-expressing macrophages, normalized to the mock condition. **(F)** Quantification of the total number of Tnfa-expressing macrophages and Tnfa-negative macrophages at 2 and 4 dpi. **(G)** Quantification of phagocytosis abundance at 2 and 4 dpi, calculated as the ratio between the total number of phagocytoses and the tumor area of each xenograft, normalized to the mock condition. **(H)** Representative confocal image of phagocytic macrophages in a SW620 CD24 KO11 tumor. Scale bar: 50 μm. All graphs show mean ± S.D. Data are from 2 independent experiments, each dot represents one zebrafish xenograft, and the total number analyzed (n) is depicted. According to data normality, the Welch’s t test or the Mann-Whitney U test was used to compare mock to CD24 (*P ≤ 0.05, **P ≤ 0.01, ***P ≤ 0.001, and ****P ≤ 0.0001).

Since the literature describes CD24 as a phagocytic “don’t eat me” signal^28^, we investigated whether zebrafish macrophages are capable of recognizing human CD24. To this end, we quantified macrophage engulfment events, which revealed a significant increase in phagocytosis of CD24 KO cells when compared to mock (**Figure 3G,H**), supporting the idea of cross-species conservation of human CD24 working as a functional ligand of a zebrafish macrophage-expressed receptor. Altogether, these findings clearly show that human CD24 suppresses the zebrafish innate immune response by modulating tumor-associated myeloid cell abundance, as well as macrophage inflammatory polarization and phagocytic activity.

To determine if CD24 suppression of tumor clearance is macrophage-mediated, we pharmacologically selectively depleted TME macrophages using clodronate liposomes (L-Clodro). Remarkably, macrophage ablation completely abolished the clearance difference between mock and CD24 KO tumors (**Figure 4**), substantiating this myeloid population as the dominant cellular mediators of CD24-driven immune evasion. Thus, loss of CD24 markedly enhances the phagocytic clearance of human tumor cells by zebrafish macrophages.

**Figure 4.**
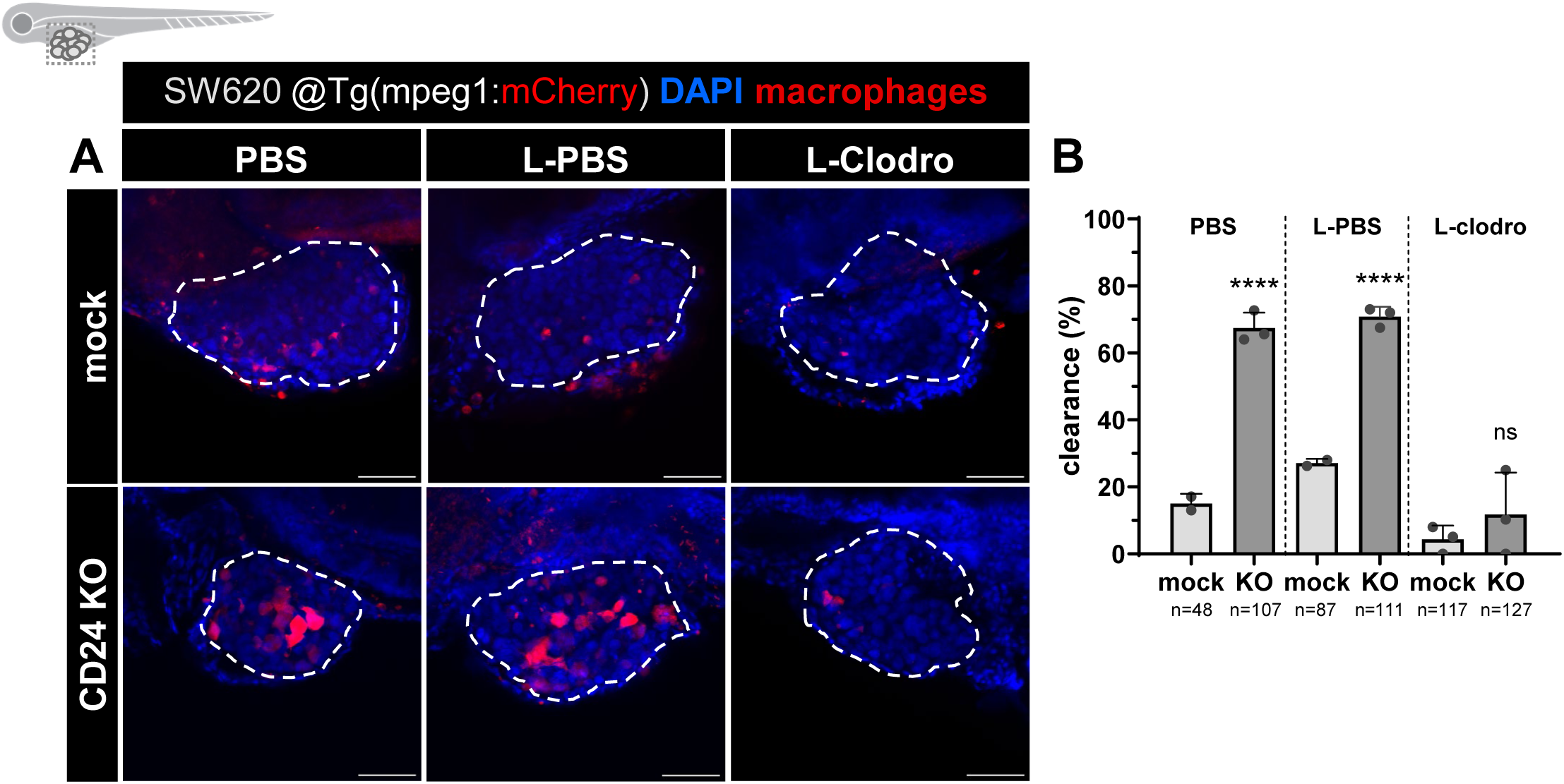
CD24-protection of tumor clearance is macrophage-dependent. **(A)** Representative confocal images of macrophages (in red) in SW620 mock 1 and CD24 KO11 at 4 dpi, in PBS, L-PBS, and L-Clodro conditions. Macrophages were pharmacologically depleted by resuspending tumor cells in clodronate-encapsulated liposomes (L-Clodro) prior to injection. L-PBS-encapsulated liposomes (L-PBS) served as the vehicle control. Tumors are outlined in white, macrophages are labelled in red [Tg(mpeg1:mCherry-F)], and nuclei are stained with DAPI in blue. All images are anterior to left, posterior to right, dorsal up, and ventral down. Scale bars: 50 μm. **(B)** Clearance percentage of SW620 mock and CD24 KO at 4 dpi, in each condition. The graph shows mean ± S.D. and each dot represents one independent experiment. The total number of xenografts analyzed per condition (n) is depicted. Data was analyzed using Fisher’s exact test (ns > 0.05, ****P<0.0001).

### CD24 shapes myeloid motility and effector dynamics in the TME

To gain deeper insight into the dynamics of tumor-immune interactions, we performed live confocal imaging of SW620 mock and CD24 KO xenografts generated in myeloid cell transgenic zebrafish lines [Tg(mpx:GFP), Tg(mpeg1:mCherry), Tg(csf1ra:GFP)] (**Figure 5A**; **Movie 1**, **Movie 2**). Tumors were imaged for approximately 16 h between 1 and 2 dpi, the time window when clearance of CD24 KO cells was most pronounced (Figure 2D). Macrophage and neutrophil tracking (**Figure 5B**, **Movie 3** -**Movie 6**) revealed that CD24 loss markedly altered myeloid cell behavior in the TME, both functionally and in terms of motility. In CD24 KO xenografts, macrophages displayed significantly increased net displacement, higher directionality, and a larger radius of gyration compared to mock controls (**Figure 5C-E**), indicating more extensive and directed exploration of the tumor. Consistently, upon CD24 loss, macrophages spent significantly more time within the tumor, showing reduced mean distance to its center, and were depleted from far–field regions, whereas mock macrophages remained more dispersed and peritumoral (**Figure 5F-H**). Neutrophils showed a reduction in mean velocity and in the variability of their distance to the tumor center (**Figure 5I,J**), indicating a more spatially constrained trajectory. No significant changes were observed in other neutrophil motility or tumor interaction metrics (**Supplementary Figure 8**), underscoring that CD24–dependent reprogramming is largely macrophage–specific at the level of cell behavior.

**Figure 5.**
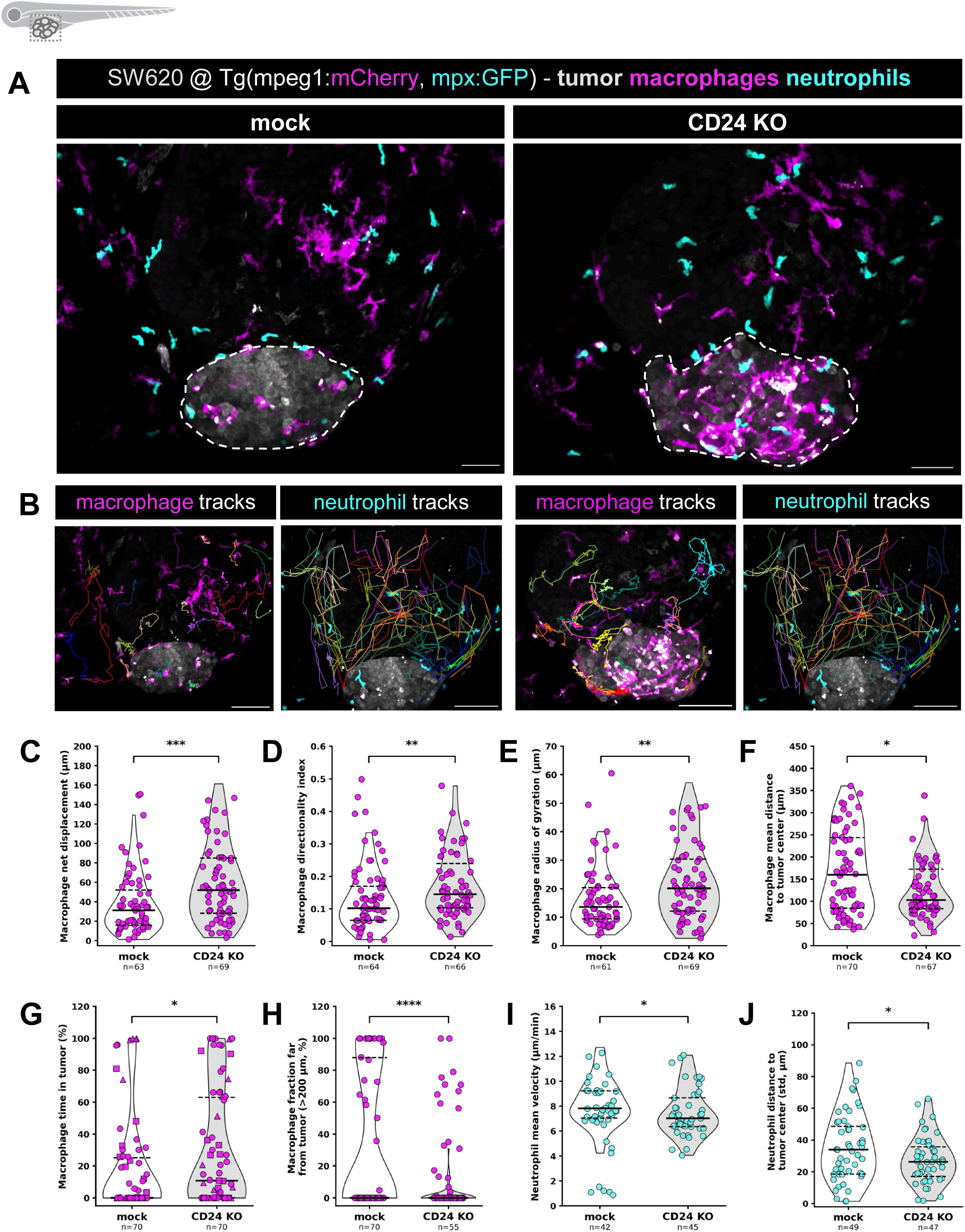
CD24 loss modulates myeloid cell kinetics and stimulates macrophage motility and tumor-directed migration. **(A)** Representative maximum-intensity projections of overnight live-imaging time-lapses of mock and CD24 KO tumors, showing tumor cells (gray), macrophages (magenta), and neutrophils (cyan). SW620 xenografts were generated in Tg(mpeg1:mCherry-F, mpx:GFP-F) larvae and imaged overnight from 1 to 2 dpi. Dashed lines outline the tumor mass. Scale bars: 50 µm. **(B)** Representation of macrophage and neutrophil tracks, in different colored lines, in tumors from each condition. Tracking was performed using the MTrackJ plugin in Fiji. **(C-J)** Quantification of myeloid motility and positioning metrics: macrophage net displacement (C), macrophage directionality index (D), macrophage radius of gyration (E), macrophage mean distance to the tumor center (F), percentage of macrophage time in the tumor (G), percentage of macrophage time spent far from the tumor center (>200 µm, H), neutrophil mean velocity (I), and standard deviation of each neutrophil’s distance to the tumor center over its track (J). Each dot represents a single cell track, violins depict the distribution with median and interquartile range overlaid, and the total number of tracks analyzed (n) is indicated. Data are pooled from 6 xenografts per condition across 3 independent experiments. Depending on data normality, Welch’s t test or Mann-Whitney U test were used to compare mock and CD24 KO conditions (*P ≤ 0.05, **P ≤ 0.01, ***P ≤ 0.001, ****P ≤ 0.0001).

In addition to changes in motility, CD24 loss also promoted macrophage effector functions and myeloid–myeloid interactions in the TME (**Figure 6A**). Macrophages in CD24 KO tumors showed increased fusion activity and tumor cell phagocytosis (**Figure 6B, C**; **Movie 7**, **Movie 8**), supporting an enhanced anti-tumor response. There was also a consistent trend towards higher frequency of macrophage–macrophage and macrophage–neutrophil touches upon CD24 loss (**Figure 6D, E**; **Movie 9**, **Movie 10**), consistent with a more interactive and coordinated myeloid compartment. Together, these data show that CD24 expression on tumor cells constrains macrophages from adopting a highly motile, tumor–directed, and functionally engaged state, thereby contributing to immune evasion.

**Figure 6.**
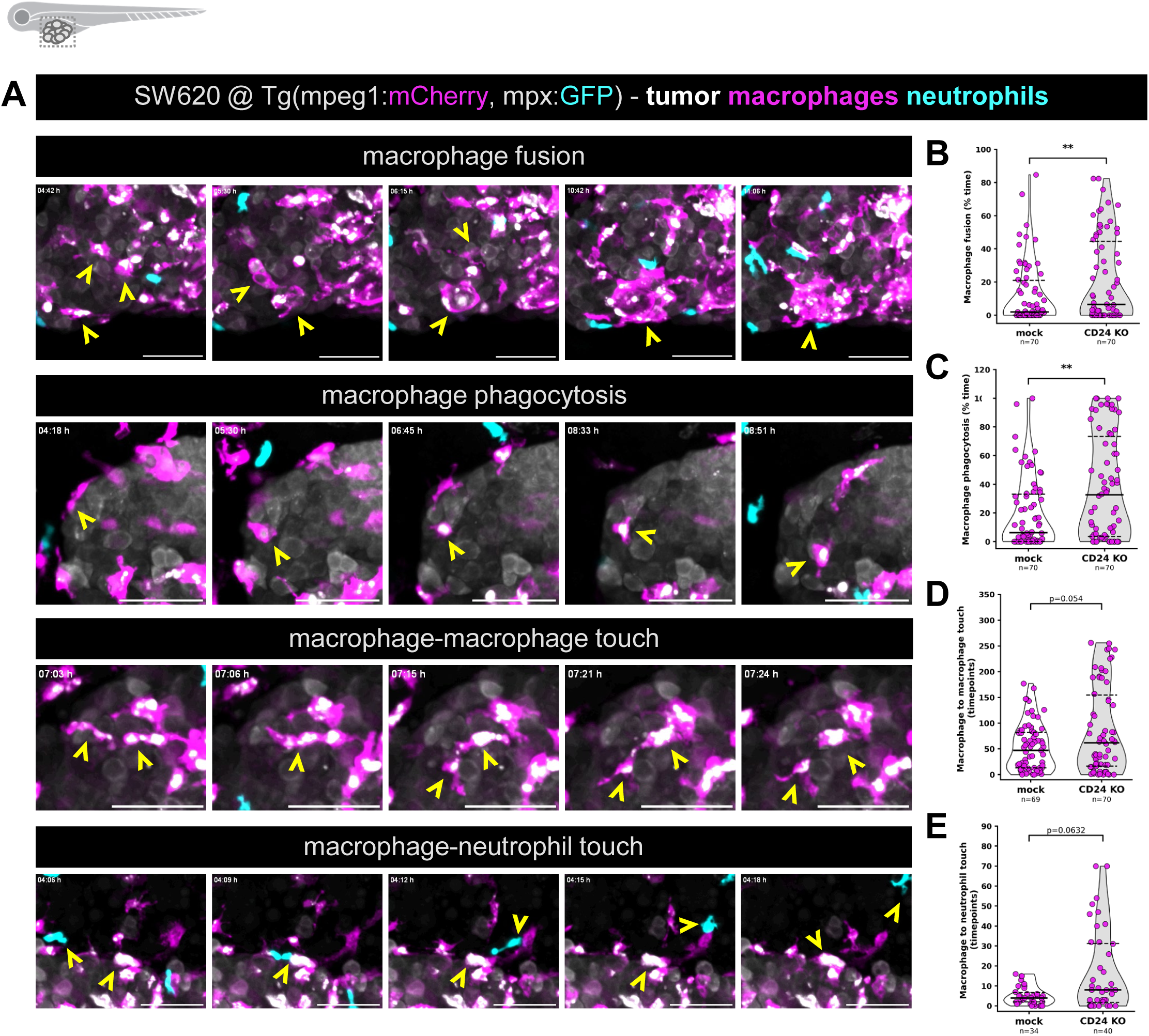
CD24 loss promotes macrophage fusion and phagocytosis events and enhances myeloid–myeloid interactions. **(A)** Representative still images from live-imaging time-lapses illustrating distinct myeloid interaction events in SW620 tumors. Xenografts were generated in Tg(mpeg1:mCherry-F, mpx:GFP-F) larvae and imaged overnight from 1 to 2 dpi. Images show tumor cells (gray), macrophages (magenta), and neutrophils (cyan). Yellow arrowheads highlight each event. Scale bars: 50 µm. **(B-E)** Quantification of macrophage functional and interaction metrics: percentage of time macrophages spend in fusion (B), percentage of time macrophages spend phagocytosing tumor cells (C), number of timepoints macrophages spend in contact with other macrophages (D), and number of timepoints macrophages spend in contact with neutrophils (E). Each dot represents a single cell track, violins depict the distribution with median and interquartile range overlaid, and the total number of tracks analyzed (n) is indicated. Data are pooled from 6 xenografts per condition across 3 independent experiments. Depending on data normality, Welch’s t test or Mann–Whitney U test were used to compare mock and CD24 KO conditions (**P ≤ 0.01).

### CD24 loss remodels the glycocalyx of SW620 cells

Changes in cellular glycosylation are considered a hallmark of cancer and can significantly impact tumor progression and immune evasion^39^. CD24 is a very small but heavily glycosylated protein, composed of only 32 amino acids, whose binding to inhibitory Siglec-10 is dependent on its sialylated glycan coating. Therefore, we sought to characterize how the loss of CD24 alters the glycocalyx composition of SW620 cells (**Figure 7, Supplementary Figure 9**). Flow cytometry analysis with lectins revealed that CD24 KO resulted in elevated cell surface expression of both α-2,6 and α-2,3-linked sialic acids (SNA and MAL-II staining; **Figure 7A, B**). In accordance with this overall increase in sialylated epitopes, binding of recombinant Siglec-3 and Siglec-15 was significantly elevated when compared to mock control cells (**Supplementary Figure 9A, B**). Yet, notably, Siglec-10 engagement was significantly reduced (**Figure 7C**). Interestingly, CD24 KO cells also exhibited increased exposure of high-mannose *N*-glycans (ConA and GNA staining; **Figure 7D, E**), which have been implicated as pro–phagocytic ‘eat me’ signals^55^.

**Figure 7.**
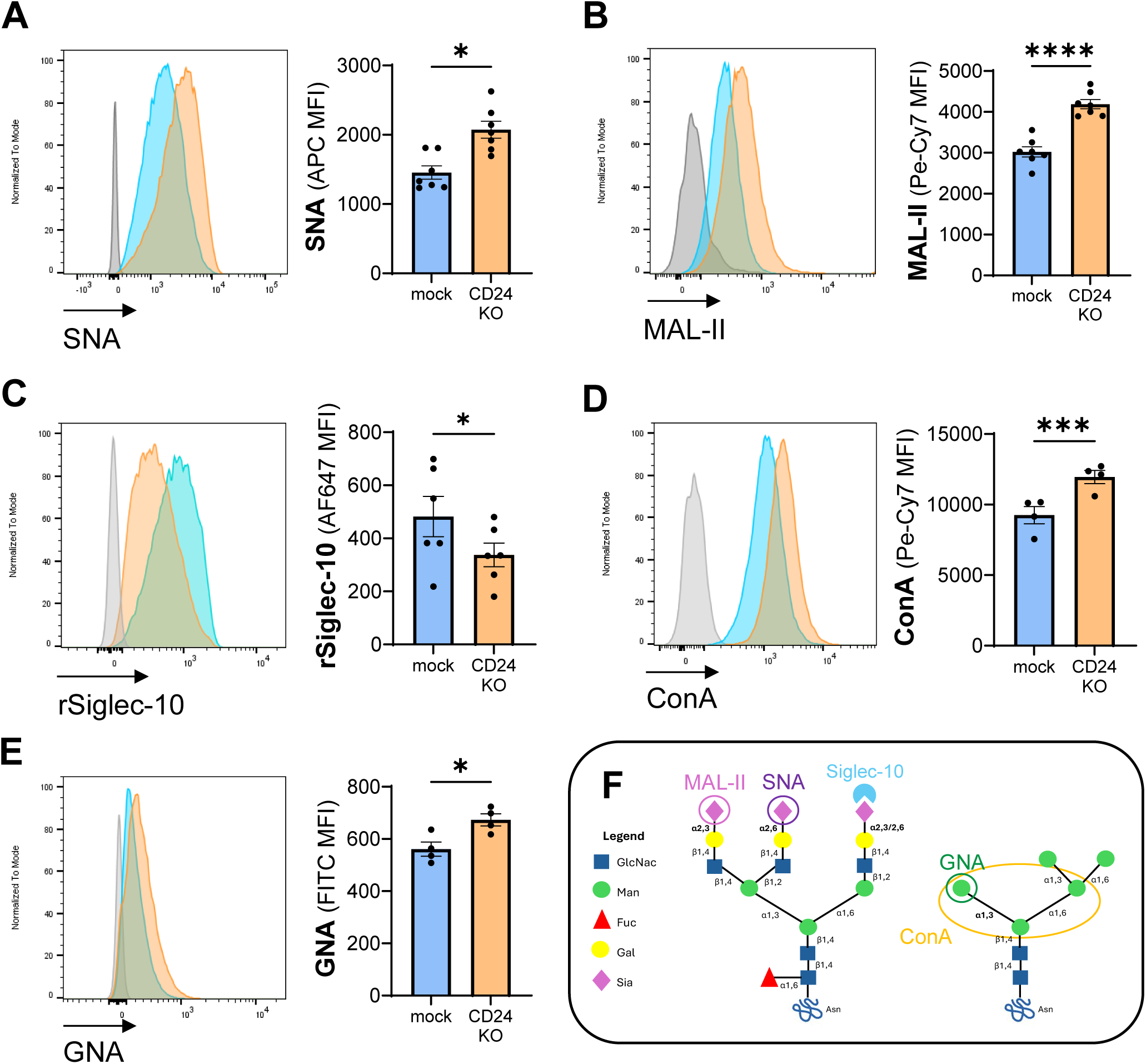
CD24 loss alters the glycocalyx of SW620 cells. Lectin staining histograms and median fluorescence intensity (MFI) of α-2,6-linked sialic acids (**A**, via SNA), α-2,3-linked sialic acids (**B**, via MAL-II), α-D-mannose and α-D-glucose residues (**D**, via ConA), α-1,3-linked mannose (**E**, via GNA) on SW620 mock and CD24 KO cells. Siglec-10 binding was quantified using a recombinant protein (rSiglec-10, **C**). Data are from 2 independent experiments. The graphs show mean ± S.E.M. and each dot represents one replicate. Data normality was analyzed using the Shapiro-Wilk test, and conditions were compared accordingly using a paired t-test or a Wilcoxon test (*P<0.05, **P<0.01, ***P<0.001, ****P<0.0001). **(F)** Schematic representation of the glycan epitopes recognized by each lectin and Siglec-10, shown on representative complex-type sialylated and high-mannose-type N-glycan structures.

Additional glycomic profiling of branching and fucosylation patterns in CD24 KO cells showed a significant increase in polylactosamine structures (LEL staining) and α-1,6-linked fucosylation (AAL staining), alongside a significant decrease in β-1,6-branched *N*-glycans (L-PHA staining), while bisecting-GlcNAc *N*-glycan (E-PHA staining) levels remained unchanged (**Supplementary Figure 9C-F**). Collectively, these results indicate that CD24 deletion triggers a dynamic glycocalyx remodeling, characterized by selective impairment of Siglec-10 engagement and increased exposure of high-mannose pro-phagocytic epitopes.

### Identification of the CD24-like candidate genes in zebrafish

CD24 has been identified in Amniotes, including placental mammals, birds, and non-avian reptiles, but the literature is limited regarding its presence in remaining clades (**Figure 8A**)^56^. To determine whether zebrafish possesses a CD24 ortholog, we sought to study its evolutionary origin. In a first assessment, BLASTp and tBLASTn searches of the human CD24 protein sequence (UniProt ref P25063) against *Danio rerio* proteomic, transcriptomic, and genomic reference databases yielded no clear direct ortholog (data not shown), possibly reflecting the challenge of identifying short protein sequences (∼32 aa) alongside possible coverage gaps in zebrafish genome assemblies. Therefore, we undertook an alternative approach and instead performed a synteny analysis of the genomic region spanning the annotated CD24 gene in selected species (human, mouse and chicken), using the flanking genes as anchors for the synteny search in representative species of Gnathostomes (**Figure 8B**). We located the genomic region of interest in the intergenic region between QRSL1 and either MTRES1 or iars1, depending on the species, but could not locate any annotated gene that resembled CD24. However, in RNASeq data of all species analyzed, we observed a consistent signature of 2-exon short RNA sequences, most annotated as long non-coding RNAs (lncRNAs) and long intergenic non-coding RNAs (lincRNAs), in a very similar configuration and length to *bona fide* CD24 genes, including si:211-214p16.1 and si:211-214p16.2 in zebrafish. Since lincRNAs are known to encode micropeptides and some lncRNA annotations are temporarily categorized as such until fully validated as genes, we hypothesized these loci could be candidate CD24-like genes. To address that, we extracted the corresponding genomic and exonic sequences to generate the respective pseudo-cDNA and amino acid sequences. Although CD24 is known as a fast-evolving protein, we could nevertheless still observe amino acid conservation in particular residues of several candidate sequences and *bona fide* CD24, including the putative proteins encoded by the 2 zebrafish candidate loci (**Figure 8C**). Since in the case of C.milli the intron start and end positions corresponded to CA-TC, which are unlikely to work as donor and acceptor sequences, we manually curated the exon border, shifting it by 4 nucleotides downstream, and hypothesize the curated model to contain GG-AG donor and acceptor sequences, where non-canonical GG could originate from a sequencing error of a canonical GT donor sequence. The resulting reconstructed protein resembled the remaining CD24 candidate proteins.

**Figure 8.**
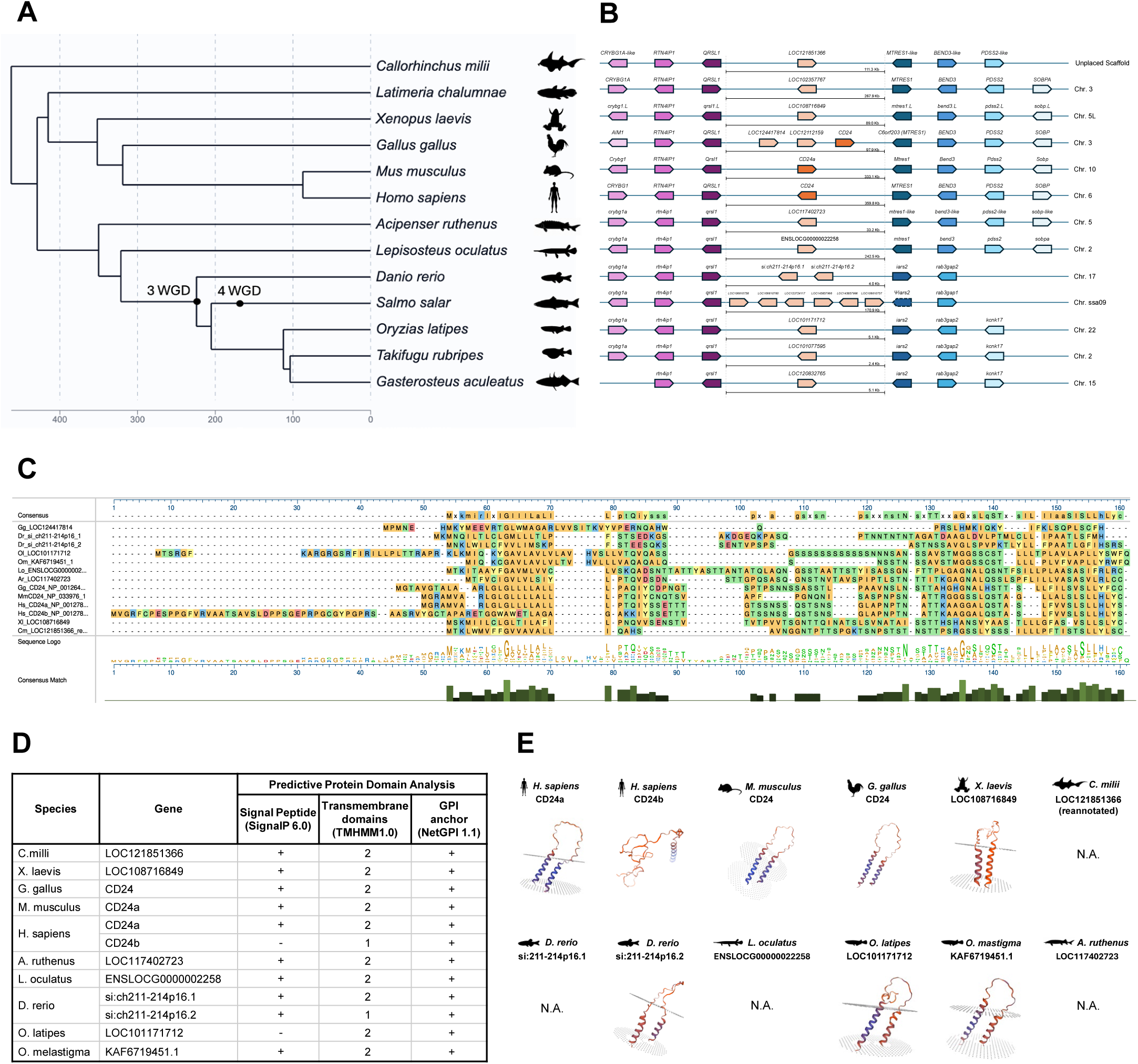
Comparative genomics and molecular evolution of CD24 and candidate CD24-like genes across Vertebrates. **(A)** Phylogenetic relationships among the vertebrate species included in the analysis; approximate divergence times obtained in Timetree are shown in millions of years before present, and the teleost-specific third whole-genome duplication (3 WGD) and salmonid-specific fourth whole-genome duplication (4 WGD) events are indicated. **(B)** Comparative synteny of the genomic region containing *CD24* in amniotes and the corresponding syntenic interval in other vertebrates; conserved flanking genes are indicated by matching colors, with arrowheads denoting transcriptional orientation. Annotated *CD24* genes are shown in orange, whereas genes located within the corresponding candidate interval are shown in light orange. The size of the interval between the conserved flanking regions is indicated for each species. Genomic intervals are represented schematically and are not drawn to scale. Chromosome or scaffold locations are indicated on the right. **(C)** Multiple amino acid sequence alignment of annotated CD24 proteins and selected proteins encoded within the candidate syntenic regions. The alignment was performed using the EMBL-EBI Multiple Sequence Alignment tool (Clustal Omega) and visualized in MegAlign Pro. Amino acids are color-coded by chemistry, and residue consensus and conservation are also displayed. **(D)** Predicted protein features of CD24 and candidate proteins. N-terminal signal peptides, transmembrane domains, and GPI-anchor signals were assessed using SignalP, TMHMM, and GPI-Net, respectively. “+” and “−” indicate predicted presence or absence of the corresponding feature. For transmembrane domains, the predicted number of domains is shown. **(E)** Predicted three-dimensional structures and membrane positioning of selected annotated CD24 and candidate proteins generated using SWISS-MODEL. N.A. indicates sequences for which no structural model was obtained.

**Figure 9.**
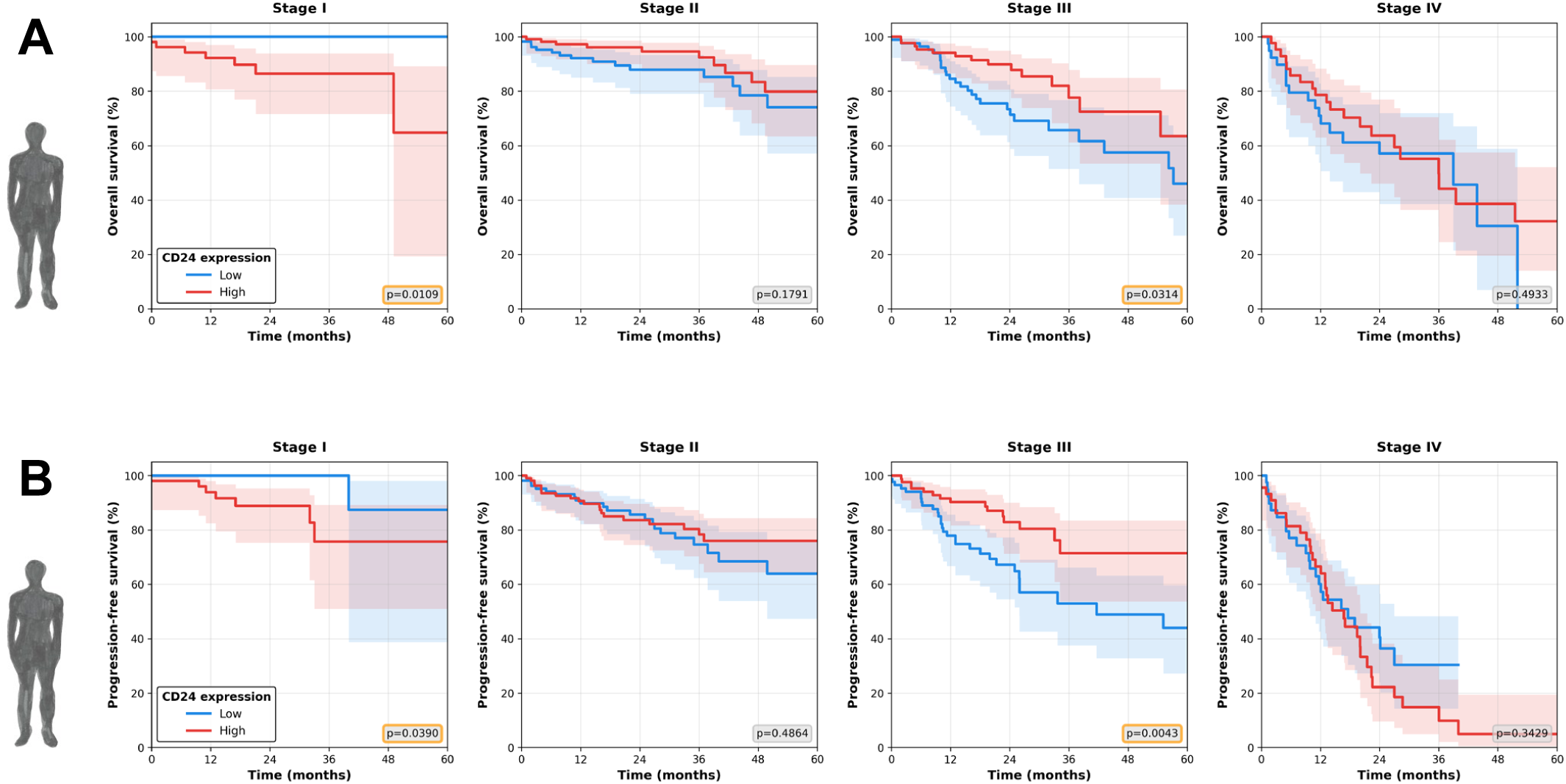
Clinical relevance of CD24 in colorectal cancer. **(A)** Overall survival (OS) and **(B)** Progression-free survival (PFI) Kaplan-Meier curves for TCGA COAD + READ patients (n=601), according to disease staging. Patients were stratified by CD24 expression (high in red, low in blue) based on median split. Data was analyzed using log-rank tests and significant comparisons are highlighted in orange boxes.

Protein motif prediction and protein modeling revealed striking resemblance to CD24 of these candidates, not only by the presence of Signal Peptide, 2 transmembrane domains and GPI-anchor motifs (**Figure 8D**), but most strikingly protein structure. Interestingly, the CD24b isoform of human, although previously thought to be non-functional, is experimentally studies and described to produce a functional phenotype *in vitro*^57^. In zebrafish, only si:211-214p16.2 returned a protein model, and this is possibly due to the lack of templates that resemble CD24 with its idiosyncratic properties and serve as basis for inference of structure. We therefore conclude that, contrary to previously thought, CD24-like sequences are present in all Gnathostomes and zebrafish contains at least one CD24-like candidate gene.

### Identification of a candidate CD24-binding receptor in zebrafish macrophages

As our data clearly indicates recognition of human CD24 by zebrafish macrophages, we next sought to identify the zebrafish receptor responsible for this interaction. In humans, CD24 is recognized by the sialic acid–binding receptor Siglec-10, with Siglec-G representing the corresponding paralog in mice^28^. However, a functional homolog has yet to be identified in zebrafish. Therefore, we performed a BLASTp search of human Siglec-10 against the *Danio rerio* proteome, using both the full-length protein sequence and its Ig-like V-type (IgV) domain, which mediates ligand binding.

Three zebrafish Siglec-like proteins, *mag*, *siglec15l*, and *si:dkey-24p1.7* (**Supplementary Table 8**), were prioritized based on sequence similarity, conservation of the IgV domain, and preservation of the critical arginine residue required for sialic acid recognition, as confirmed by multiple sequence alignment (**Supplementary Figure 10A, B**). All three candidates are predicted to be cell membrane-associated, based on structural domains, and show annotated homology to other members of the mammalian Siglec family, supporting potential functional orthology to Siglec-10 in zebrafish.

Next, we made use of publicly available zebrafish single-cell RNA sequencing data^58^ to determine if these candidates are expressed by macrophages during the developmental period of our xenograft assay (2 to 6 dpf). Within the datasets analyzed, *siglec15l* showed no expression, *mag* showed very low expression, and *si:dkey-24p1.7* showed high expression in macrophages (**Supplementary Figure 10C, D**). Additionally, only *si:dkey-24p1.7* was present in our TME zebrafish RNAseq data (**Supplementary Figure 2D**), making it the strongest candidate for CD24 sensing in our model.

### CD24 expression predicts poor survival specifically in early-stage CRC disease

To assess the clinical relevance of CD24 in human CRC, we analyzed RNAseq data and clinical outcomes from 601 patients in the TCGA COAD and READ cohorts. Within each pathological stage cohort (stages I-IV), patients were stratified by median split of CD24 expression into high and low groups (**Supplementary Table 10**). Survival analysis revealed a stage-dependent relationship between CD24 expression and patient outcomes (**Supplementary Data 4**). In Stage I disease (n=105), high CD24 expression was associated with significantly worse overall survival (OS, *p*=0.011; **Figure 8A**) and progression-free survival (PFI, *p*=0.039, HR=6.74; **Figure 8B**). Intriguingly, this relationship reversed in Stage III disease, where high CD24 expression was associated with both improved OS (*p*=0.031, HR=0.50) and PFI (*p*=0.004, HR=0.41). No significant associations were observed in Stage II or Stage IV disease. This stage-dependent divergence in prognostic value suggests that CD24 function is context-dependent and varies across tumor evolution, which warrants careful interpretation and has important clinical implications. Nevertheless, these results establish CD24 as a stage-dependent prognostic marker in CRC, with high expression predicting poor outcomes in early-stage disease.

## Discussion

The innate immune system represents the first line of defense against malignancies, yet how tumors evade myeloid-mediated clearance is still being actively uncovered. We first identified CD24 as a candidate mediator of innate immune evasion while developing CRC zebrafish xenografts. A transcriptomic comparison of SW480-regressor vs SW620-progressor xenografts revealed CD24 as one of the most highly upregulated genes in immune-evasive progressors. This was notable as CD24 has been recently identified as an innate immune checkpoint ligand that suppresses macrophage phagocytosis, with antibody blockade demonstrating therapeutic potential in preclinical models of several cancer types^28,59–63^. These convergent findings led us to investigate whether CD24 functionally drives the immune-evasive progressor phenotype of our zebrafish CRC model and the possible conservation of this checkpoint mechanism across vertebrate evolution.

To answer these questions, CD24 was genetically inactivated in SW620 cells. CD24 deletion was sufficient to convert the progressor phenotype into an immune-sensitive regressor phenotype. CD24 KO tumors were not only cleared at a ∼4-fold increase compared to mock controls, with only 20-30% of tumors persisting by 4 dpi, but this clearance also showed very early onset, with around half of the tumors no longer detectable by 2 dpi. Critically, pharmacological macrophage depletion completely abolished this clearance phenotype, validating macrophages as the principal mediators of CD24-driven immune escape. These results establish CD24 as a major determinant of immune evasion in this system.

Using transgenic zebrafish lines and confocal imaging to visualize tumor-myeloid cell dynamics *in vivo*, we dissected the cellular mechanisms underlying CD24-mediated immune suppression. Our findings reveal that CD24 shapes macrophage biology across three interconnected levels – activation state, behavior within the TME, and effector output – collectively preventing the productive myeloid response required for tumor clearance. At the level of activation, CD24 KO tumors harbored a higher proportion of TNFα-expressing macrophages. This is particularly consequential considering the well-established role of TNFα as a hallmark of pro-inflammatory macrophage polarization and as a key mediator of tumor cell killing through direct cytotoxicity and induction of apoptotic pathways^64–66^. CD24-mediated suppression of this anti-tumoral activation is accompanied by profound changes in macrophage kinetics, as revealed by live imaging. In the absence of CD24, macrophages became markedly more motile and tumor-directed, spending more time within the tumor and being depleted from peritumoral regions. This transition from a dispersed, patrol-like state to one of committed intratumoral engagement suggests that CD24 actively confines macrophages to a restrained mode that limits productive tumor contact. At the effector level, and consistently with its role as a “don’t eat me” signal, CD24-deficient tumors showed markedly increased macrophage phagocytosis, similarly to what has been seen in other studies^28,67,68^. Strikingly, CD24 loss also significantly increased macrophage fusion activity. Fusing phagocytic macrophages morphologically resemble multinucleated giant cells, a macrophage-specific and highly evolutionarily conserved phenomenon whose formation has been proposed to confer enhanced phagocytic and antimicrobial capacity^66,69^. The co-occurrence of these fusion events with increased phagocytosis and more frequent macrophage–macrophage and macrophage– neutrophil contacts points to a more coordinated and effector-biased myeloid compartment upon CD24 loss. Together, these findings demonstrate how CD24 orchestrates a coordinated and multilayered suppression of innate immunity, dampening macrophage polarization, tumor engagement, and effector capacity, to prevent the coordinated myeloid response that drives efficient tumor clearance.

Perhaps our most striking finding is the observation that zebrafish macrophages recognize and respond to human CD24, despite considerable evolutionary divergence. Furthermore, we identified at least one candidate CD24-like ortholog in zebrafish. The fact that its presence has been hidden in plain sight is likely explained by the nature of CD24 itself: a bicistronic gene whose expression could easily be confounded with lncRNAs or lincRNAs in RNAseq datasets. Also CD24 produces heavily glycosylated, GPI-anchored, mucin-like protein that evolves rapidly and lacks one-to-one orthologs outside mammals^63,70^. This led us to explore the possibility that the molecular basis of this cross-species conservation lies at the level of signaling logic rather than strict ligand homology^71^. Since the mammalian CD24-Siglec axis binding depends on the sialylated glycan coating on cancer cells^72^, we profiled the glycocalyx of SW620 cells. CD24 deletion triggered a dynamic remodeling of cell surface glycans, including increased overall sialylation that likely reflects compensatory redistribution of these modifications to other glycoproteins upon removal of a major mucin–like acceptor^39,73^. Crucially, despite increased sialylated epitopes, Siglec-10 engagement was markedly reduced in CD24 KO cells, indicating that CD24 does not merely act as a major sialylated ligand among many, but provides a spatial and structural glycan context that particularly favors immune inhibitory signaling^63^. Increased binding of Siglec–3 and Siglec–15 further suggests that CD24 loss results in a remodeling of the cell surface sialylation that is recognized by distinct Siglec receptors, consistent with selective disruption of the Siglec–10 recognition axis rather than a global collapse in inhibitory sialylation^74^. Beyond this, CD24 ablation also led to significantly increased abundance of high-mannose glycans, recognized as immunogenic “eat-me” cues by C–type lectin receptors, such as the mannose receptor (CD206) on macrophages^55,75^. We propose that CD24 functions as a steric “glycan shield” and that its removal unmasks these pro-phagocytic underlying epitopes. This dual event, selective loss of Siglec-10 engagement and exposure of pro-phagocytic mannose ligands, likely synergizes to drive phagocytosis and clearance of CD24 KO cells *in vivo*. These findings allow us to hypothesize that zebrafish macrophages, similarly to their mammalian counterparts, rely on sialic acid-dependent mechanisms for recognition of human CD24, establishing a molecular basis for this cross-species interaction. This supports the idea that inhibitory recognition of sialylated glycans by Siglec-mediated pathways is an evolutionarily conserved strategy of innate immune regulation, a degree of conservation that typically marks mechanisms of fundamental biological importance^76,77^ and that offer more difficulty for tumors to evade through alternative resistance routes^78–80^.

On the other side of this inhibitory axis, we have CD24’s binding partner, Siglec-10, which also lacks a clear ortholog in zebrafish. Siglecs are sialylated glycan-recognizing proteins, primarily expressed on immune cells, which can be divided into two groups according to their homology and evolutionary conservation: conserved Siglecs – which include Siglec-1, CD22, Siglec-4 (MAG), and Siglec-15 – and CD33-related Siglecs, of which Siglec-10 is part^26,81^. This large protein family has undergone rapid evolution and diversification in vertebrates, with dramatic lineage-specific expansions and losses^82,83^. As such, teleost fish possess a smaller Siglec repertoire than mammals, with only clear orthologs identified for the conserved Siglecs^84–86^, likely reflecting an ancestral state with broader ligand specificity per receptor that enables functional compensation through promiscuous ligand recognition. Our bioinformatic analysis identified three candidate zebrafish receptors for CD24 – *mag*, *siglec15l*, and *si:dkey-24p1.7* – that share key structural features with human SIGLEC10, including conservation of the Ig-like V-type domain and the critical sialic acid-binding arginine residue. Single-cell expression profiling revealed that *si:dkey-24p1.7* shows robust expression in zebrafish macrophages during our xenograft window, nominating it as the most likely functional analog for CD24 recognition. Functional validation, e.g., through genetic ablation, is ongoing to validate *si:dkey-24p1.7* as the zebrafish mediator of this interaction.

Our TCGA analysis of 601 CRC patients revealed a striking stage-dependent relationship: high CD24 expression predicted poor survival specifically in Stage I disease, whereas this relationship reversed in Stage III, where high CD24 expression was associated with improved outcomes. This divergence mirrors the inconsistent literature on CD24’s prognostic value in CRC, as some cohorts report poor outcomes connected to high expression^87,88^, while others report the opposite association^89^ or no significant relationship at all, despite a clear link to tumor differentiation^90^. This stage-dependent reversal suggests that CD24 integrates opposing, context-dependent roles in tumor biology that evolve throughout disease progression.

As an innate immune checkpoint ligand, CD24-mediated suppression of macrophage-mediated clearance would be expected to predict a worse outcome. Consistently, CD24 expression has been linked to multiple aspects of CRC progression, including enhanced proliferation^91,92^, angiogenesis^93^, stemness and EMT^94,95^. Accordingly, we propose that in early-stage disease, where tumor burden is low and innate immune surveillance is presumably still a major barrier to persistence, these checkpoint, proliferative, and stemness-associated functions of CD24 predominate, consistent with the poor prognosis we observe in Stage I. However, this contrasts with settings in which CD24 expression instead associates with more differentiated, less aggressive tumor phenotypes/subclones^96,97^, which could potentially explain the outcome reversal observed in stage III. Subcellular compartment adds a further layer of context-dependence on top of these functional roles, with membranous, cytoplasmic, and nuclear CD24 each independently associated with distinct, and sometimes contradictory, prognostic outcomes across cancer types^98–100^.

Given this functional multiplicity, the relationship between CD24 expression and outcome does not converge on a single direction, even within CRC alone, meaning it can hardly carry consistent prognostic meaning alone. Whether CD24 signals immune evasion, proliferative and invasive capacity, stemness, or differentiation likely depends on which of these programs predominates at a given disease stage and context. Further, CD24 levels can also be shaped by factors independent of intrinsic tumor biology, such as therapeutic pressure: CD24-low, CD133⁺ CRC cells have been identified as a chemoresistant, stem-like subpopulation that expands following 5-FU treatment^101^, while conversely, chemo-radiotherapy has been shown to upregulate CD24 on tumor cells and Siglec-9/10 on macrophages in cervical cancer^102^. Collectively, these findings have actionable translational implications for ongoing therapeutic developments, such as anti-CD24 antibodies currently in clinical trials^52,53,103^. This is particularly relevant where CD47- and PD-1-directed therapies underperform: CD24 expression has been linked to poor response to PD-1/L1 blockade^104^ and has outperformed PD-1 as a target in preclinical models^105^, motivating combinatory approaches across distinct checkpoint axes^51,106–108^. Given CD24’s context-dependence, bulk expression alone is an inherently noisy proxy for its immune-evasive contribution, but combined biomarkers, such as CD24 expression paired with disease stage and Siglec-10⁺ TAM infiltration, could instead better isolate this specific function to inform therapeutic decision-making. Together, these considerations position CD24 as a potent, but context-dependent immunoregulatory agent for which therapeutic translation requires biomarker-aligned design^109^.

This study validates zebrafish as a powerful platform for innate tumor immunity research, offering unique advantages: temporal separation of innate and adaptive immunity, high functional conservation to humans, and optical transparency enabling whole-body live imaging at single-cell resolution^16,18^. The possibility of cross-species molecular compatibility of human ligands to zebrafish receptors reveals translational potential for screening human-directed immunotherapies, such as innate checkpoint-blocking antibodies and glycan-targeting strategies. This approach can also leverage the model’s established strengths for high-throughput *in vivo* screening – speed and cost-effectiveness^110–113^ – which has the potential to substantially accelerate drug development. Notably, zebrafish xenografts have demonstrated success as predictive platforms for evaluating drug responses, where treatment efficacy is assessed by quantifying tumor cell apoptosis and/or clearance^66,114–117^, further supporting their translational utility.

In conclusion, our findings establish CD24-mediated immune evasion as an ancient evolutionary mechanism, enacted through sialic acid-dependent glycan recognition. We validate CD24 as a high-priority therapeutic target in CRC and emphasize the relevance of zebrafish xenografts as a platform for dissecting innate immune-tumor interactions and accelerating therapeutic discovery and development.

## Materials and Methods

### Zebrafish husbandry

Zebrafish (*Danio rerio*) were handled and maintained according to the standard protocols of the European Animal Welfare Legislation, Directive 2010/63/EU (European Commission, 2016), and the Champalimaud Foundation Fish Platform. All protocols were approved by the Champalimaud Animal Ethical Committee and Portuguese institutional organizations – Órgão de Bem-Estar e Ética Animal (ORBEA, Animal Welfare and Ethics Body) and Direção Geral de Alimentação e Veterinária (DGAV, Directorate General for Food and Veterinary). Zebrafish between 3 and 18 months of age were reared in 3.5 L tanks filled with E3 medium, at a density of 10 fish/L. The rearing temperature was 28°C and the animals were kept in a light/dark cycle of 14 h/10 h (lights on from 08:00 until 22:00). Zebrafish were fed three times per day, artemia in the mornings, and powder (Sparos 400-600, Techniplast) in the afternoons and evenings.

### Zebrafish transgenic lines

According to the purpose of each experiment, different genetically modified zebrafish lines, in the Tübingen background, were used in this study: Tg(*mpx:GFP*) i114Tg^118^, Tg(*mpeg1:mCherry*) ump2^119^, Tg(*csf1ra:GFP*) sh377Tg^120^, Tg(*mpeg1:mCherry-F; tnfa:GFP-F*) ump2Tg;ump5Tg^121^, Tg(*fli:GFP*) y1Tg^122^.

### Human cancer cell lines and culture

The human colorectal cancer cell lines SW480 and SW620 were obtained from the American Type Culture Collection (ATCC). To guarantee replication conditions, the cell lines were expanded and frozen at low passages in 90% FBS (Sigma-Aldrich) plus 10% DMSO (Sigma-Aldrich) for long-term cell cryopreservation in liquid nitrogen. When thawed, the cells were cultured in Dulbecco’s Modified Eagle Medium (DMEM) High Glucose (Biowest), supplemented with 10% FBS (Sigma-Aldrich) and antibiotics (100 U ml−1 penicillin and 100 μg ml−1 streptomycin, Hyclone), maintained in a humidified atmosphere of 5% CO2 at 37 °C. Splitting was performed using TrypLE (Thermo Fisher Scientific) every ∼3/4 days or when cells reached ∼70-80% confluence, and cells were used for experiments only up to 12 passages after thawing. All cell lines were authenticated by short tandem repeat (STR) profile analysis and tested routinely for mycoplasma contamination by PCR.

### Cell staining

Tumor cells were grown to 70% confluence in T-75 flasks, washed with DPBS 1x (Biowest), and detached enzymatically using TrypLE (Thermo Fisher Scientific). Cell suspensions were collected in 15 ml centrifuge tubes, spun down at 300 g for 4 mins, and resuspended in DPBS 1x. Cells were then stained in 1.5 ml microcentrifuge tubes using lipophilic dyes – Vybrant CM-DiI (4 µl/ml in DPBS 1x) or Deep Red Cell Tracker (1 µl/ml in DPBS 1x, 10 mM stock) (Life Technologies) – for 15 min at 37 °C, while protected from light. Cells were then spun down at 300 g for 5 min and resuspended in complete medium. Viability was assessed by the Trypan Blue exclusion method, and cell number was determined by hemocytometer counting. Cells were resuspended in DPBS 1x to a final concentration of 0.25 × 10^6^ cells/μl.

### Zebrafish xenograft generation

On injection day, 2 dpf zebrafish embryos were separated from unhatched eggs, and Pronase (Roche) 1x was added to the E3 medium of the eggs to boost hatching. During injection, embryos were anesthetized by incubation in Tricaine 1x (1.6 g/L, Sigma-Aldrich) for 5 min, transferred to an agarose plate, and carefully aligned with the help of a hairpin loop. Fluorescently labeled cancer cells were injected using a microinjection needle under a fluorescence scope (Zeiss Axio Zoom V16) with a milli-pulse pressure injector (Applied Scientific Instrumentation, MPPI-3). Approximately 500-1000 cells were injected into the perivitelline space (PVS) of each embryo. After injection, embryos were kept in Tricaine for 10 min to allow for wound closing, followed by transfer to fresh E3 medium and incubation at 34 °C.

At 1-day post-injection (dpi), zebrafish xenografts were screened according to the presence or absence of a tumor, under a fluorescence scope. Xenografts with edema, cells in the yolk sac, and cellular debris were discarded, whereas successfully injected ones were grouped according to their tumor size, classified by comparison with the embryos’ eye size. Daily, from 1 to 4 dpi, the E3 medium was refreshed, and zebrafish xenografts were analyzed to quantify tumor clearance rate as follows:

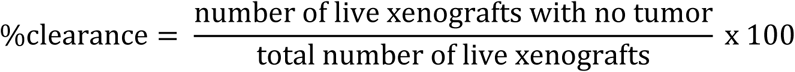

At 4 dpi, xenografts were sacrificed by Tricaine overdose, fixed with 4% Formaldehyde (FA) (Thermo Scientific) overnight at 4 °C, and preserved at −20 °C in 100% methanol.

### Bulk RNA sequencing

#### Sample preparation and sequencing

Raw bulk RNAseq data from previous work from the lab^21^, which had been deposited in NCBI Gene Expression Omnibus (GEO) (accession number <u>GSE163746</u>), was used. Briefly, SW480 and SW620 tumors were dissected (∼30 tumors per sample) from zebrafish xenografts at 2 dpi and pooled from multiple independent experiments. Total RNA was extracted using TRIzol (Thermo Fisher Scientific, #15596026) and further purified with the RNeasy Plus Micro Kit (Qiagen, # 74034). mRNA libraries were prepared using the Smart-seq2 protocol and sequenced on an Illumina NextSeq 500, generating unstranded single-end reads of 76 bp, and averaging ∼38 million reads per sample.

### Data analysis

Quality control was assessed using FastQC^123^ (v0.11.8, **Supplementary Figure 1A**, **Supplementary Data 1**). Quality filtering and transcript-level quantification were performed using Salmon (v1.10.3)^124^ against the human (GRCh38.p14) and zebrafish (GRCz11) reference transcriptomes (Ensembl release 115), separately. Using R, transcript-level abundance estimates were summarized to gene-level counts using tximport^125^, with counts derived from length-scaled TPMs. Human and zebrafish genes were analyzed independently using species-specific transcript-to-gene mappings derived from Ensembl GTF annotations. Differential expression analysis was conducted using the edgeR^126^ and limma^127^ workflow. Lowly expressed genes were filtered using the filterByExpr function, library sizes were normalized using the trimmed mean of M-values (TMM) method, and expression values were transformed using voom^128^ prior to linear modeling. Differential expression was assessed using limma-treat^129^, applying thresholds of absolute log2 fold-change > 1 and false discovery rate (FDR) < 0.05. Gene set enrichment analysis (GSEA) was performed using gseapy^130^ on gene lists derived from the differential expression analyses, ranked by the limma moderated t-statistic, ordered from most enriched in SW620 to most enriched in SW480. For the human dataset, enrichment was assessed using the Hallmark gene sets from MSigDB^131^ (v2020). For the zebrafish dataset, enrichment was performed using species-appropriate Enrichr^132^ collections (KEGG_2019, WikiPathways_2018, GO_Biological_Process_2018). Preranked GSEA was run with 2,000 permutations and enrichment was defined as FDR < 0.25. Analysis code is available at <u>GitHub</u>.

#### *CD24* knockout SW620 cell line generation

*CD24* knockout (KO) was performed in SW620 cells using a CRISPR/Cas9 kit, following the manufacturer’s instructions (Santa Cruz, sc-418762-KO-2, **Supplementary Figure 4A**). gRNA sequences can be found in **Supplementary Table 1**. To better verify transfection and enable selection of KO cells, plasmids containing RFP and puromycin resistance genes (Santa Cruz, sc-418762-HDR-2) were co-transfected with the GFP-tagged gRNA plasmids, allowing for insertion of these genes through homology-directed DNA repair (**Supplementary Table 2**). Mock cells were subjected to the same transfection protocols, in the absence of plasmids. Twenty–four hours after transfection, successfully double-transfected cells (GFP and RFP positive) were selected by fluorescence-activated cell sorting (FACS), using a BD FACSAria Fusion Cell Sorter (BD Biosciences). Cells were then further selected for successful knock-in by culturing them in puromycin-supplemented (0.75 ug/mL, Santa Cruz) medium for a month, followed by FACS for live (1:5000, DAPI staining) RFP-positive cells. To ensure a pure population, both mock and KO single-cell-derived clones were created using a serial dilution method in 96-well plates, followed by amplification in complete medium supplemented with 20% FBS. Gene editing was validated using quantitative PCR (qPCR) and Western Blot (WB, **Supplementary Figure 4B,C**), confirming complete loss of transcript and protein in two independent clones (KO7 and KO11). To characterize the resulting mutation, clone *CD24* KO11 was further analyzed by Sanger sequencing (**Supplementary Figure 5**) and selected for further experiments.

#### Real-time quantitative PCR

SW620 mock and CD24 KO cells in culture were detached, centrifuged, and resuspended in TRIzol (Thermo Fisher Scientific, #15596026). RNA was isolated using phenol:chloroform (Sigma-Aldrich, #P3803) to extract the aqueous phase, followed by the RNeasy Mini Kit (Qiagen, #74104), following the manufacturer’s instructions. cDNA was synthesized using the Xpert cDNA synthesis kit (GRiSP, #GK80.0100), according to the manufacturer’s instructions, in a MyCycler Thermal Cycler (Bio-Rad). qPCR was performed using the Xpert Fast SYBR Blue mastermix (GRiSP, #GE22.5100), primers (**Supplementary Table 3**, Integrated DNA Technologies) at 0.4 μM, and 100ng of template cDNA per reaction, in a CFX96 Real-Time System C1000 Touch Thermal Cycler (Bio-Rad). Gene expression levels were normalized to the housekeeping gene *EEF1A1*. Relative *CD24* mRNA expression levels were calculated using the 2^−ΔΔCt^ method, where (1) ΔCt equals the cycle threshold value (Ct) of the gene of interest minus the Ct of the housekeeping gene and (2) ΔΔCt equals the ΔCt of the experimental condition (CD24 KO) minus the ΔCt of the control condition (mock).

#### Western Blot

To extract protein from SW620 cells, samples were snap frozen using liquid nitrogen and resuspended in SDS–PAGE 2x sample buffer. Cells were then homogenized using the tissue homogenizer (MM400 TissueLyser, Retsch), incubated with Benzonase for 20 min when required, heated at 50 °C for 10 min, and centrifuged at a maximum speed for 5 min. Protein quantification was measured using the NanoDrop 1000 at 280 nm. Protein extracts were separated with 12% SDS-polyacrylamide gel electrophoresis in running buffer (3.02 g Tris base, 14.42 g glycine, and 1 g SDS in 1 L distilled water) using the Mini-PROTEAN Tetra electrophoresis system (Bio-Rad). Proteins were transferred to PVDF membranes using the Mini Trans-Blot Cell (Bio-Rad) in chilled transfer buffer (5.82 g Tris and 2.93 g glycine in 1 L distilled water), followed by blocking in 5% milk in TBST (20 mM Tris, 150 mM NaCl, 0.1% Tween-20, and distilled water, pH 7.4–7.6) with agitation. Protein loading was verified through GelCode Blue Safe Protein Stain. Membranes were incubated with primary antibodies overnight at 4 °C with agitation, washed with TBST, and incubated with HRP-coupled secondary antibodies for 1 h at room temperature (RT). The antibodies used can be found in **Supplementary Table 4.** Signal was detected using the SuperSignal West Pico Chemiluminescent Substrate HRP, and images were acquired using Amersham Imager 680 RGB (GE Healthcare).

#### PCR and Sanger Sequencing

To confirm CD24 editing, a PCR was performed using the Xpert Fast Hotstart DNA Polymerase (GRiSP, #GE35.0001), according to the manufacturer’s instructions, in a MyCycler Thermal Cycler (Bio-Rad). The PCR amplicon was purified using the GRS PCR & Gel Band Purification Kit (GRiSP) and sequenced by Sanger sequencing (STAB VIDA). The primers used are listed in **Supplementary Table 3** and **Supplementary Figure 5C** shows the sequenced region in green.

### *In vitro* viability and proliferation assay

Sterile round coverslips (VWR, 13mm #1.5) were placed in 12-well plate wells and seeded with SW620 mock or CD24 KO cells (2 x 10^4^ cells/well) in complete cell culture medium (DMEM supplemented with 10% FBS and 1% P/S), which was renewed daily. Four replicates were seeded for each condition, and the plates were incubated at 37°C and 5% CO_2_. After 4 days, the media was discarded, wells were washed with DPBS 1x, fixed with FA 4% for 10 min at RT, and washed again. The cells were then permeabilized with PBS Triton X-100 0.1% (PBSTx) for 5 min at RT and blocked for 1h at RT with PBDX-GS blocking solution (1% bovine serum albumin, 0.05% Triton, 1.5% goat serum, 1% DMSO in DPBS 1X). The cells were then stained with cleaved caspase-3 and phospho-Histone H3 primary antibodies overnight at 4°C, in a humid chamber and protected from light. After washing steps with PBSTx, the cells were then incubated with secondary antibodies and conjugated anti-phalloidin 488, overnight at 4 °C. Finally, cells were counterstained with DAPI (50 μg/ml) for 15min at RT, washed with ddH20, and the coverslips were mounted on top of glass microscope slides using Mowiol mounting medium (Sigma-Aldrich) and stored at 4 °C. The antibodies used can be found in **Supplementary Table 5**. Images were acquired using an Andor BC43 Confocal Microscope (20x objective, 512 x 512 px, FUSION Software, Oxford Instruments). Ten different random fields of view were acquired per coverslip (4 coverslips per condition).

#### Zebrafish macrophage ablation with clodronate liposomes

For selective macrophage depletion, liposome-encapsulated PBS (L-PBS) and liposome-encapsulated clodronate (L-Clodronate, 5 mg/ml) were purchased from Liposoma (CP-005-005). Immediately prior to injection into zebrafish, cells were resuspended in either PBS, L-PBS, or L-Clodronate (100%).

#### Live imaging of zebrafish xenografts and analysis

SW620 mock and CD24 KO xenografts were generated in myeloid-labelling transgenic zebrafish [Tg(mpx:GFP), Tg(csf1ra:GFP), Tg(mpeg1:mCherry), or Tg(mpeg1:mCherry, mpx:GFP)]. At 1 dpi, tricaine-anesthetized xenografts were mounted in 0.8% low-melting agarose in glass–bottom dishes (FluoroDish, WPI) filled with Tricaine 1x in E3. The dish was placed inside the incubator of an Andor BC43 Confocal Microscope (20x objective, Oxford Instruments) at 34°C. Using the Fusion software, Z-stack images were acquired using a 3 µm step size every 3 min. Xenografts were imaged overnight between 1 and 2 dpi for approximately 16h, followed by euthanasia.

Time-lapse movies were analyzed using Fiji^133^ (ImageJ, v1.54r). Individual macrophages and neutrophils were manually tracked using the MTrackJ plugin^134^, allowing export of xyz positions over time, distance to the tumor center, velocity profiles, and track length of each cell. In parallel, tumor contact, fusion, phagocytosis, and myeloid–myeloid contact events were manually annotated at each timepoint of every track. The resulting datasets were analyzed in Python to compute motility, spatial, and functional metrics, which are described in Error! Reference source not found.. In total, 12 xenografts (6 per condition) from 3 independent experiments were analyzed, comprising 140 tracked macrophages and 100 neutrophils.

#### Zebrafish whole-mount immunofluorescence

Xenografts stored in 100% methanol were rehydrated through a decreasing percentage series (75% > 50% > 25% methanol in PBSTx). Next, xenografts were permeabilized with ice-cold acetone at −20°C for 7 min, followed by blocking in PBDX_GS [1X PBS containing 1% (w/v) BSA, 1% (v/v) DMSO, 1.5% (v/v) goat serum, and 0.0005% (v/v) Triton X-100] for 1 h at RT. Staining with primary antibodies was performed overnight, followed by another overnight incubation with secondary antibody and DAPI (50μg/mL) for nuclear counterstaining. Washing was performed between all steps. The antibodies used can be found in **Supplementary Table 6**. Stained xenografts were then mounted between two rectangular coverslips using Mowiol mounting media (Sigma-Aldrich), allowing for double-sided imaging.

#### Imaging and analysis of zebrafish xenografts

All images were obtained using an Andor BC43 Confocal Microscope (20x objective, FUSION Software, Oxford Instruments), in Z-stacks of 5 μm intervals. Generated images were analyzed using the Fiji/ImageJ software. Tumor area was quantified by measuring the area occupied by the fluorescently labeled tumor cells in all slices of the Z-stack of each xenograft. Macrophages and neutrophils were quantified in every slice, using the cell counter plugin, and myeloid cell abundance was quantified as the ratio between the total number of each cell type and the tumor area of each xenograft. Macrophage engulfment events were defined as colocalization of fluorescent tumor cells within macrophages (**Figure 3H**).

#### Glycocalyx analysis

Cells were incubated for 30 min with APC–Cy7–fixable viability dye (FVD) (1:2000, BioLegend), for dead cell exclusion, followed by lectin staining for 15 min on ice: Phaseolus vulgaris leucoagglutinin (L-PHA; Fluorescein); Galanthus nivalis Lectin (GNA; Fluorescein), Concanavalin A (ConA; biotinylated), Sambucus Nigra Lectin (SNA; Cy5), Maackia Amurensis Lectin II (MAL-II; biotinylated), Lycopersicon Esculentum Lectin (LEL; CY5), Aleuria Aurantia Lectin (AAL; biotinylated) and Phaseolus Vulgaris Erythroagglutinin (E-PHA; biotinylated) (1:1000, all from Vector Laboratories). Biotinylated lectins were further incubated with streptavidin (1:200, PE/Cyanine7, BioLegend) for 30 min on ice. For Siglec binding, cells were incubated for 30 min with APC–Cy7–fixable viability dye (FVD) (1:2000, BioLegend), for dead cell exclusion, followed by incubation with recombinant proteins for 20 min on ice: rSiglec-3 (1:300), rSiglec-10 (1:200) and rSiglec-15 (1:600). Cells were then incubated with a secondary antibody, 6x-His Tag Monoclonal Antibody (1:1000; clone: 4E3D10H2/E3; Invitrogen) for 30 min on ice. Data acquisition was performed using a fluorescence-activated cell sorting (FACS) Canto II flow cytometer (BD Biosciences) using the FACSDiva software (BD Biosciences). Data were analyzed using the FlowJo v10 software (TreeStar Inc.).

#### Phylogenetic and comparative genomic analyses of CD24 and CD24 candidate sequences

Phylogenetic relationships and divergence times among the vertebrate species examined were obtained using TimeTree 5^135^ with default settings. The resulting tree was exported in Newick format (.nwk) and imported into BigTreeViewer^136^ for visualization and figure generation. Animal silhouettes obtained from PhyloPic were subsequently added.

Comparative synteny analysis was based on the genomic locations of the genes catalogued in **Supplementary Data 3**. In species with an annotated *CD24* gene, *CD24* served as the reference locus, and the ten nearest annotated genes on either side were catalogued in genomic coordinate order. In species without an annotated *CD24* gene, the corresponding candidate interval was identified using *QRSL1* as the left flanking anchor and *MTRES1* or *IARS2* as the right flanking anchor, and ten flanking genes on each side were catalogued. Selected genes informative for conserved synteny and candidate locus identification were depicted schematically in Microsoft PowerPoint. Corresponding genes were assigned matching colors, and arrowheads indicated transcriptional orientation. Genomic intervals were not drawn to scale, and the distance between the conserved flanking regions was indicated for each species.

Candidate intervals were examined using gene and transcript annotation tracks in the UCSC Genome Browser or Ensembl. Transcript models, including lncRNAs and lincRNAs, were inspected for exon numbers, lengths, and arrangements resembling those of annotated *CD24* genes. Candidate DNA sequences were extracted, and putative open reading frames (ORFs) were reconstructed and translated for protein sequence analysis. The candidate ORF from *Callorhinchus milii* was manually curated by adjusting its reading frame, guided by amino acid sequence similarity to the other CD24 and candidate CD24-like sequences. This sequence was treated as a putative reconstruction. Accession numbers and stable identifiers for the selected proteins and candidate loci are provided in **Supplementary Table 7**.

#### Protein sequence and structural analyses of CD24 and CD24 candidate sequences

Amino acid sequences of annotated CD24 proteins and translated candidate ORFs were aligned using Clustal Omega through the EMBL-EBI Job Dispatcher web server with default alignment parameters^137,138^. The alignment was imported into Lasergene MegAlign Pro 19.0.2 (DNASTAR) for visualization and figure preparation. A consensus threshold of >50% was applied, and residues matching the consensus sequence were highlighted.

Protein sequence features were assessed using SignalP 6.0^139^, DeepTMHMM 1.0^140^, and NetGPI 1.1^141^ with default prediction settings to predict N-terminal signal peptides, transmembrane regions, and GPI-anchoring signals, respectively. Predictions were summarized as the presence or absence of signal peptides and GPI-anchoring signals, together with the number of predicted transmembrane regions.

Comparative protein structure modelling was attempted for selected CD24 and candidate CD24-like amino acid sequences using the SWISS-MODEL web server with default settings^142–146^. Structural models and associated membrane-positioning visualizations were used for figure preparation. Sequences for which no structural model was obtained were designated “N.A.”.

#### Bioinformatic identification of putative zebrafish Siglec-10 orthologs

Protein sequences for human SIGLEC10 and zebrafish putative orthologs were retrieved from UniProt (**Supplementary Table 8**, **Supplementary Table 9**). Both the full protein sequence of human SIGLEC10 and, due to its relevance for CD24 binding, its Ig-like V-type domain were used as queries for BLASTp searches against the *Danio rerio* proteome. Human-zebrafish orthology relationships were independently assessed using the HGNC Comparison of Orthology Predictions (HCOP) tool (https://www.genenames.org/tools/hcop)^147^. Three zebrafish ortholog candidates - mag, siglec15l, and si:dkey-24p1.7 - were prioritized based on BLASTp sequence similarity and conservation of IgV domain and the sialic-acid binding arginine. A multiple sequence alignment of the human SIGLEC10 IgV domain and the full protein sequences of the zebrafish candidates was performed using the EMBL-EBI Clustal Omega MSA platform (https://www.ebi.ac.uk/jdispatcher/msa/clustalo)^138^ and imported into Jalview 2.11.5.0^148^ for visualization. Jalview was used to calculate residue consensus and conservation, and to generate the final figure. Single-cell RNA sequencing datasets of zebrafish larvae at 2, 3, and 5 dpf were obtained from Zebrahub^58^. Clusters corresponding to macrophages were identified based on Zebrahub annotation and high expression of known macrophage marker genes (*mpeg1.1*, *csf1ra*, *mfap4*). The expression of the candidates (*siglec15l*, *mag*, *si:dkey-24p1.7*) was then examined across clusters. All analyses were performed in Python and figures were generated using matplotlib^149^. Analysis code is available at <u>GitHub</u>.

#### TCGA Survival Analysis

Publicly available bulk RNA sequencing and clinical data for colon adenocarcinoma (COAD) and rectal adenocarcinoma (READ) patients were obtained from the UCSC Xena Browser (https://xenabrowser.net/, in November 2025). Gene-level expression was taken from the GDC STAR-derived FPKM-UQ matrices (log2(FPKM-UQ+1)) for TCGA-COAD and TCGA-READ, and survival endpoints from the TCGA Pan-Cancer Clinical Data Resource^150^. Expression and clinical datasets were harmonized and merged at the patient level and restricted to primary tumor samples. For patients with multiple samples available, a single representative sample was retained to ensure statistical independence, and if a patient had multiple clinical records with discordant staging, the highest reported AJCC pathological stage was retained. Patients lacking AJCC pathological stage, CD24 expression, or overall survival data were excluded, yielding 601 unique patients (COAD n=444, READ n=157, **Supplementary Table 10**). CD24 expression was extracted using Ensembl identifier ENSG00000272398 and, within each pathological stage cohort (I–IV), patients were stratified into CD24-high and CD24-low groups by a median split. Overall survival (OS) and progression-free interval (PFI) were censored at 5 years to limit bias from sparse long-term follow-up, estimated by Kaplan-Meier analysis, and compared between stratification groups using log-rank tests. Hazard ratios (HR) were estimated by univariate Cox proportional hazards regression, and non-convergent or implausibly large estimates (HR > 50) arising from low event counts were not reported. Analyses were conducted in Python using pandas^151^, lifelines^152^, and matplotlib^149^. Analysis code is available at <u>GitHub</u>.

#### Statistical Analysis

Unless stated otherwise, statistical analysis was performed using the GraphPad Prism software version 10.4.1. The ROUT method was used with a Q=1% to identify outliers, which were excluded from further analysis. Data normality was assessed by the D’Agostino & Pearson or the Shapiro-Wilk tests. In general, data sets with a Gaussian distribution were analyzed by parametric unpaired two-tailed t-test or ordinary one-way ANOVA with multiple comparisons, whereas the sets that did not pass the normality tests were analyzed by non-parametric unpaired Mann–Whitney U test. Most often, each experimental data set was compared to the respective control, and two-sided tests had a confidence interval of 95%. Statistical differences were considered significant whenever P-value (*P*)<0.05 and statistical output was represented by stars as follows: non-significant (*ns*)>0.05, \**P*<0.05; \*\**P*<0.01; \*\*\**P*<0.001; \*\*\*\**P*<0.0001. All graphs present the data as average ± standard deviation (SD).

## Data Availability

Output data from bulk RNAseq and TCGA analysis are available as Supplementary Data. Analysis code, and relevant software and package versions are available at <u>GitHub</u> and archived at Zenodo (DOI: <u>10.5281/zenodo.22084075</u>). Time-lapse microscopy movies are available through the BioImage Archive (accession <u>S-BIAD3811</u>).

## Acknowledgements

We are grateful to all members of the Fior Lab for their support and critical discussion. To the Molecular Mechanisms of Disease Group at CE3C (Ana Rita Carlos, Diogo Fernandes, Rita Zilhão) for their collaboration and support in the validation of the CD24 KO cell line. To the Immunology, Cancer & GlycoMedicine Group at i3S (Catarina Azevedo, Salomé Pinho) for their collaboration and support with glycobiology experiments. To Dr. António Gomes from Hospital Prof. Doutor Fernando Fonseca for their support in the TCGA analysis. To the Champalimaud Foundation (CF) Fish Platform (Catarina Certal, Joana Monteiro and team) for excellent animal care and the CF Molecular and Transgenic Tools Platform (Raquel Tomás) for their technical support. We are also grateful to the zebrafish community for sharing zebrafish strains (S. Renshaw, F. Djouad, and Z. Wen).

## Author contributions

R.F. conceptualized and supervised the research; RNAseq was performed by V.P.; experiments performed by A.B.M., C.R.A., C.M.A., D.R.F, and F.M.; data analysis performed by A.B.M., C.M.A, N.V., and I.T.; J.E.-O. and J.J.B. provided recombinant Siglec proteins; manuscript writing by A.B.M., I.T., and R.F. with revisions by C.M.A and S.S.P. All authors contributed with critical reading of the manuscript. All authors agreed to the published version of the manuscript.

## References

1. Sharma, P. & Allison, J. P. The future of immune checkpoint therapy. Science 348, 56–61 (2015).

2. Sharma, P., Hu-Lieskovan, S., Wargo, J. A. & Ribas, A. Primary, Adaptive, and Acquired Resistance to Cancer Immunotherapy. Cell 168, 707–723 (2017).

3. Fridman, W. H., Zitvogel, L., Sautès–Fridman, C. & Kroemer, G. The immune contexture in cancer prognosis and treatment. Nat Rev Clin Oncol 14, 717–734 (2017).

4. Demaria, O. et al. Harnessing innate immunity in cancer therapy. Nature 574, 45– 56 (2019).

5. Khalaji, A. et al. Don’t eat me/eat me signals as a novel strategy in cancer immunotherapy. Heliyon 9, (2023).

6. Liu, X., Kwon, H., Li, Z. & Fu, Y. Is CD47 an innate immune checkpoint for tumor evasion? Journal of Hematology & Oncology 10, 12 (2017).

7. Majeti, R. et al. CD47 Is an Adverse Prognostic Factor and Therapeutic Antibody Target on Human Acute Myeloid Leukemia Stem Cells. Cell 138, 286–299 (2009).

8. Tseng, D. et al. Anti-CD47 antibody-mediated phagocytosis of cancer by macrophages primes an effective antitumor T-cell response. Proc Natl Acad Sci U S A 110, 11103–11108 (2013).

9. Xu, Y., Jiang, P., Xu, Z. & Ye, H. Opportunities and challenges for anti-CD47 antibodies in hematological malignancies. Front. Immunol. 15, (2024).

10. Wilde, L. & Kasner, M. Targeting CD47: many misses; hopeful for a hit. Blood 145, 460–462 (2025).

11. Fior, R. et al. Single-cell functional and chemosensitive profiling of combinatorial colorectal therapy in zebrafish xenografts. Proceedings of the National Academy of Sciences 114, E8234–E8243 (2017).

12. Jin, H. et al. Definitive hematopoietic stem/progenitor cells manifest distinct differentiation output in the zebrafish VDA and PBI. Development 136, 1397 (2009).

13. Soza-Ried, C., Hess, I., Netuschil, N., Schorpp, M. & Boehm, T. Essential role of c-myb in definitive hematopoiesis is evolutionarily conserved. Proceedings of the National Academy of Sciences 107, 17304–17308 (2010).

14. Gut, P., Reischauer, S., Stainier, D. Y. R. & Arnaout, R. Little Fish, Big Data: Zebrafish as a Model for Cardiovascular and Metabolic Disease. Physiological Reviews 97, 889–938 (2017).

15. Renshaw, S. A. & Trede, N. S. A model 450 million years in the making: zebrafish and vertebrate immunity. Dis Model Mech 5, 38–47 (2012).

16. Gamble, J. T., Elson, D. J., Greenwood, J. A., Tanguay, R. L. & Kolluri, S. K. The Zebrafish Xenograft Models for Investigating Cancer and Cancer Therapeutics. Biology 10, 252 (2021).

17. Fontana, C. M. & Van Doan, H. Zebrafish xenograft as a tool for the study of colorectal cancer: a review. Cell Death Dis 15, 23 (2024).

18. Barbosa, G. R. et al. Zebrafish as a Model for Translational Immuno-Oncology. Journal of Personalized Medicine 15, 304 (2025).

19. Leibovitz, A. et al. Classification of human colorectal adenocarcinoma cell lines. Cancer Res 36, 4562–4569 (1976).

20. Hewitt, R. E. et al. Validation of a model of colon cancer progression. The Journal of Pathology 192, 446–454 (2000).

21. Póvoa, V. et al. Innate immune evasion revealed in a colorectal zebrafish xenograft model. Nat Commun 12, 1156 (2021).

22. Pirruccello, S. J. & LeBien, T. W. The human B cell-associated antigen CD24 is a single chain sialoglycoprotein. J Immunol 136, 3779–3784 (1986).

23. Kristiansen, G. et al. CD24 Is Expressed in Ovarian Cancer and Is a New Independent Prognostic Marker of Patient Survival. Am J Pathol 161, 1215–1221 (2002).

24. Sagiv, E. et al. CD24 Is a New Oncogene, Early at the Multistep Process of Colorectal Cancer Carcinogenesis. Gastroenterology 131, 630–639 (2006).

25. Chen, G.-Y., Tang, J., Zheng, P. & Liu, Y. CD24 and Siglec-10 selectively repress tissue damage-induced immune responses. Science 323, 1722–1725 (2009).

26. Macauley, M. S., Crocker, P. R. & Paulson, J. C. Siglec-mediated regulation of immune cell function in disease. Nat Rev Immunol 14, 653–666 (2014).

27. Yin, S.-S. & Gao, F.-H. Molecular Mechanism of Tumor Cell Immune Escape Mediated by CD24/Siglec-10. Front. Immunol. 11, (2020).

28. Barkal, A. A. et al. CD24 signalling through macrophage Siglec-10 is a target for cancer immunotherapy. Nature 572, 392–396 (2019).

29. Liu, Y. et al. Neutrophil extracellular traps impede cancer metastatic seeding via protease-activated receptor 2-mediated downregulation of phagocytic checkpoint CD24. J Immunother Cancer 13, (2025).

30. Wei, Y. et al. KRAS Inhibition Activates an Actionable CD24 “Do Not Eat Me” Signal in Pancreatic Cancer. Cancer Res 85, 4825–4838 (2025).

31. Aigner, S. et al. CD24 mediates rolling of breast carcinoma cells on P-selectin. The FASEB Journal 12, 1241–1251 (1998).

32. Mittelheisser, V. et al. Tumoral CD24 tunes platelets binding and pro-metastatic functions. 2025.04.09.648049 Preprint at 10.1101/2025.04.09.648049 (2025).

33. Myung, J. H. et al. Direct Measurements on CD24-Mediated Rolling of Human Breast Cancer MCF-7 Cells on E-selectin. Anal Chem 83, 1078–1083 (2011).

34. Kadmon, G., Kowitz, A., Altevogt, P. & Schachner, M. The neural cell adhesion molecule N-CAM enhances L1-dependent cell-cell interactions. J Cell Biol 110, 193–208 (1990).

35. Kleene, R., Yang, H., Kutsche, M. & Schachner, M. The Neural Recognition Molecule L1 Is a Sialic Acid-binding Lectin for CD24, Which Induces Promotion and Inhibition of Neurite Outgrowth *. Journal of Biological Chemistry 276, 21656–21663 (2001).

36. Runz, S. et al. CD24 induces localization of β1 integrin to lipid raft domains. Biochemical and Biophysical Research Communications 365, 35–41 (2008).

37. Su, N. et al. Lyn is involved in CD24-induced ERK1/2 activation in colorectal cancer. Mol Cancer 11, 43 (2012).

38. Schabath, H., Runz, S., Joumaa, S. & Altevogt, P. CD24 affects CXCR4 function in pre-B lymphocytes and breast carcinoma cells. J Cell Sci 119, 314–325 (2006).

39. Pinho, S. S. & Reis, C. A. Glycosylation in cancer: mechanisms and clinical implications. Nat Rev Cancer 15, 540–555 (2015).

40. Pinho, S. S., Macauley, M. S. & Läubli, H. Tumor glyco-immunology, glyco-immune checkpoints and immunotherapy. J Immunother Cancer 13, (2025).

41. Alves, I., Fernandes, Â., Santos-Pereira, B., Azevedo, C. M. & Pinho, S. S. Glycans as a key factor in self and nonself discrimination: impact on the breach of immune tolerance. FEBS Letters 596, 1485–1502 (2022).

42. O’Neill, A. et al. Stromal cells modulate innate immune cell phenotype and function in colorectal cancer via the Sialic acid/Siglec axis. J Immunother Cancer 13, (2025).

43. Seagen Inc. *A Phase 1 Study of SGN-STNV in Advanced Solid Tumors*. https://clinicaltrials.gov/study/NCT04665921 (2025).

44. National Cancer Institute (NCI). *Phase III Randomized Study of Chimeric Antibody 14.18 (Ch14.18) in High Risk Neuroblastoma Following Myeloablative Therapy and Autologous Stem Cell Rescue*. https://clinicaltrials.gov/study/NCT00026312 (2025).

45. St. Anna Kinderkrebsforschung. *High Risk Neuroblastoma Study 1 of SIOP-Europe (SIOPEN)*. https://clinicaltrials.gov/study/NCT01704716 (2020).

46. OBI Pharma, Inc. A Phase 1/2, Open-Label, Dose-Escalation and Cohort-Expansion Study Evaluating the Safety, Pharmacokinetics, and Therapeutic Activity of OBI-999 in Patients With Advanced Solid Tumors. https://clinicaltrials.gov/study/NCT04084366 (2025).

47. Hoff, P. M. G. *A Multicenter Phase II Study of Treatment With Hu3S193 in Women With Advanced Breast Cancer That Progressed After Hormonal Therapy*. https://clinicaltrials.gov/study/NCT01370239 (2019).

48. Palleon Pharmaceuticals, Inc. *A Phase 1/2, Open-Label, Single-Arm, Dose-Escalation and Dose-Expansion Study of the Safety, Tolerability, Pharmacokinetic, and Antitumor Activity of E-602 as a Single Agent and in Combination With Cemiplimab in Patients With Advanced Cancers*. https://clinicaltrials.gov/study/NCT05259696 (2025).

49. Gettinger, S. *The Study of NC318 Alone or in Combination With Pembrolizumab in Patients With Advanced Non-Small Cell Lung Cancer*. https://clinicaltrials.gov/study/NCT04699123 (2025).

50 . Galecto Biotech AB. *An Open Label Study Followed by a Randomised, Double-Blind, Placebo-Controlled, Parallel Group and an Extension Study to Investigate the Safety and Efficacy of GB1211 (a Galectin-3 Inhibitor) in Combination With Atezolizumab in Patients With Non-Small Cell Lung Cancer (NSCLC)*. https://clinicaltrials.gov/study/NCT05240131 (2024).

51. Li, S. et al. IMM47, a humanized monoclonal antibody that targets CD24, exhibits exceptional anti-tumor efficacy by blocking the CD24/Siglec-10 interaction and can be used as monotherapy or in combination with anti-PD1 antibodies for cancer immunotherapy. Antibody Ther 6, 240–252 (2023).

52. Antengene Biologics Limited. *A First-in-Human Phase I Study of ATG-031 in Patients With Advanced Solid Tumors or B-Cell Non-Hodgkin Lymphomas*. https://clinicaltrials.gov/study/NCT06028373 (2025).

53. Pheast Therapeutics. *An Open-Label, Phase 1a/1b, Dose Escalation and Dose Expansion Study Investigating the Safety, Pharmacokinetics, Pharmacodynamics, and Antitumor Activity of PHST001 in Adult Patients With Advanced Relapsed and/or Refractory Solid Tumors*. https://clinicaltrials.gov/study/NCT06840886 (2025).

54. Martinez-Lopez, M., Póvoa, V. & Fior, R. Generation of Zebrafish Larval Xenografts and Tumor Behavior Analysis. Journal of Visualized Experiments (JoVE) e62373 (2021).

55. Cao, H. et al. Red blood cell mannoses as phagocytic ligands mediating both sickle cell anaemia and malaria resistance. Nat Commun 12, 1792 (2021).

56. Ayre, D. C. et al. Analysis of the structure, evolution, and expression of CD24, an important regulator of cell fate. Gene 590, 324–337 (2016).

57. Mazar, J. et al. Zika virus as an oncolytic treatment of human neuroblastoma cells requires CD24. PLOS ONE 13, e0200358 (2018).

58. Lange, M. et al. A multimodal zebrafish developmental atlas reveals the state-transition dynamics of late-vertebrate pluripotent axial progenitors. Cell 187, 6742–6759.e17 (2024).

59. Fang, J. et al. Targeting the CD24-Siglec10 Axis: A Potential Strategy for Cancer Immunotherapy. BIO Integration 5, 997 (2024).

60. Tsuchiya, H. et al. Immune evasion from macrophages by NEAT1-induced CD24 in liver cancer. Oncogene 44, 3652–3664 (2025).

61. Zhu, B. et al. PPDPF promotes the progression of esophageal squamous cell carcinoma via c-Myc/CD24 axis. J Immunother Cancer 13, (2025).

62. Khatib-Massalha, E. et al. Defective neutrophil clearance in JAK2V617F myeloproliferative neoplasms drives myelofibrosis via immune checkpoint CD24. Blood 146, 717–731 (2025).

63. Zhao, K. et al. From mechanism to therapy: the journey of CD24 in cancer. Front. Immunol. 15, (2024).

64. Balkwill, F. Tumour necrosis factor and cancer. Nat Rev Cancer 9, 361–371 (2009).

65. Kratochvill, F. et al. TNF Counterbalances the Emergence of M2 Tumor Macrophages. Cell Reports 12, 1902–1914 (2015).

66. Martínez-López, M. F. et al. Macrophages directly kill bladder cancer cells through TNF signaling as an early response to BCG therapy. Dis Model Mech 17, dmm050693 (2024).

67. Chan, S.-H., Lin, C.-Y., Tseng, H.-J. & Wang, L.-H. CD24a knockout results in an enhanced macrophage- and CD8^+^ T cell-mediated anti-tumor immune responses in tumor microenvironment in a murine triple-negative breast cancer model. J Biomed Sci 32, 73 (2025).

68. Zhang, Y. et al. Targeted release of a bispecific fusion protein SIRPα/Siglec-10 by oncolytic adenovirus reinvigorates tumor-associated macrophages to improve therapeutic outcomes in solid tumors. J Immunother Cancer 13, (2025).

69. Pagán, A. J. & Ramakrishnan, L. The Formation and Function of Granulomas. Annual Review of Immunology 36, 639–665 (2018).

70. Kleinmanns, K., Fosse, V., Bjørge, L. & McCormack, E. The Emerging Role of CD24 in Cancer Theranostics-A Novel Target for Fluorescence Image-Guided Surgery in Ovarian Cancer and Beyond. J Pers Med 10, 255 (2020).

71. Grandchamp, A., Piégu, B. & Monget, P. Genes Encoding Teleost Fish Ligands and Associated Receptors Remained in Duplicate More Frequently than the Rest of the Genome. Genome Biol Evol 11, 1451–1462 (2019).

72. Chen, G.-Y. et al. Preserving Sialic Acid-dependent Pattern Recognition by CD24-Siglec G Interaction for Therapy of Polybacterial Sepsis. Nat Biotechnol 29, 428–435 (2011).

73. Harris, E. S. et al. Reduced sialylation of airway mucin impairs mucus transport by altering the biophysical properties of mucin. Sci Rep 14, 16568 (2024).

74. Mantuano, N. R. & Läubli, H. Sialic acid and Siglec receptors in tumor immunity and immunotherapy. Seminars in Immunology 74–75, 101893 (2024).

75. Alves, I. et al. Host-derived mannose glycans trigger a pathogenic γδ T cell/IL-17a axis in autoimmunity. Science Translational Medicine 15, eabo1930 (2023).

76. Bergmiller, T., Ackermann, M. & Silander, O. K. Patterns of Evolutionary Conservation of Essential Genes Correlate with Their Compensability. PLoS Genet 8, e1002803 (2012).

77. Bernheim, A., Cury, J. & Poirier, E. Z. The immune modules conserved across the tree of life: Towards a definition of ancestral immunity. PLoS Biol 22, e3002717 (2024).

78. Lei, Z., et al. Understanding and targeting resistance mechanisms in cancer. MedComm (2020) 4, e265 (2023).

79. Baldassarre, G., L. de la Serna, I. & Vallette, F. M. Death-ision: the link between cellular resilience and cancer resistance to treatments. Molecular Cancer 24, 144 (2025).

80. Lv, W. et al. The drug target genes show higher evolutionary conservation than non-target genes. Oncotarget 7, 4961–4971 (2015).

81. Varki, A. & Angata, T. Siglecs—the major subfamily of I-type lectins. Glycobiology 16, 1R–27R (2006).

82. Angata, T. & Varki, A. Discovery, classification, evolution and diversity of Siglecs. Molecular Aspects of Medicine 90, 101117 (2023).

83. Angata, T. Possible Influences of Endogenous and Exogenous Ligands on the Evolution of Human Siglecs. Front. Immunol. 9, (2018).

84. Lehmann, F., Gäthje, H., Kelm, S. & Dietz, F. Evolution of sialic acid–binding proteins: molecular cloning and expression of fish siglec-4. Glycobiology 14, 959–968 (2004).

85. Angata, T., Tabuchi, Y., Nakamura, K. & Nakamura, M. Siglec-15: an immune system Siglec conserved throughout vertebrate evolution. Glycobiology 17, 838–846 (2007).

86. Bornhöfft, K. F. et al. Characterization of Sialic Acid-Binding Immunoglobulin-Type Lectins in Fish Reveals Teleost-Specific Structures and Expression Patterns. Cells 9, 836 (2020).

87. Weichert, W. et al. Cytoplasmic CD24 Expression in Colorectal Cancer Independently Correlates with Shortened Patient Survival. Clin Cancer Res 11, 6574– 6581 (2005).

88. Choi, Y. L. et al. Enhanced CD24 Expression in Colorectal Cancer Correlates with Prognostic Factors. J Pathol Transl Med 40, 103–111 (2006).

89. Nersisyan, S. et al. Low expression of CD24 is associated with poor survival in colorectal cancer. Biochimie 192, 91–101 (2022).

90. Choi, D. et al. Cancer stem cell markers CD133 and CD24 correlate with invasiveness and differentiation in colorectal adenocarcinoma. World Journal of Gastroenterology 15, 2258–2264 (2009).

91. Wang, W. et al. CD24-dependent MAPK pathway activation is required for colorectal cancer cell proliferation. Cancer Sci 101, 112–119 (2010).

92. Chen, Z. et al. Antibody-based targeting of CD24 enhances antitumor effect of cetuximab via attenuating phosphorylation of Src/STAT3. Biomed Pharmacother 90, 427– 436 (2017).

93. Wang, X. et al. CD24 promoted cancer cell angiogenesis via Hsp90-mediated STAT3/VEGF signaling pathway in colorectal cancer. Oncotarget 7, 55663–55676 (2016).

94. Okano, M. et al. Human colorectal CD24+ cancer stem cells are susceptible to epithelial-mesenchymal transition. International Journal of Oncology 45, 575–580 (2014).

95. Vermeulen, L. et al. Single-cell cloning of colon cancer stem cells reveals a multi-lineage differentiation capacity. Proceedings of the National Academy of Sciences 105, 13427–13432 (2008).

96. Yeo, M.-K. et al. Up-regulation of Cytoplasmic CD24 Expression Is Associated with Malignant Transformation but Favorable Prognosis of Colorectal Adenocarcinoma. Anticancer Res 36, 6593–6598 (2016).

97. Lubeseder-Martellato, C. et al. Membranous CD24 drives the epithelial phenotype of pancreatic cancer. Oncotarget 7, 49156–49168 (2016).

98. Majores, M. et al. Membranous CD24 expression as detected by the monoclonal antibody SWA11 is a prognostic marker in non-small cell lung cancer patients. BMC Clin Pathol 15, 19 (2015).

99. Duex, J. E. et al. Nuclear CD24 Drives Tumor Growth and Is Predictive of Poor Patient Prognosis. Cancer Res 77, 4858–4867 (2017).

100. Ahmed, M. a. H., et al. CD24 shows early upregulation and nuclear expression but is not a prognostic marker in colorectal cancer. Journal of Clinical Pathology 62, 1117– 1122 (2009).

101. Paschall, A. V. et al. CD133 + CD24 lo defines a 5-Fluorouracil-resistant colon cancer stem cell-like phenotype. Oncotarget 7, 78698–78712 (2016).

102. Kong, W. et al. Chemoradiotherapy facilitates siglec-10+/siglec-9+ macrophage-mediated impairment of CD24+/MUC16+ tumor cell elimination and enhances PD-L2 dependent immunosuppression in cervical cancer. Cancer Immunol Immunother 75, 97 (2026).

103. ImmuneOnco Biopharmaceuticals (Shanghai) Inc. *A Phase I, Open-Label, Multicenter, Dose-Escalation And Cohort-Expansion Study Of IMM47 In Subjects With Advanced Solid Tumors*. https://clinicaltrials.gov/study/NCT05985083 (2023).

104. Ozawa, Y. et al. CD24, not CD47, negatively impacts upon response to PD-1/L1 inhibitors in non–small-cell lung cancer with PD-L1 tumor proportion score < 50. Cancer Science 112, 72–80 (2021).

105. Ji, L. et al. CD24 Is a Superior Immunotherapeutic Target to PD-1 in a Mouse Model of Helicobacter-Induced Gastric Cancer. Gastro Hep Advances 1, 79–82 (2022).

106. Zhang, W. et al. An in-situ peptide-antibody self-assembly to block CD47 and CD24 signaling enhances macrophage-mediated phagocytosis and anti-tumor immune responses. Nat Commun 15, 5670 (2024).

107. Yang, Y. et al. PPAB001, a novel bispecific antibody against CD47 and CD24, enhances anti-PD-L1 efficacy in triple-negative breast cancer via reprogramming tumor-associated macrophages towards M1 phenotype. International Immunopharmacology 144, 113740 (2025).

108. Nan, L. et al. Harnessing the innate immune system: a novel bispecific antibody targeting CD47 and CD24 for selective tumor clearance. J Immunother Cancer 13, (2025).

109. Zhou, W., Huang, X., Jia, G., Lu, C. & Ma, W. CD24 as an innate immune checkpoint in solid tumors: biology, biomarker stratification, and therapeutic translation. Front. Immunol. 17, (2026).

110. MacRae, C. A. & Peterson, R. T. Zebrafish as tools for drug discovery. Nat Rev Drug Discov 14, 721–731 (2015).

111. Zanandrea, R., Bonan, C. D. & Campos, M. M. Zebrafish as a model for inflammation and drug discovery. Drug Discovery Today 25, 2201–2211 (2020).

112. Sturtzel, C. et al. Refined high-content imaging-based phenotypic drug screening in zebrafish xenografts. *npj Precis*. Onc. 7, 44 (2023).

113. Zhan, T. et al. Zebrafish live imaging: a strong weapon in anticancer drug discovery and development. Clin Transl Oncol 26, 1807–1835 (2024).

114. Rebelo de Almeida, C., et al. Zebrafish xenografts as a fast screening platform for bevacizumab cancer therapy. Commun Biol 3, 299 (2020).

115. Costa, B. et al. Zebrafish Avatar-test forecasts clinical response to chemotherapy in patients with colorectal cancer. Nat Commun 15, 4771 (2024).

116. Mendes, R. V. et al. Zebrafish Avatar testing preclinical study predicts chemotherapy response in breast cancer. *npj Precis*. Onc. 9, 94 (2025).

117. Estrada, M. F., et al. zAvatar-test—A functional precision model to personalize ovarian cancer treatments: Results from a co-clinical study. CR Med 7, (2026).

118. Renshaw, S. A. et al. A transgenic zebrafish model of neutrophilic inflammation. Blood 108, 3976–3978 (2006).

119. Ellett, F., Pase, L., Hayman, J. W., Andrianopoulos, A. & Lieschke, G. J. mpeg1 promoter transgenes direct macrophage-lineage expression in zebrafish. Blood 117, e49– e56 (2011).

120. Dee, C. T. et al. CD4-Transgenic Zebrafish Reveal Tissue-Resident Th2- and Regulatory T Cell–like Populations and Diverse Mononuclear Phagocytes. J Immunol 197, 3520–3530 (2016).

121. Nguyen-Chi, M. et al. Identification of polarized macrophage subsets in zebrafish. eLife 4, e07288 (2015).

122. Lawson, N. D. & Weinstein, B. M. *In Vivo* Imaging of Embryonic Vascular Development Using Transgenic Zebrafish. Developmental Biology 248, 307–318 (2002).

123. Andrews, S. FastQC A Quality Control tool for High Throughput Sequence Data. https://www.bioinformatics.babraham.ac.uk/projects/fastqc/.

124. Patro, R., Duggal, G., Love, M. I., Irizarry, R. A. & Kingsford, C. Salmon provides fast and bias-aware quantification of transcript expression. Nat Methods 14, 417–419 (2017).

125. Soneson, C., Love, M. I. & Robinson, M. D. Differential analyses for RNA-seq: transcript-level estimates improve gene-level inferences. F1000Res 4, 1521 (2016).

126. Robinson, M. D., McCarthy, D. J. & Smyth, G. K. edgeR: a Bioconductor package for differential expression analysis of digital gene expression data. Bioinformatics 26, 139– 140 (2010).

127. Ritchie, M. E. et al. limma powers differential expression analyses for RNA-sequencing and microarray studies. Nucleic Acids Res 43, e47 (2015).

128. Law, C. W., Chen, Y., Shi, W. & Smyth, G. K. voom: Precision weights unlock linear model analysis tools for RNA-seq read counts. Genome Biol 15, R29 (2014).

129. McCarthy, D. J. & Smyth, G. K. Testing significance relative to a fold-change threshold is a TREAT. Bioinformatics 25, 765–771 (2009).

130. Fang, Z., Liu, X. & Peltz, G. GSEApy: a comprehensive package for performing gene set enrichment analysis in Python. Bioinformatics 39, btac757 (2023).

131. Liberzon, A. et al. The Molecular Signatures Database (MSigDB) hallmark gene set collection. Cell Syst 1, 417–425 (2015).

132. Kuleshov, M. V. et al. Enrichr: a comprehensive gene set enrichment analysis web server 2016 update. Nucleic Acids Res 44, W90–97 (2016).

133. Schneider, C. A., Rasband, W. S. & Eliceiri, K. W. NIH Image to ImageJ: 25 years of image analysis. Nat. Methods 9, 671–675 (2012).

134. Meijering, E., Dzyubachyk, O. & Smal, I. Methods for cell and particle tracking. Methods Enzymol 504, 183–200 (2012).

135. Kumar, S. et al. TimeTree 5: An Expanded Resource for Species Divergence Times. Mol Biol Evol 39, msac174 (2022).

136. Allard, J. B. & Kumar, S. BigTreeViewer: A browser-based viewer and figure renderer for ultra-large phylogenetic trees. In preparation (2026).

137. Sievers, F. et al. Fast, scalable generation of high-quality protein multiple sequence alignments using Clustal Omega. Mol Syst Biol 7, MSB201175 (2011).

138. Madeira, F. et al. The EMBL-EBI Job Dispatcher sequence analysis tools framework in 2024. Nucleic Acids Res 52, W521–W525 (2024).

139. Teufel, F. et al. SignalP 6.0 predicts all five types of signal peptides using protein language models. Nat Biotechnol 40, 1023–1025 (2022).

140. Hallgren, J. et al. DeepTMHMM predicts alpha and beta transmembrane proteins using deep neural networks. 2022.04.08.487609 Preprint at 10.1101/2022.04.08.487609 (2022).

141. Gíslason, M. H., Nielsen, H., Almagro Armenteros, J. J. & Johansen, A. R. Prediction of GPI-anchored proteins with pointer neural networks. Current Research in Biotechnology 3, 6–13 (2021).

142. Guex, N., Peitsch, M. C. & Schwede, T. Automated comparative protein structure modeling with SWISS-MODEL and Swiss-PdbViewer: A historical perspective. ELECTROPHORESIS 30, S162–S173 (2009).

143. Bienert, S. et al. The SWISS-MODEL Repository—new features and functionality. Nucleic Acids Res 45, D313–D319 (2017).

144. Bertoni, M., Kiefer, F., Biasini, M., Bordoli, L. & Schwede, T. Modeling protein quaternary structure of homo- and hetero-oligomers beyond binary interactions by homology. Sci Rep 7, 10480 (2017).

145. Waterhouse, A. et al. SWISS-MODEL: homology modelling of protein structures and complexes. Nucleic Acids Res 46, W296–W303 (2018).

146. Studer, G. et al. QMEANDisCo—distance constraints applied on model quality estimation. Bioinformatics 36, 1765–1771 (2020).

147. Seal, R. L. et al. Genenames.org: the HGNC resources in 2023. Nucleic Acids Res 51, D1003–D1009 (2023).

148. Waterhouse, A. M., Procter, J. B., Martin, D. M. A., Clamp, M. & Barton, G. J. Jalview Version 2—a multiple sequence alignment editor and analysis workbench. Bioinformatics 25, 1189–1191 (2009).

149. Hunter, J. Matplotlib: A 2D Graphics Environment. Computing in Science & Engineering 9, 90–95 (2007).

150. Liu, J. et al. An Integrated TCGA Pan-Cancer Clinical Data Resource to Drive High-Quality Survival Outcome Analytics. Cell 173, 400–416.e11 (2018).

151. McKinney, W. Data Structures for Statistical Computing in Python. SciPy 2010 10.25080/Majora-92bf1922-00a (2010) doi:10.25080/Majora-92bf1922-00a.

152. Davidson-Pilon, C. lifelines: survival analysis in Python. Journal of Open Source Software 4, 1317 (2019).

