## Supplementary figures and tables for "CD24 Acts as an Evolutionarily Conserved Innate Immune Checkpoint in Colorectal Cancer"

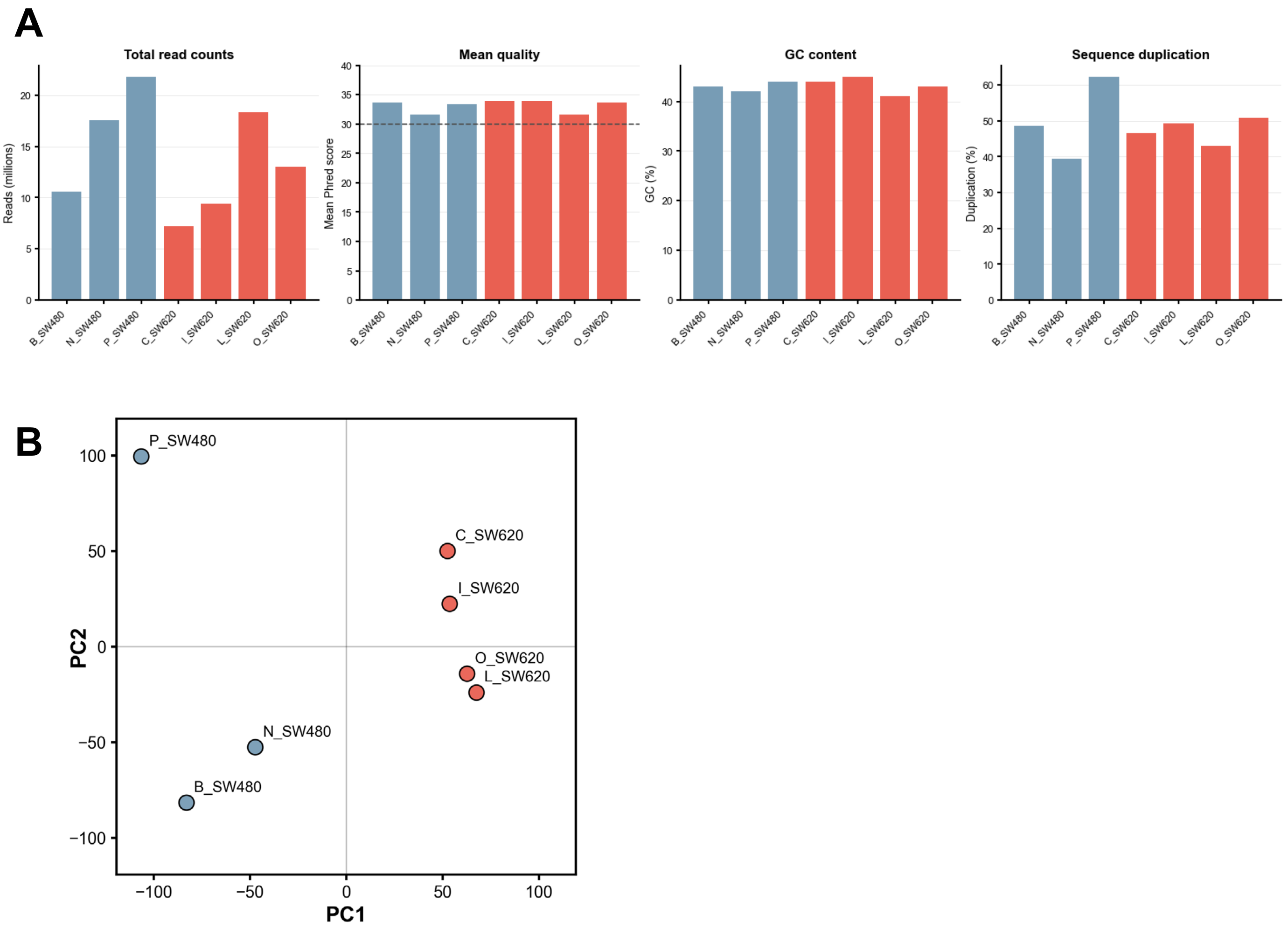


**Supplementary Figure 1 – Bulk RNAseq data analysis.**

**(A)** Sequencing quality control metrics for all bulk RNAseq samples, including total read counts, mean Phred quality score, GC content, and sequence duplication levels. **(B)** Principal component analysis (PCA) of bulk RNA-seq data was performed on voom-normalized expression values of human genes. Each point represents one biological replicate and colors indicate tumor origin (SW480, blue; SW620, red).

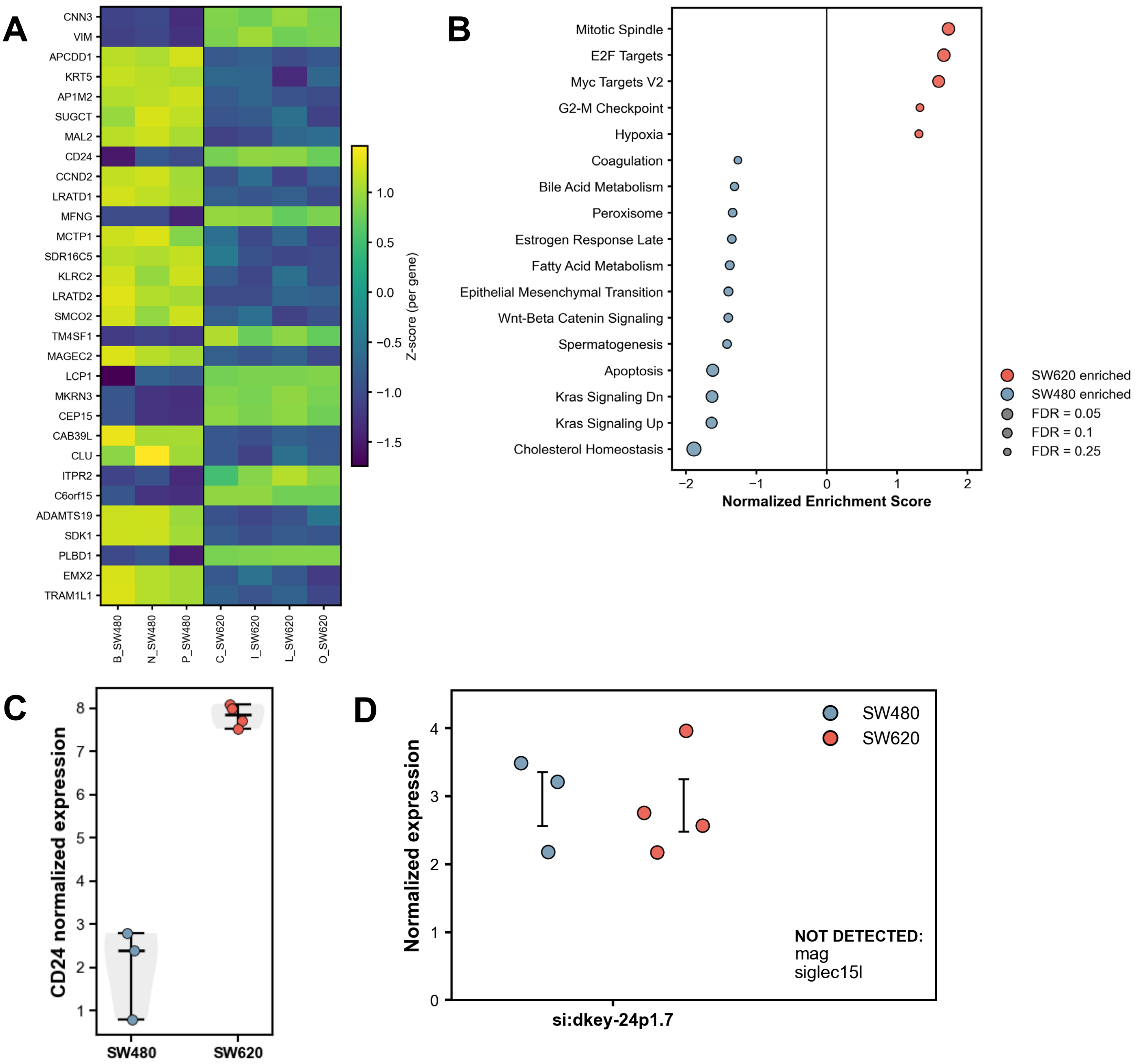


**Supplementary Figure 2 – Differential expression analysis of bulk RNAseq.**
**(A)** Heatmap of the top 30 differentially expressed human genes (FDR < 0.05, $|\log_2 FC| > 1$) between SW480 and SW620 tumors, shown as Z-scores per gene. **(B)** Hallmark gene set enrichment analysis (GSEA) performed on a pre-ranked list of all expressed human genes (ranked by limma t-statistic; SW620 vs SW480). Dot color indicates enrichment direction (SW620-enriched, red; SW480-enriched, blue) and dot size represents significance (−log10 FDR). **(C)** Normalized expression of CD24 in SW480 versus SW620. Each dot represents one sample. **(D)** Normalized expression of candidate CD24 receptor genes in the zebrafish component of SW480 and SW620 samples, showing detectable expression of si:dkey-24p1.7 but absence of mag and siglec15l. Each dot represents one sample.

#### Enrichment in SW480

| Pathway | Gene Set | NES | FDR (log10) |
|---|---|---|---|
| **Immune sensing & inflammation** | | | |
| Response to molecule of bacterial origin | GO | **-2.07** | **1.96** |
| Toll-like receptor signaling pathway | KEGG | **-1.75** | **1.74** |
| Cytosolic DNA-sensing pathway | KEGG | **-1.68** | **1.53** |
| NOD-like receptor signaling pathway | KEGG | **-1.62** | **1.40** |
| T Cell Receptor Signaling Pathway | WP | **-1.66** | **1.07** |
| Myeloid leukocyte differentiation | GO | **-1.74** | **0.98** |
| **Antigen processing / cellular stress** | | | |
| Proteasome | KEGG | **-2.34** | **6.00** |
| Spliceosome | KEGG | **-1.83** | **1.88** |
| Protein processing in endoplasmic reticulum | KEGG | **-1.78** | **1.83** |
| **Inflammatory metabolism & bioenergetic demand** | | | |
| Heme biosynthetic process | GO | **-2.40** | **6.00** |
| Oxidative phosphorylation | KEGG | **-2.27** | **6.00** |
| Electron Transport Chain | WP | **-2.09** | **2.75** |
| Iron-sulfur cluster assembly | GO | **-2.16** | **2.51** |

#### Enrichment in SW620

| Pathway | Gene Set | NES | FDR (log10) |
|---|---|---|---|
| **Metabolic insulation & lipid remodeling** | | | |
| Nuclear receptors in lipid metabolism and toxicity | WP | **+1.82** | **1.72** |
| Sphingolipid metabolism | KEGG | **+1.86** | **1.57** |
| PPAR signaling pathway | KEGG | **+1.61** | **1.18** |
| Fatty acid biosynthetic process | GO | **+1.57** | **0.86** |
| **Signaling & adhesion programs** | | | |
| Neuroactive ligand-receptor interaction | KEGG | **+2.13** | **2.63** |
| Cell adhesion molecules (CAMs) | KEGG | **+1.74** | **1.37** |
| Alpha6-Beta4 Integrin Signaling Pathway | WP | **+1.71** | **1.37** |
| Calcium signaling pathway | KEGG | **+1.75** | **1.32** |
| **Developmental plasticity / tolerance programs** | | | |
| Positive regulation of cell migration | GO | **+2.13** | **2.63** |
| Negative regulation of axon guidance | GO | **+2.07** | **2.25** |
| Regulation of BMP signaling pathway | GO | **+1.98** | **1.97** |

**Supplementary Figure 3 – Gene set enrichment analysis of the zebrafish transcriptome of bulk RNAseq samples.**

Gene set enrichment analysis (GSEA) of transcriptomic profiles from zebrafish xenografts, comparing SW620 and SW480 tumors. Pathways are grouped into manually curated biological themes and displayed by enrichment direction (left: SW480-enriched, NES < 0; right: SW620-enriched, NES > 0). For each pathway, the gene set source, normalized enrichment score (NES), and significance (FDR, shown as −log10(FDR), capped at 6) are indicated.

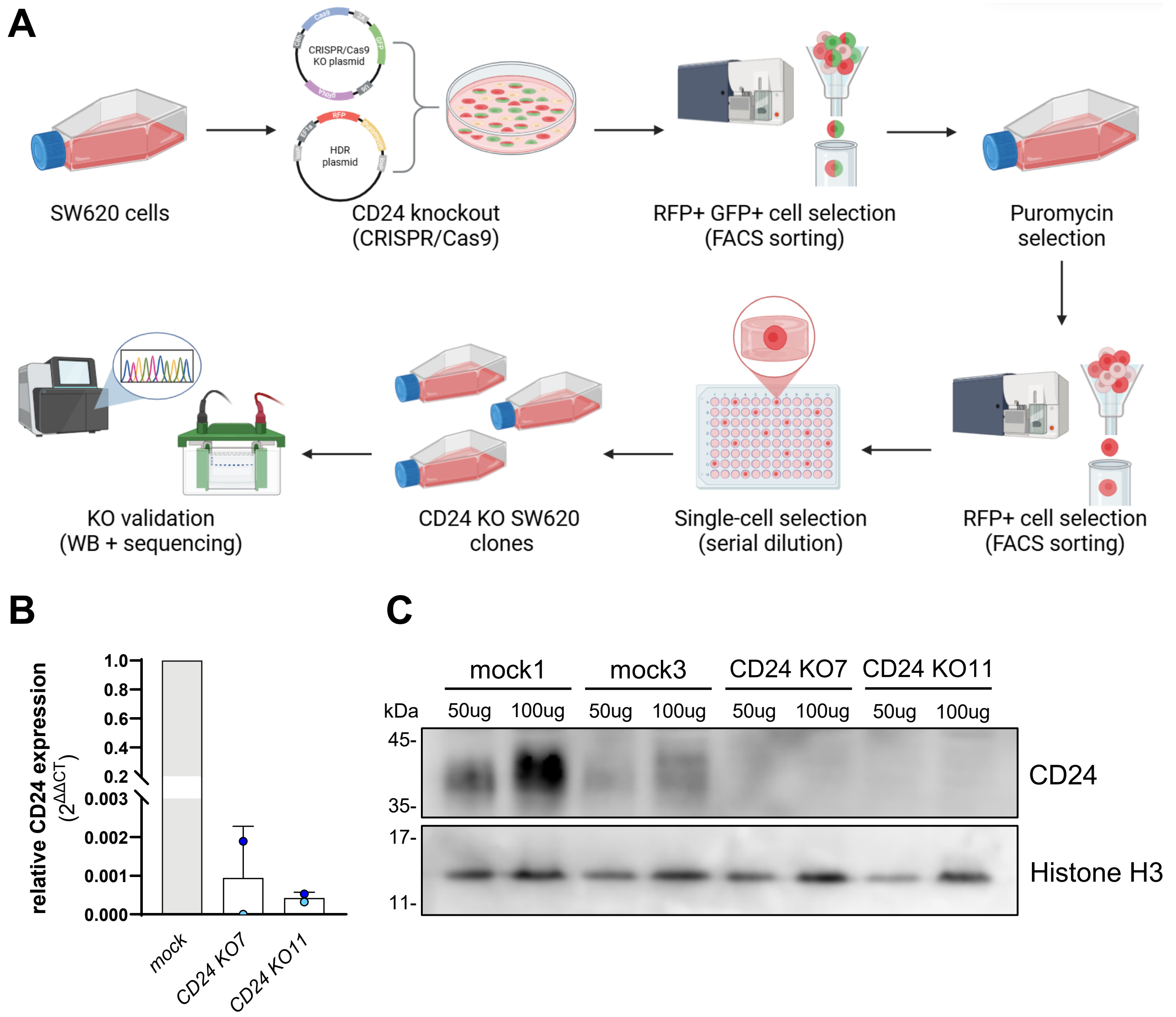


**Supplementary Figure 4 - CD24 gene knockout in SW620 cells.**
**(A)** Schematic representation of *CD24* knockout SW620 cell line generation using CRISPR/Cas9 technology. Image created using BioRender. **(B)** Quantitative real-time PCR (qPCR) analysis of *CD24* mRNA expression in mock and KO single-cell-derived clones, normalized to housekeeping gene EEF1A1. ΔΔCT was calculated as ΔCt of each KO clone minus ΔCt of the average of the mock 1 and mock 3 clones. The graph shows mean ± S.D. Each dot represents one independent experiment. Data (ΔCT values) were analyzed using the Friedman test. **(C)** Western blot analysis of *CD24* protein expression in SW620 mock and knockout (KO) single-cell-derived clones. Histone H3 was used as a loading control. Two independent single-cell clones were validated for both the mock (1 and 3) and CD24 KO (7 and 11) conditions. For mock1 and CD24 KO11 clone, two independently collected samples were tested (n=2).

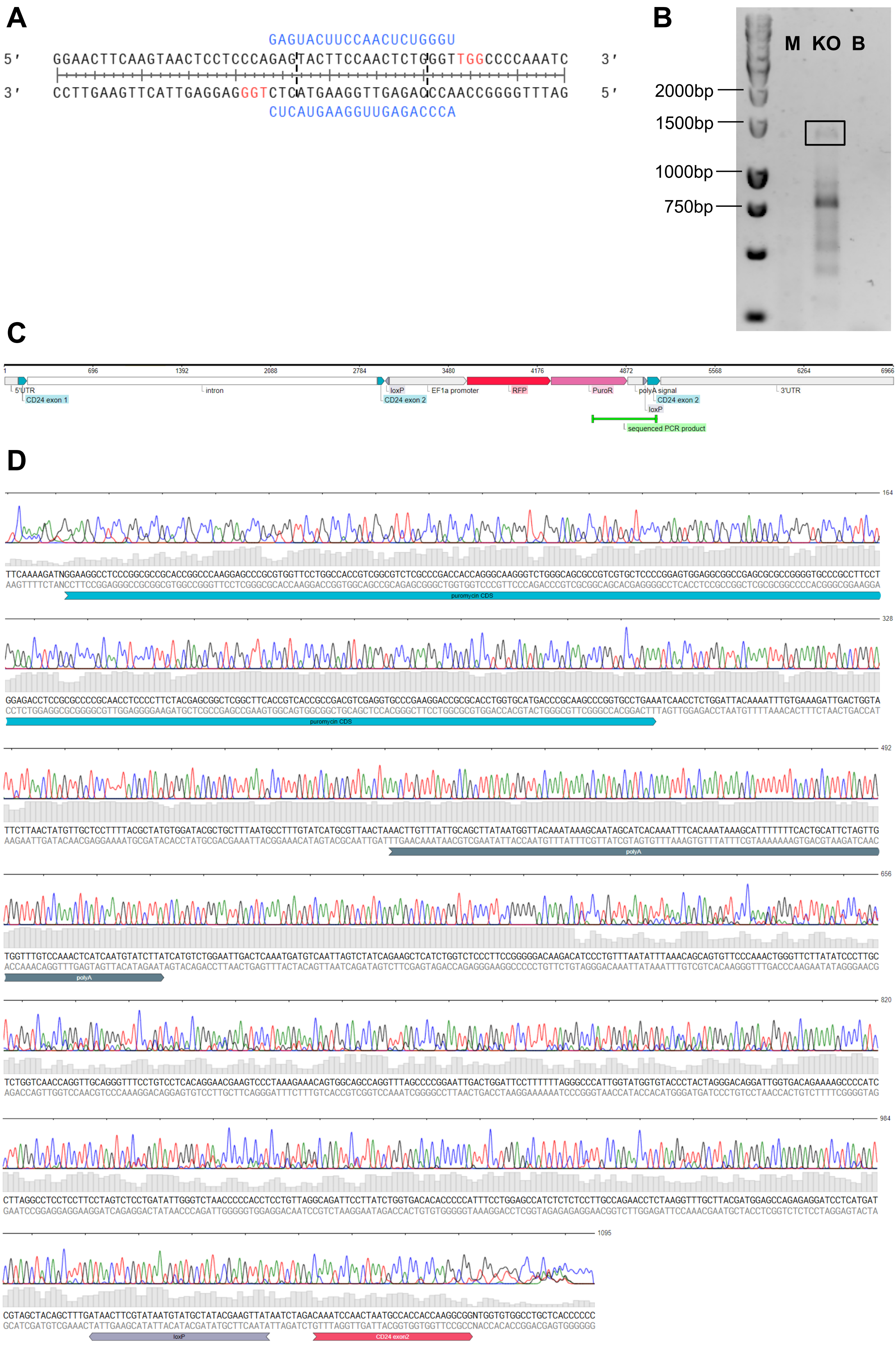

A
GAGUACUUCCAACUCUGGGU
5′ GGAACTTCAAGTAACTCCTCCCAGAGTACTTCCAACTCTGGGTTGGCCCCAAATC 3′
3′ CCTTGAAGTTCATTGAGGAGGGTCTCATGAAGGTTGAGACCCAACCGGGGTTTAG 5′
CUCAUGAAGGUUGAGACCCA
B
M KO B
2000bp
1500bp
1000bp
750bp
C
1
696
1392
2088
2784
3480
4176
4872
5568
6264
6966
5'UTR
CD24 exon 1
intron
loxP
CD24 exon 2
EF1a promoter
RFP
PuroR
polyA signal
CD24 exon 2
loxP
3'UTR
sequenced PCR product
D
164
puromycin CDS
328
puromycin CDS
492
polyA
656
polyA
820
984
1095
loxP
CD24 exon2

**Supplementary Figure 5 - Gene editing characterization of SW620 CD24 KO11 clone.**
**(A)** Sequence comparison of the *CD24* gene with the gRNA used for CRISPR/Cas9, targeting exon 2. The gRNA and PAM sequences are shown in blue and red, respectively. The potential cleavage sites of the gRNAs are indicated by dashed lines. **(B)** The genomic DNA fragment of *CD24* KO11 (KO) SW620 single-cell-derived clone was amplified through PCR (indicated by the black rectangle) to confirm the presence of the construct in the CD24 gene after recombination. The primers were selected to bind to the puromycin-resistant gene and the CD24 exon 2, resulting in a 1095 bp PCR product. SW620 mock 1 (M) and the negative control ($H_2O$, B) showed no amplification, as expected. **(C)** Schematic representation of the edited *CD24* gene in the SW620 CD24 KO11 single-cell-derived clone. It shows the HDR insert location, containing the RFP and Puromycin resistance (PuroR) genes, as well as the sequenced amplicon. Created using VectorBee. **(D)** Sanger sequencing chromatogram of the amplified DNA amplicon, showing that it contains the end of the PuroR gene and the beginning of CD24 exon 2, as expected. Created using VectorBee.

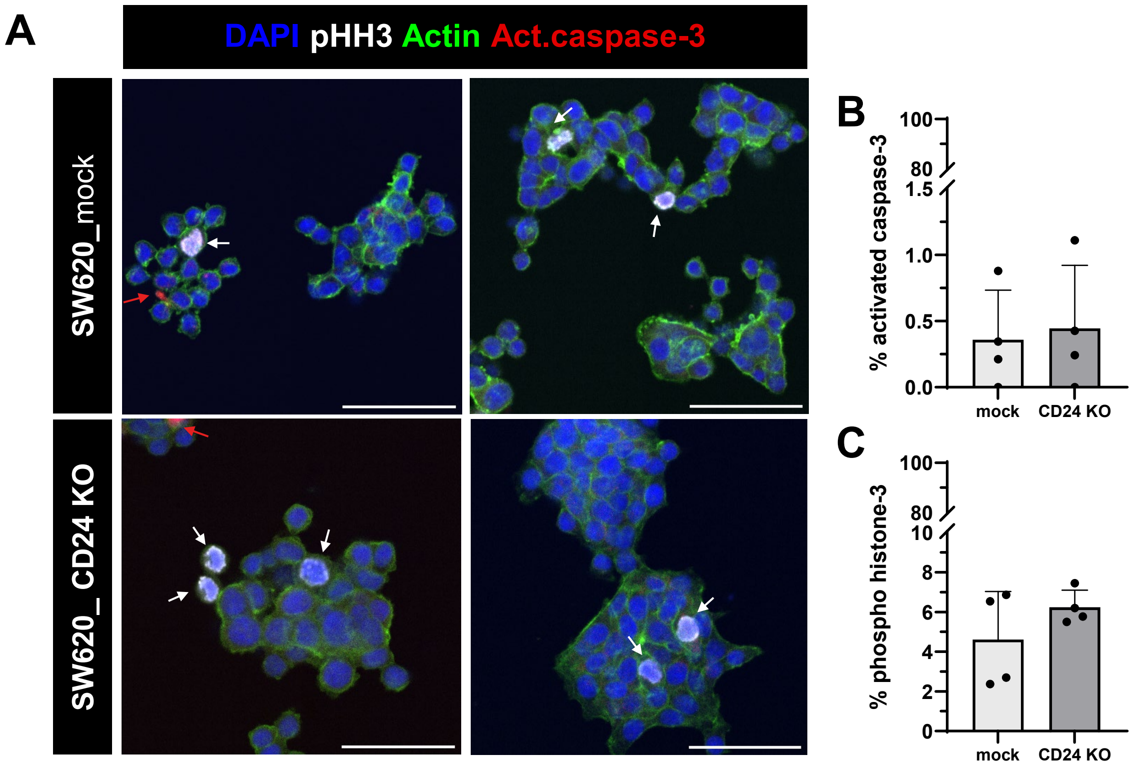


**Supplementary Figure 6 – CD24 KO has no effect on SW620 viability or proliferation rate *in vitro*.**

**(A)** Representative confocal images of SW620 mock clone 1 and CD24 KO clone 11 cells stained for actin filaments (phalloidin, in green), apoptosis (activated caspase-3, in red), proliferation (phospho-histone H3, in white), and nuclei (DAPI, in blue). Scale bars: 50 µm. (**B**) Quantification of the percentage of apoptotic cells (activated caspase-3 positive). (**C**) Quantification of the percentage of dividing cells (phospho-histone H3 positive). Graphs show mean ± S.D., and each dot represents the average of 10 random fields of view of one coverslip. Data was analyzed using the Mann-Whitney U test ($p > 0.05$).

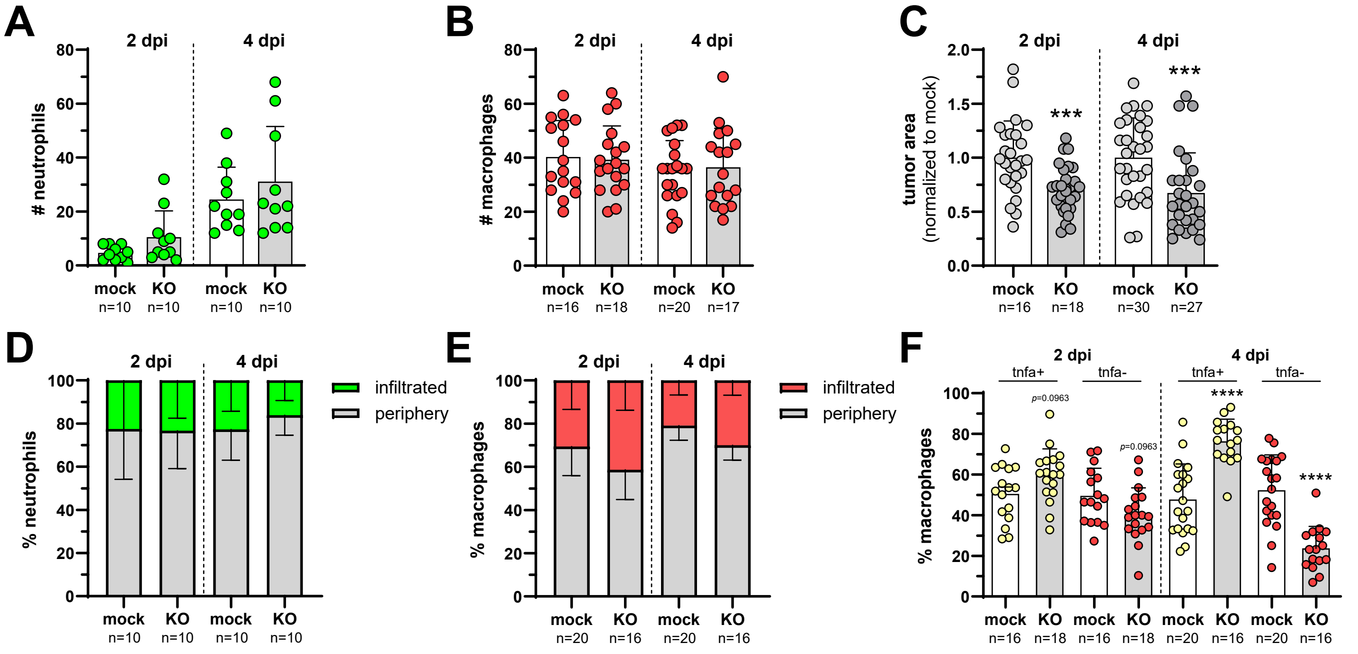


**Supplementary Figure 7 – Innate immune analysis of SW620 mock and CD24 KO xenografts.**

**(A,B)** Quantification of the total number of neutrophils (A) and macrophages (B) in SW620 mock 1 and CD24 KO 11, at 2 and 4 dpi. **(C)** Quantification of tumor area, normalized to mock. **(D,E)** Quantification of the percentage of neutrophils (D) and macrophages (E) in the tumor periphery versus infiltrated inside the tumor. **(F)** Quantification of the percentage of Tnfa-expressing macrophages and Tnfa-negative macrophages. All graphs show mean ± S.D. Each dot represents one zebrafish xenograft and the total number analyzed (n) is also depicted. According to data normality, the Welch's t test or the Mann-Whitney U test were used to compare mock to CD24 (ns > 0.05, ***P ≤ 0.001, and ****P ≤ 0.0001).

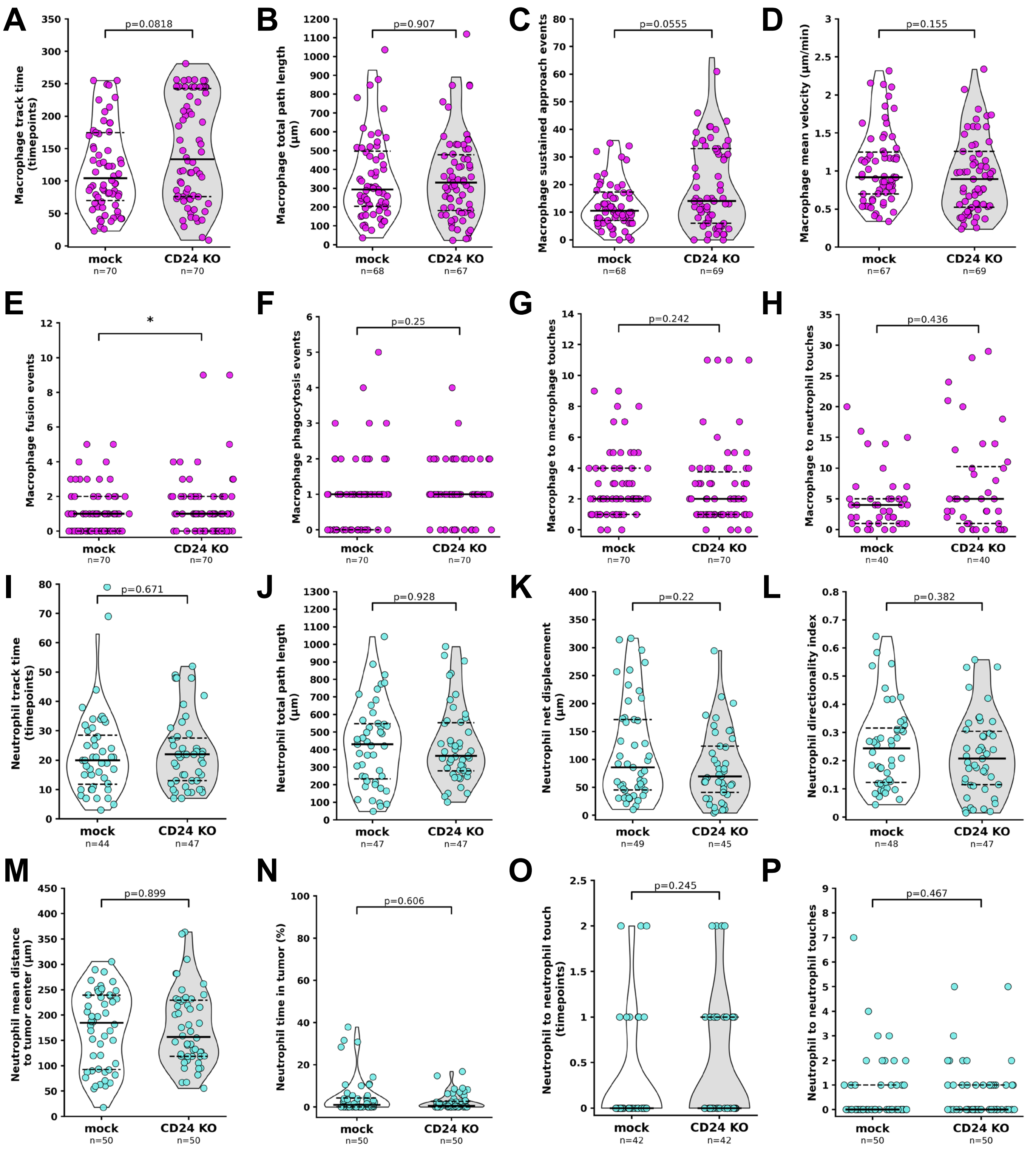
A
Macrophage track time (timepoints)
p=0.0818
mock n=70
CD24 KO n=70
B
Macrophage total path length (µm)
p=0.907
mock n=68
CD24 KO n=67
C
Macrophage sustained approach events
p=0.0555
mock n=68
CD24 KO n=69
D
Macrophage mean velocity (µm/min)
p=0.155
mock n=67
CD24 KO n=69
E
Macrophage fusion events
*
mock n=70
CD24 KO n=70
F
Macrophage phagocytosis events
p=0.25
mock n=70
CD24 KO n=70
G
Macrophage to macrophage touches
p=0.242
mock n=70
CD24 KO n=70
H
Macrophage to neutrophil touches
p=0.436
mock n=40
CD24 KO n=40
I
Neutrophil track time (timepoints)
p=0.671
mock n=44
CD24 KO n=47
J
Neutrophil total path length (µm)
p=0.928
mock n=47
CD24 KO n=47
K
Neutrophil net displacement (µm)
p=0.22
mock n=49
CD24 KO n=45
L
Neutrophil directionality index
p=0.382
mock n=48
CD24 KO n=47
M
Neutrophil mean distance to tumor center (µm)
p=0.899
mock n=50
CD24 KO n=50
N
Neutrophil time in tumor (%)
p=0.606
mock n=50
CD24 KO n=50
O
Neutrophil to neutrophil touch (timepoints)
p=0.245
mock n=42
CD24 KO n=42
P
Neutrophil to neutrophil touches
p=0.467
mock n=50
CD24 KO n=50

**Supplementary Figure 8 – Extended live imaging analysis of myeloid behavior in SW620 mock and CD24 KO xenografts.**

**(A–H)** Extended macrophage motility and interaction metrics. Panels show total track length in terms of time (timepoints, A) and distance (µm, B), number of sustained approach events towards the tumor (C), mean velocity (D), number of fusion events (E), number of phagocytosis events (F), number of macrophage-macrophage contact events (G), and number of macrophage-neutrophil contact events (H). **(I–P)** Extended neutrophil motility and interaction metrics. Panels show total track length in terms of time (timepoints, I) and distance (µm, J), net displacement (K), directionality index (L), mean distance to the tumor center (M), percentage of time in the tumor (N), number of timepoints in contact with other neutrophils (O), and number of neutrophil-neutrophil contact events (P). Each dot represents a single cell track, violins depict the distribution with median and interquartile range overlaid, and the total number of tracks analyzed (n) is indicated. Data are pooled from 6 xenografts per condition across 3 independent experiments. Depending on data normality, Welch's t test or Mann-Whitney U test were used to compare mock and CD24 KO conditions (*$P \leq 0.05$).

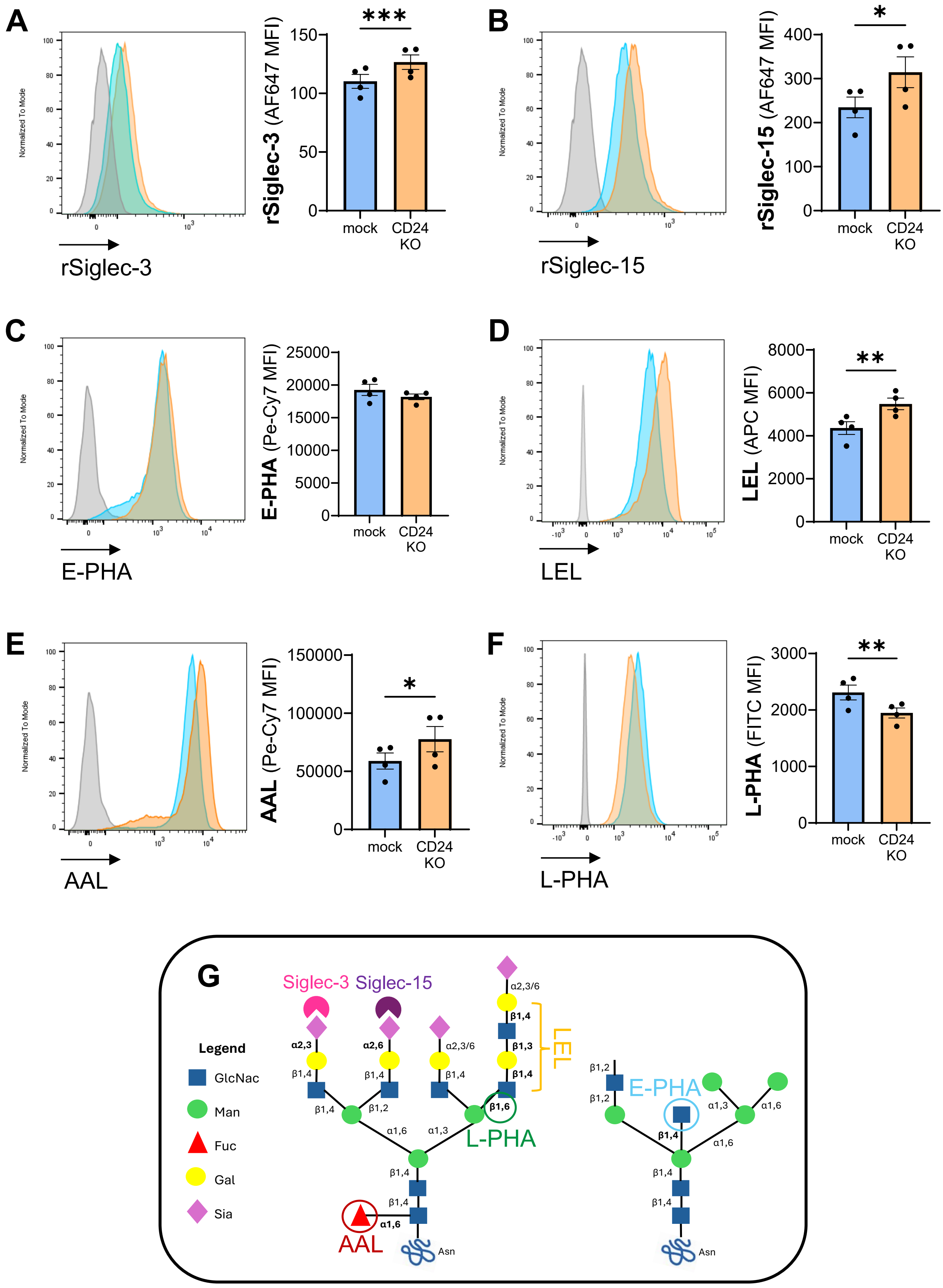
A
rSiglec-3
rSiglec-3 (AF647 MFI)
***
mock
CD24 KO
B
rSiglec-15
rSiglec-15 (AF647 MFI)
*
C
E-PHA
E-PHA (Pe-Cy7 MFI)
D
LEL
LEL (APC MFI)
**
E
AAL
AAL (Pe-Cy7 MFI)
F
L-PHA
L-PHA (FITC MFI)
G
Legend
GlcNac
Man
Fuc
Gal
Sia
Siglec-3
Siglec-15
LEL
L-PHA
E-PHA
AAL
Asn

**Supplementary Figure 9 – Glycocalyx analysis of SW620 cells.**
Flow cytometry was used to analyze Siglec and lectin binding on SW620 mock and CD24 KO cells. Staining histograms and median fluorescence intensity (MFI) plots are shown. Siglec-3 (**A**) and Siglec-15 (**B**) binding was quantified using recombinant proteins, while bisecting-GlcNAc *N*-glycans (**C**, via E-PHA), polylactosamine structures (**D**, via LEL), α-1,6 linked fucose to the core *N*-acetylglucosamine (**E**, via AAL), and β1-6-GlcNAc branched *N*-glycans **(F,** via L-PHA) were bound by lectins. The graphs show mean ± S.E.M. and each dot represents one replicate. Data normality was analyzed using the Shapiro-Wilk test, and conditions were compared accordingly using a paired t-test or a Wilcoxon test (*P<0.05, **P<0.01, ***P<0.001). **(G)** Schematic representation of the glycan epitopes recognized by Siglec-3, Siglec-15, E-PHA, LEL, AAL, and L-PHA, shown on a representative tetra-antennary N-glycan structure.

**A**

| Zebrafish protein | E-value | | Identity percentage | | Query cover | | Described human homology |
|---|---|---|---|---|---|---|---|
| | IgV domain | Whole protein | IgV domain | Whole protein | IgV domain | Whole protein | |
| mag | 2e-13 | 3e-54 | 30.34% | 23.47% | 83% | 95% | MAG/SIGLEC4 |
| siglec15l | 2e-09 | 3e-07 | 29.47% | 29.47% | 84% | 38% | SIGLEC15 |
| si:dkey-24p1.7 | 2e-08 | 4e-14 | 25.26% | 19.69% | 89% | 100% | CD33 multiple SIGLECs |

**B**

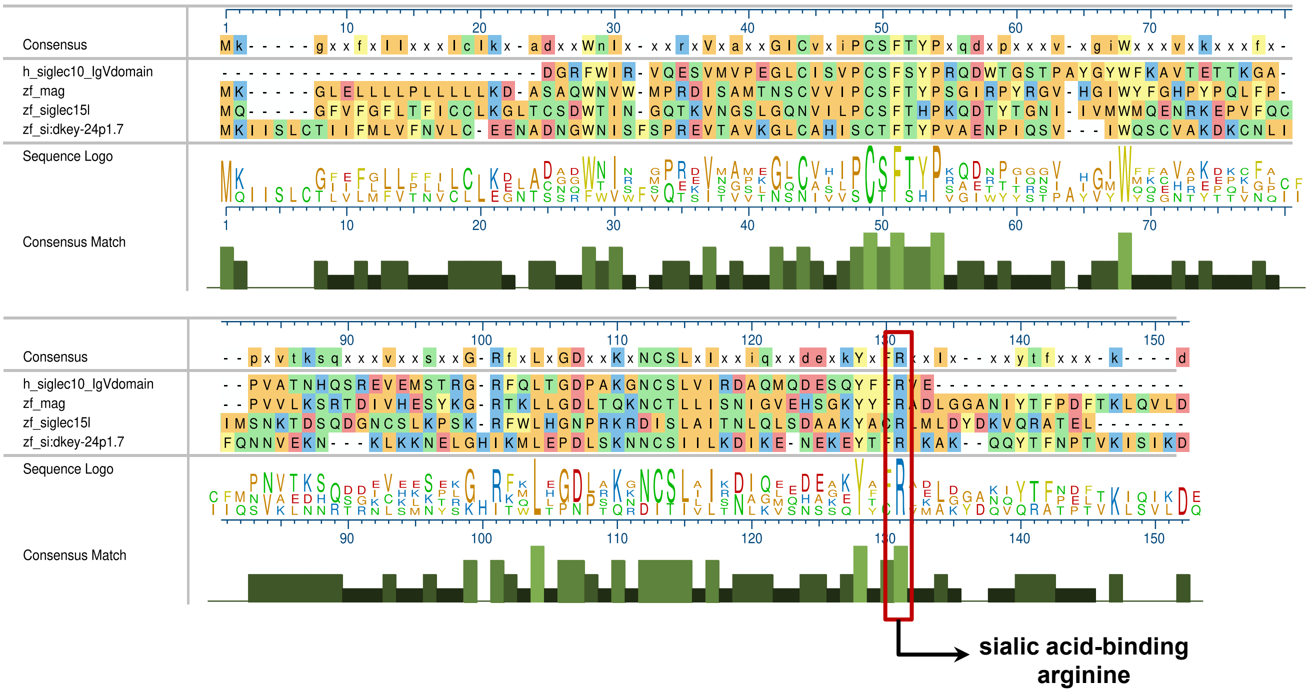


**C**

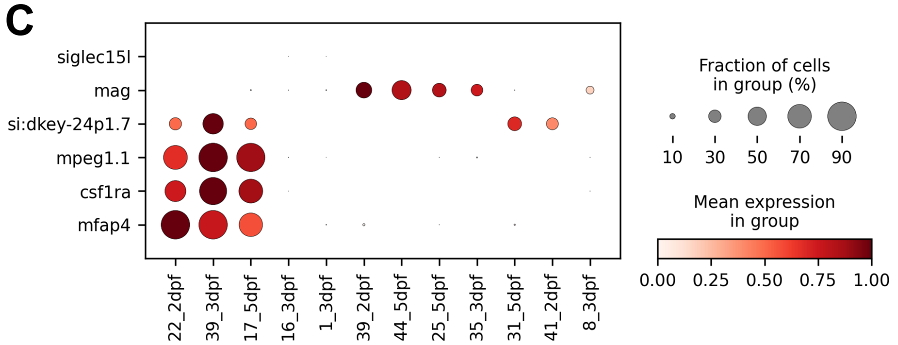


**D**

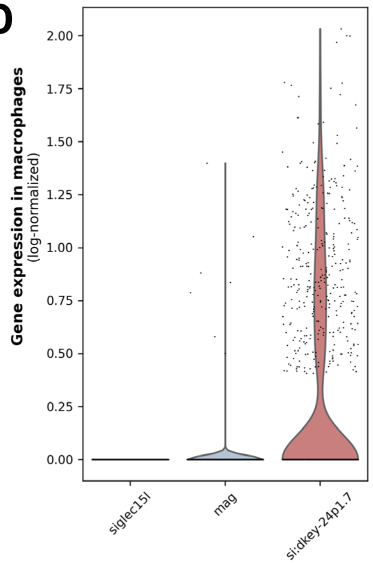

**Supplementary Figure 10 - Identification of candidate zebrafish CD24-binding proteins.** **(A)** Summary of BLASTp results comparing human SIGLEC10 (IgV domain and full-length sequence) with the three putative zebrafish homologs (*siglec15l*, *mag*, and si:dkey-24p1.7). E-values, identity, and query coverage are shown for each comparison. Orthology relationships were evaluated using the HGNC Comparison of Orthology Predictions (HCOP) tool. **(B)** Multiple sequence alignment of the IgV (V-set) domain from human SIGLEC10 and the zebrafish homolog candidates. The alignment was performed using the EMBL-EBI Multiple Sequence Alignment tool (Clustal Omega) and visualized in MegAlign Pro. Amino acids are color-coded by chemistry, and residue consensus and conservation are also displayed. The sialic acid–binding arginine (R119 in human SIGLEC10) is fully conserved and highlighted in red. **(C)** Dot plot showing scaled expression of the three candidate orthologs in zebrafish macrophages. Single-cell RNA-seq data from zebrafish larvae at 2, 3, and 5 dpf, extracted from Zebrahub, were analyzed to identify macrophage-enriched clusters at each developmental timepoint (22_2dpf, 39_3dpf, 17_5dpf), using cluster annotation and the expression of established macrophage markers (*mpeg1.1*, *csf1ra*, *mfap4*). A selection of additional non-macrophage clusters showing detectable expression of candidate genes is also included. Dot size indicates the proportion of cells expressing each gene, and color intensity represents the average standardized (z-scored) expression. **(D)** Violin plot showing the expression of the three candidate homologs in zebrafish macrophages across all timepoints (2, 3, and 5 dpf). Violin width shows the distribution of log-normalized gene expression across cells within each cluster, and individual points represent single-cell values.

Movies 1–10 have been deposited in the BioImage Archive under accession S-BIAD3811.

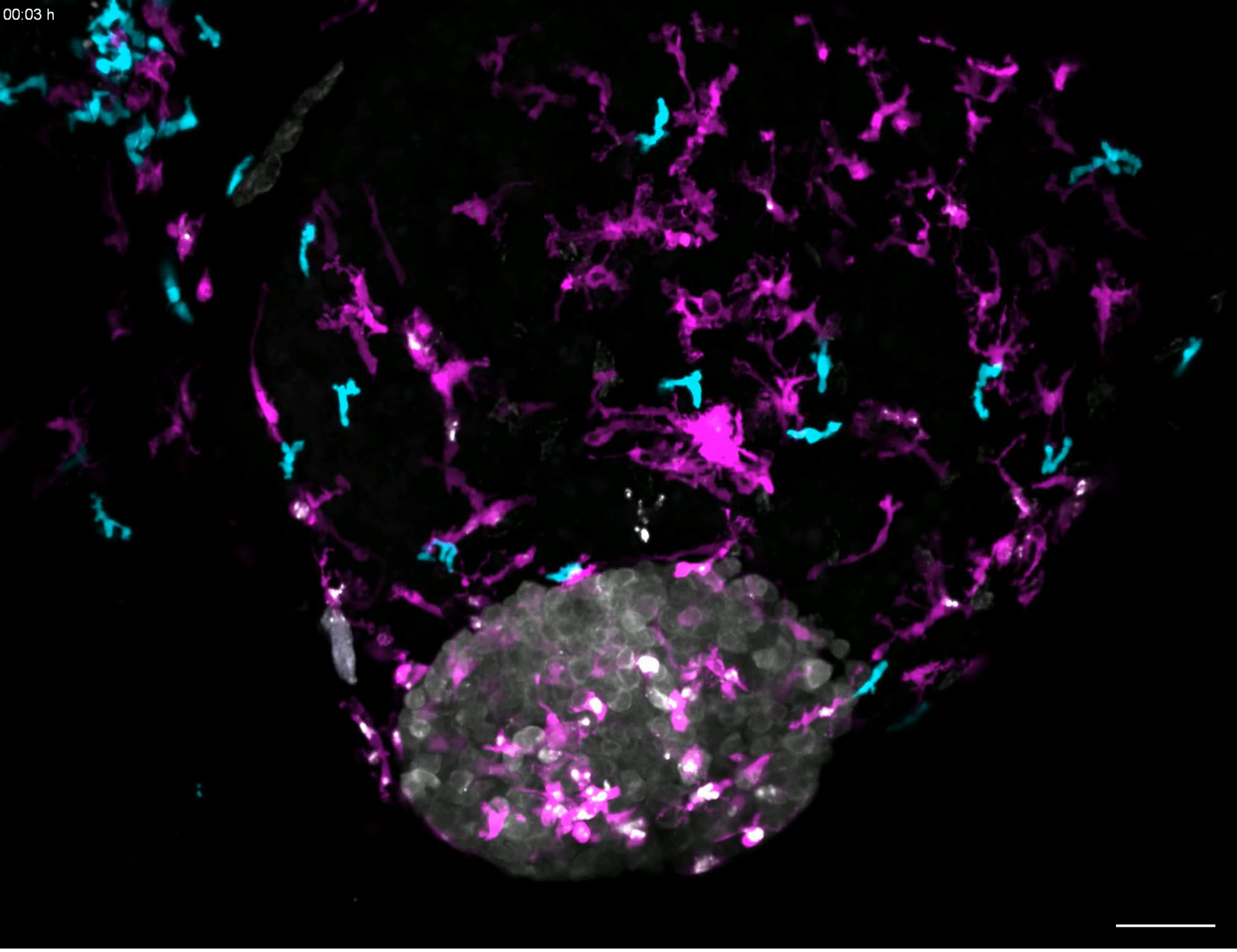


**Movie 1 – Representative time-lapse of macrophage and neutrophil kinetics in a mock tumor.** SW620 mock xenograft generated in Tg(mpeg1:mCherry-F, mpx:GFP-F) larvae showing tumor cells (gray), macrophages (magenta), and neutrophils (cyan). The tumor was acquired overnight between 1 and 2 dpi, in stacks with 3-µm intervals every 3 minutes, and is shown as a maximum projection. Scale bar: 50 µm. Available from the BioImage Archive (accession S-BIAD3811).

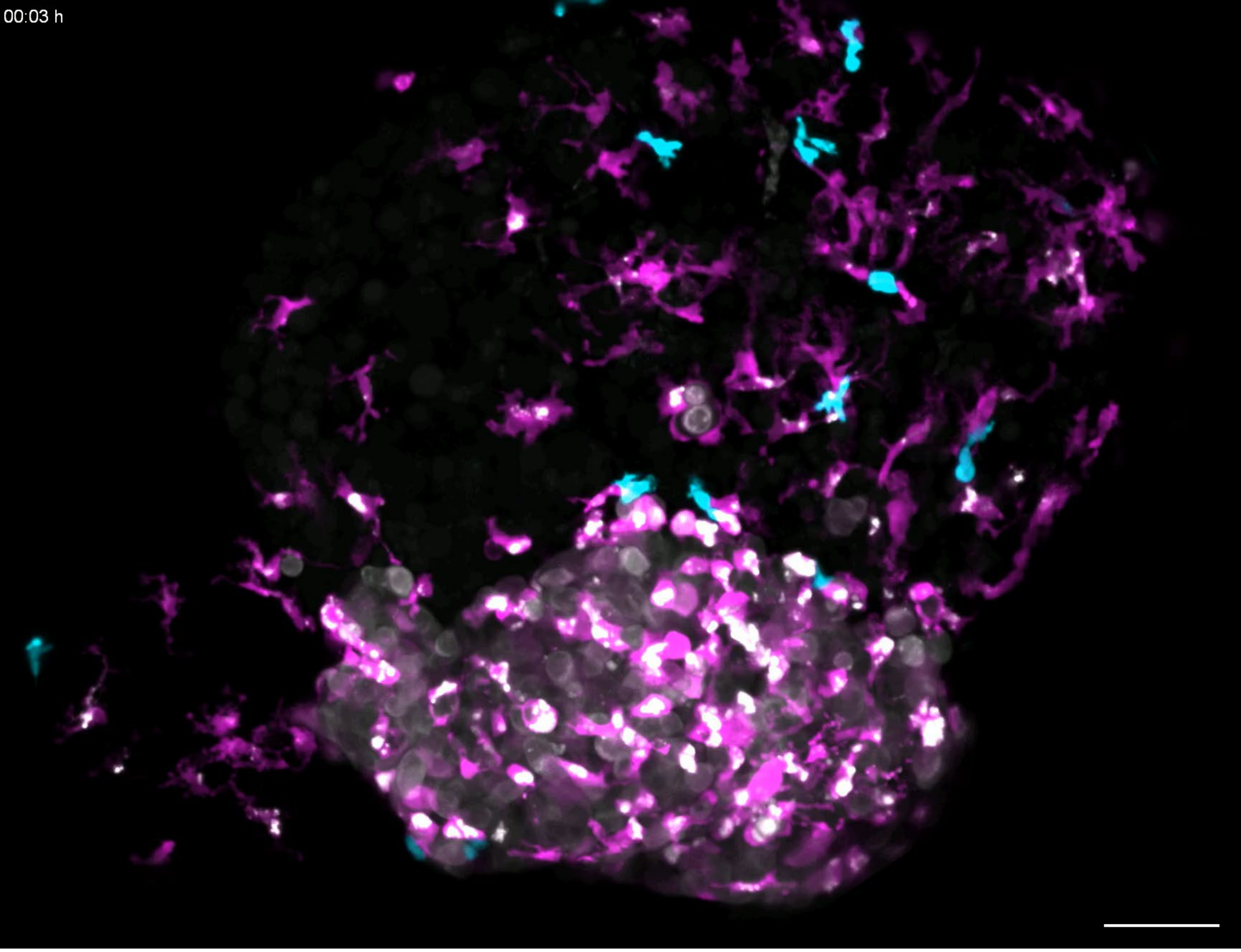


**Movie 2 – Representative time-lapse of macrophage and neutrophil kinetics in a CD24 KO tumor.** SW620 CD24 KO xenograft generated in Tg(mpeg1:mCherry-F, mpx:GFP-F) larvae showing tumor cells (gray), macrophages (magenta), and neutrophils (cyan). The tumor was acquired overnight between 1 and 2 dpi, in stacks with 3-μm intervals every 3 minutes, and is shown as a maximum projection. Scale bar: 50 μm. Available from the BioImage Archive (accession S-BIAD3811).

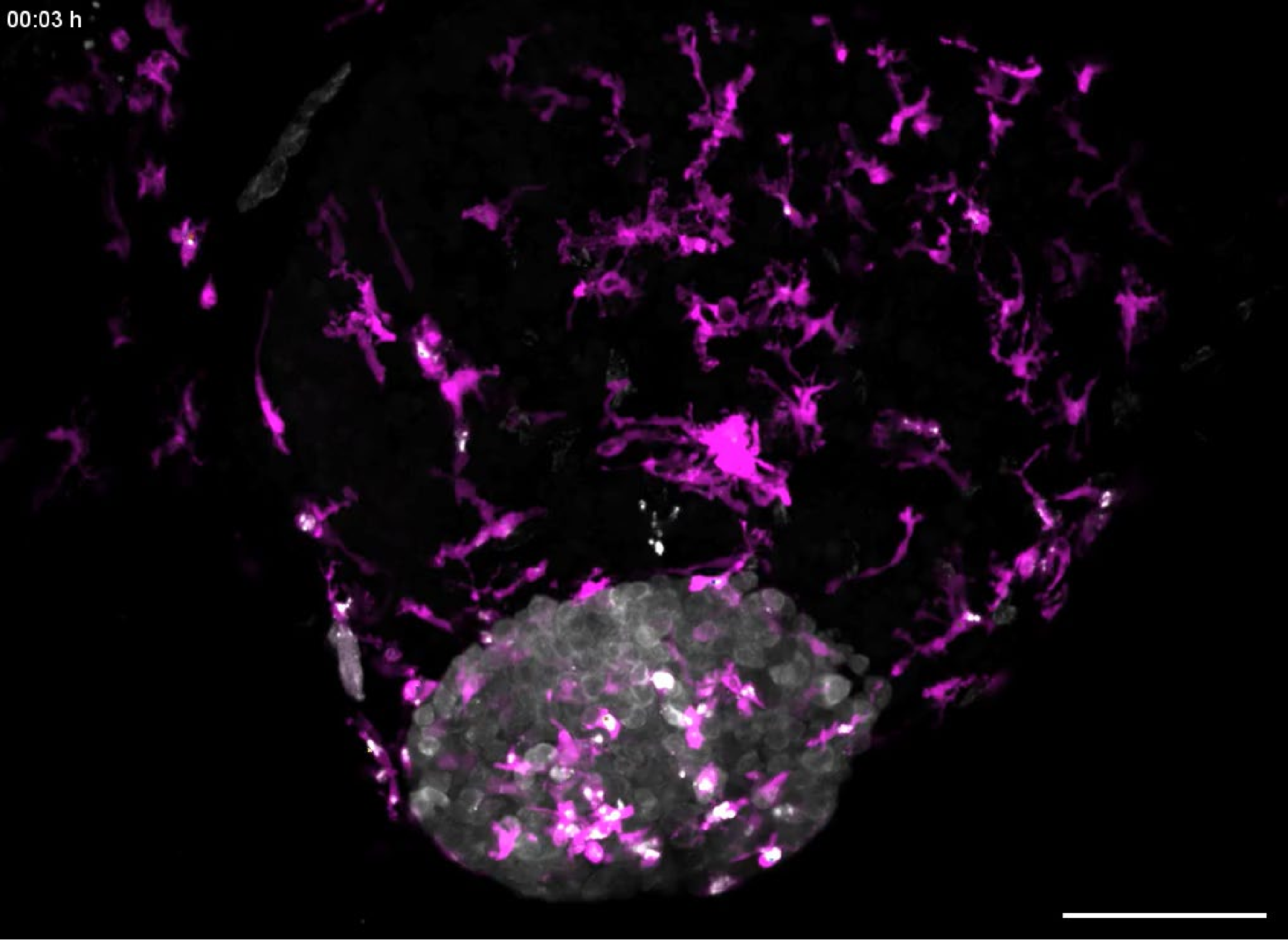


**Movie 3 – Representative time-lapse of macrophage cell tracks in a mock tumor.** SW620 mock xenograft generated in Tg(mpeg1:mCherry-F) larvae showing tumor cells (gray) and macrophages (magenta). The tumor was acquired in stacks with 3-μm intervals every 3 minutes and is shown as a maximum projection. Each cell track is shown in a different color. Scale bar: 50 μm. Available from the BioImage Archive (accession S-BIAD3811).

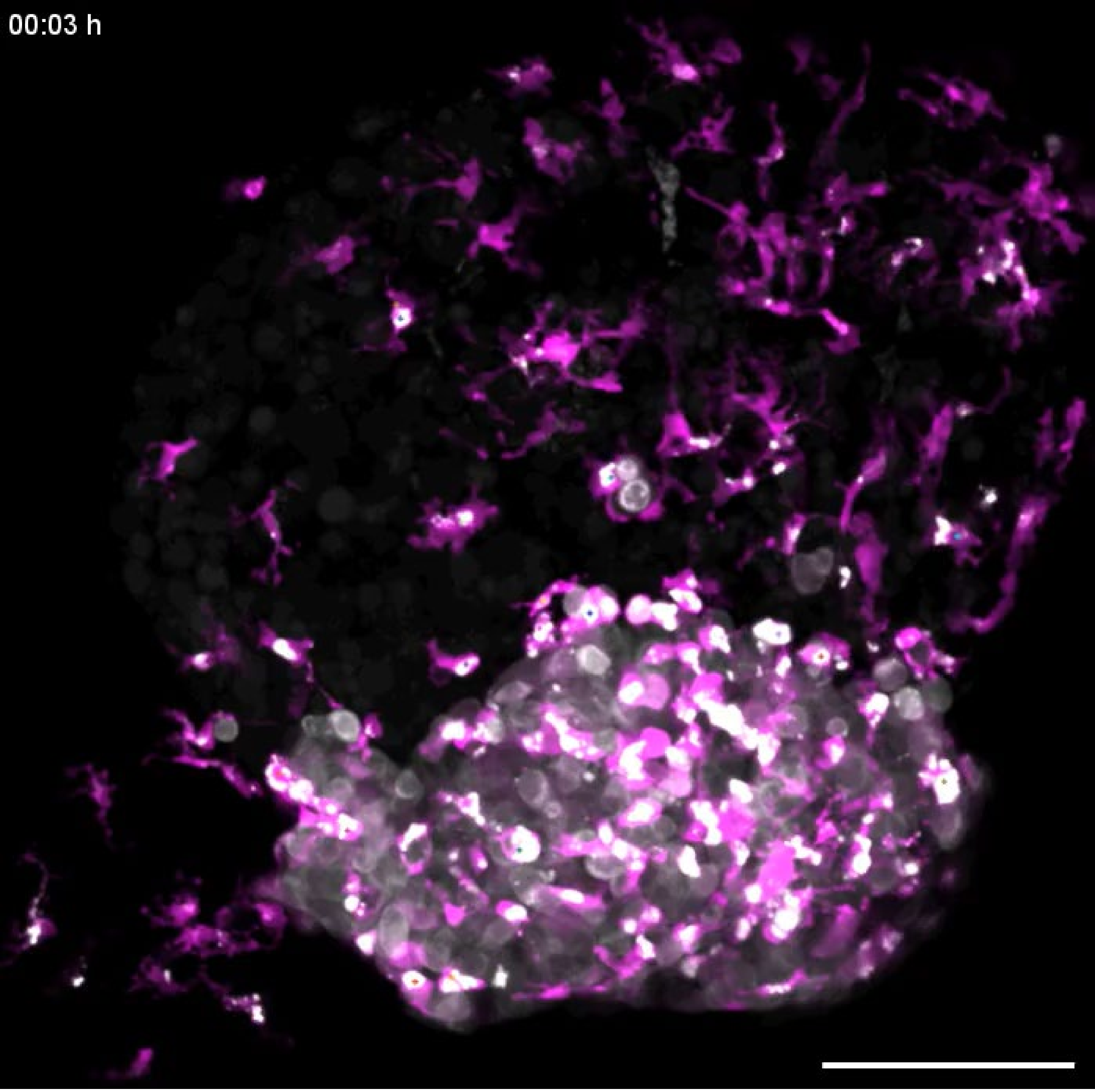


**Movie 4 – Representative time-lapse of macrophage cell tracks in a CD24 KO tumor.** SW620 CD24 KO xenograft generated in Tg(mpeg1:mCherry-F) larvae showing tumor cells (gray) and macrophages (magenta). The tumor was acquired in stacks with 3-μm intervals every 3 minutes and is shown as a maximum projection. Each cell track is shown in a different color. Scale bar: 50 μm. Available from the BioImage Archive (accession S-BIAD3811).

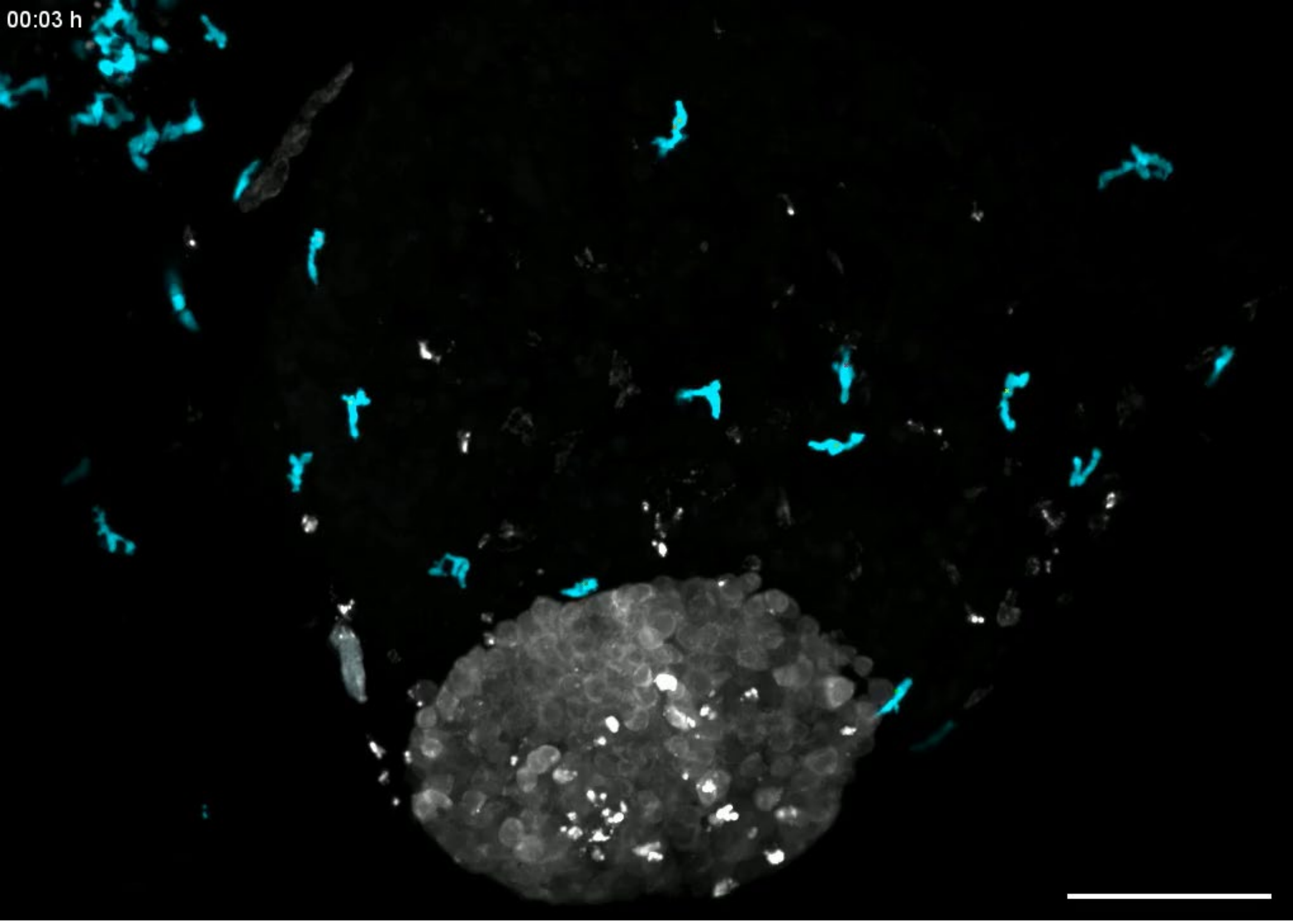


**Movie 5 – Representative time-lapse of neutrophil cell tracks in a mock tumor.** SW620 mock xenograft generated in Tg(mpx:GFP-F) larvae showing tumor cells (gray) and neutrophils (cyan). The tumor was acquired in stacks with 3-μm intervals every 3 minutes and is shown as a maximum projection. Each cell track is shown in a different color. Scale bar: 50 μm. Available from the BioImage Archive (accession S-BIAD3811).

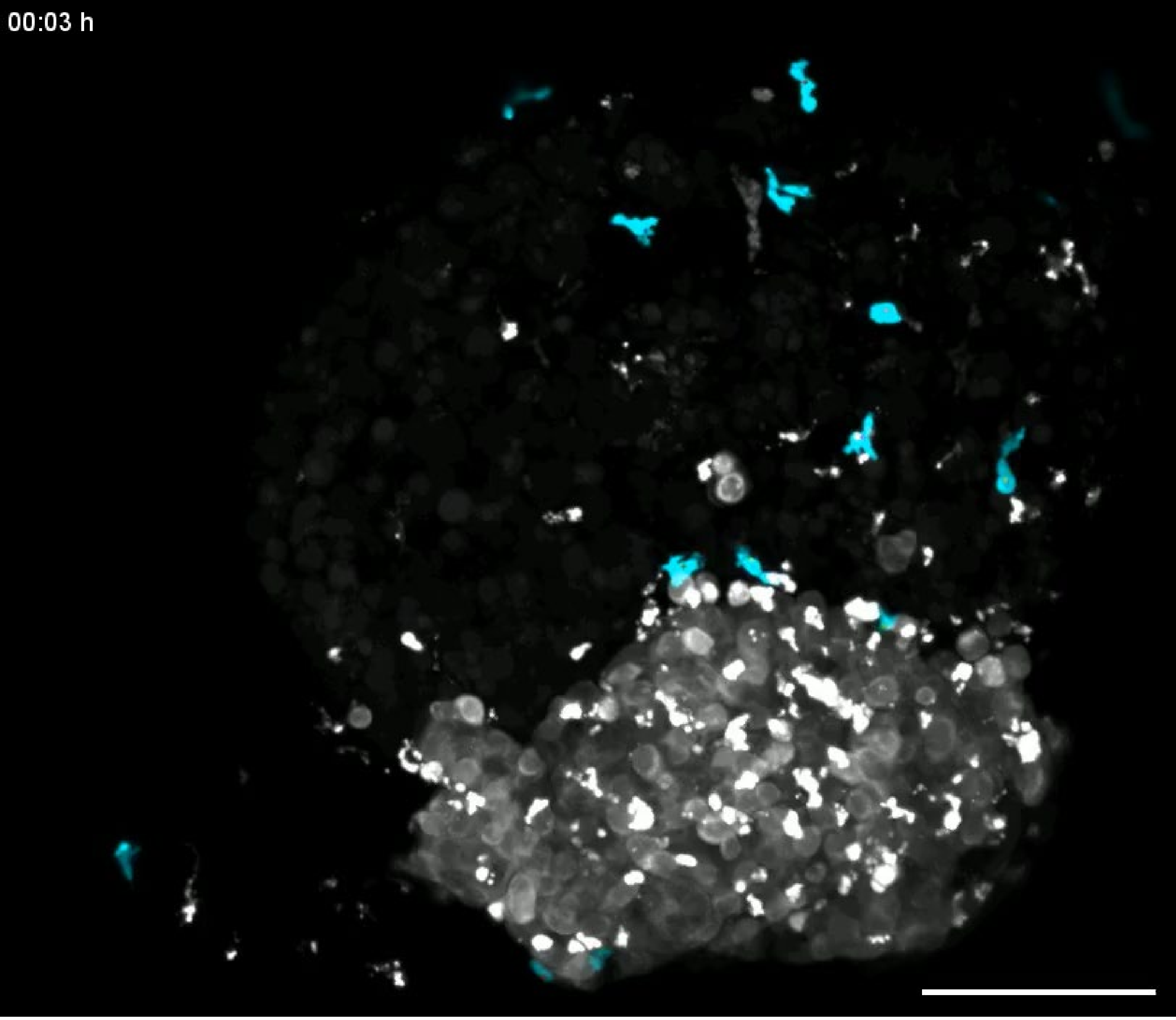


**Movie 6 – Representative time-lapse of neutrophil cell tracks in a CD24 KO tumor.** SW620 CD24 KO xenograft generated in Tg(mpx:GFP-F) larvae showing tumor cells (gray) and neutrophils (cyan). The tumor was acquired in stacks with 3-μm intervals every 3 minutes and is shown as a maximum projection. Each cell track is shown in a different color. Scale bar: 50 μm. Available from the BioImage Archive (accession S-BIAD3811).

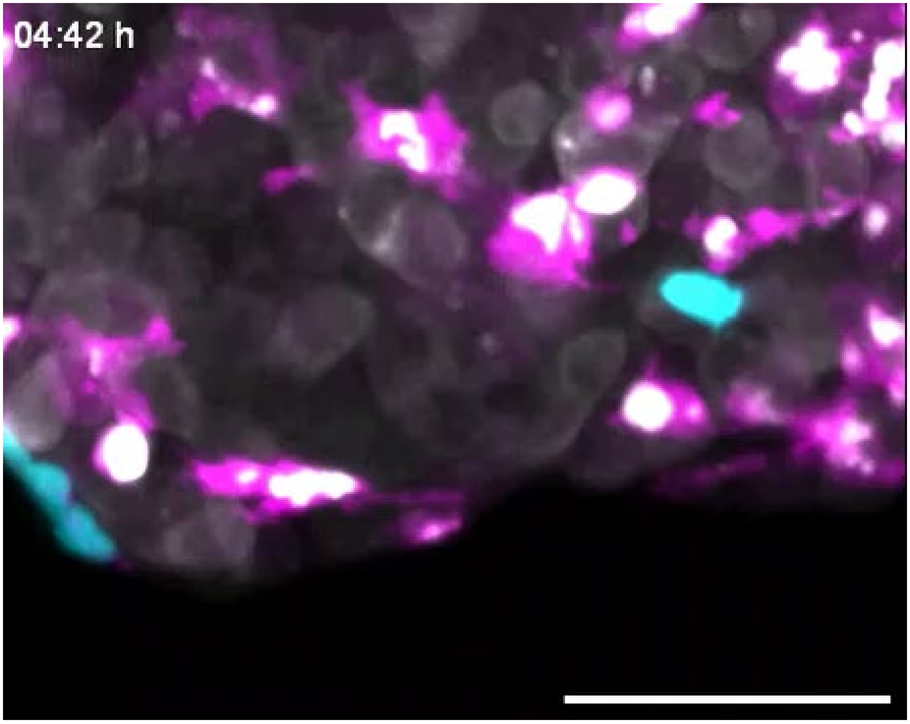


**Movie 7 – Representative time-lapse of a macrophage fusion event.** SW620 xenograft generated in Tg(mpeg1:mCherry-F, mpx:GFP-F) larvae showing tumor cells (gray), macrophages (magenta), and neutrophils (cyan). The tumor was acquired overnight between 1 and 2 dpi, in stacks with 3-µm intervals every 3 minutes, and is shown as a maximum projection. Scale bar: 50 µm. Available from the BioImage Archive (accession S-BIAD3811).

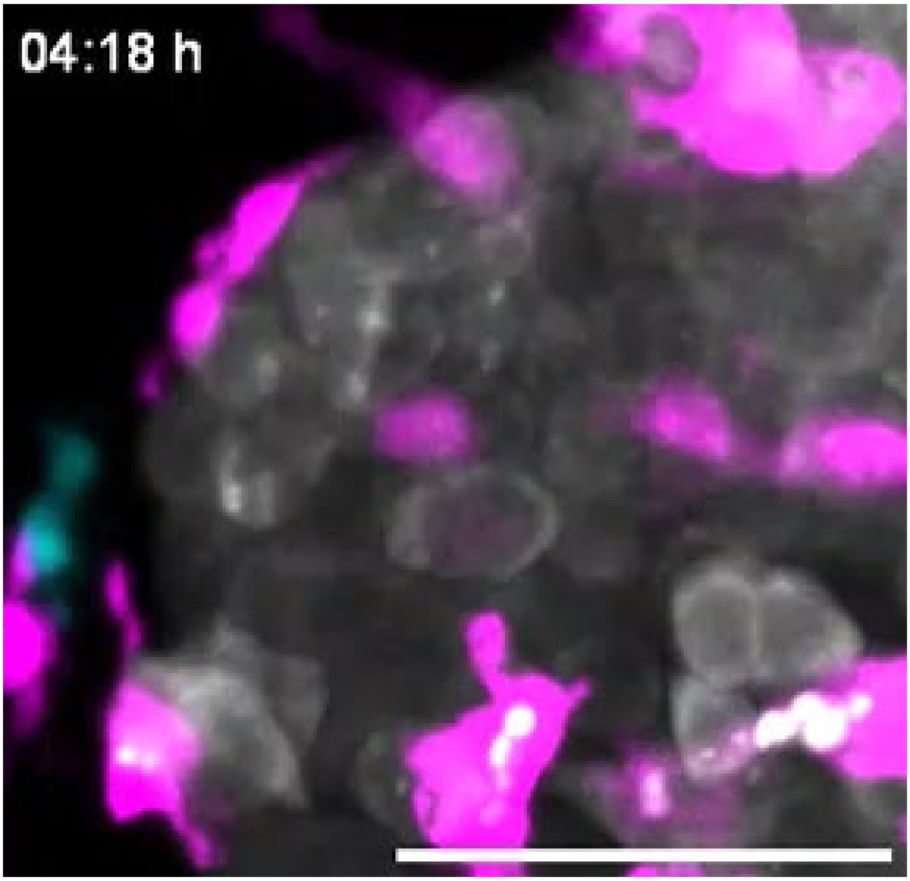


**Movie 8 – Representative time-lapse of a macrophage phagocytosing a tumor cell.** SW620 xenograft generated in Tg(mpeg1:mCherry-F, mpx:GFP-F) larvae showing tumor cells (gray), macrophages (magenta), and neutrophils (cyan). The tumor was acquired overnight between 1 and 2 dpi, in stacks with 3-µm intervals every 3 minutes, and is shown as a maximum projection. Scale bar: 50 µm. Available from the BioImage Archive (accession S-BIAD3811).

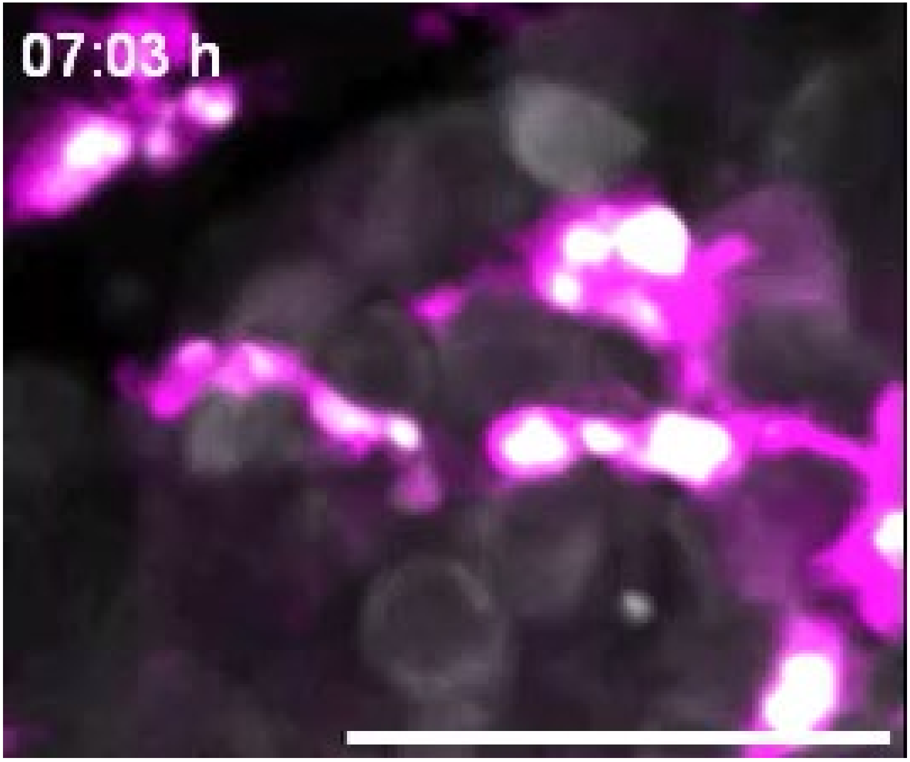


**Movie 9 – Representative time-lapse of a macrophage-macrophage touching event.** SW620 xenograft generated in Tg(mpeg1:mCherry-F, mpx:GFP-F) larvae showing tumor cells (gray), macrophages (magenta), and neutrophils (cyan). The tumor was acquired overnight between 1 and 2 dpi, in stacks with 3-μm intervals every 3 minutes, and is shown as a maximum projection. Scale bar: 50 μm. Available from the BioImage Archive (accession S-BIAD3811).

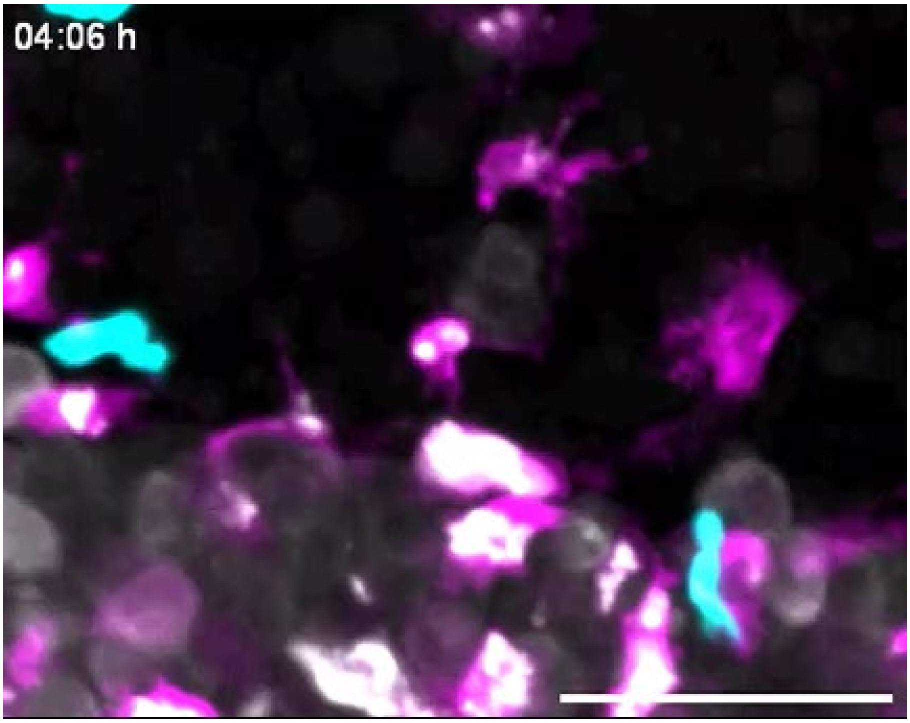


**Movie 10 – Representative time-lapse of a macrophage-neutrophil touching event.** SW620 xenograft generated in Tg(mpeg1:mCherry-F, mpx:GFP-F) larvae showing tumor cells (gray), macrophages (magenta), and neutrophils (cyan). The tumor was acquired overnight between 1 and 2 dpi, in stacks with 3-μm intervals every 3 minutes, and is shown as a maximum projection. Scale bar: 50 μm. Available from the BioImage Archive (accession S-BIAD3811).

### Supplementary tables

**Table S1 –** Sequences of sgRNAs used for CRISPR/Cas9 *CD24* knockout in SW620 cells.

| Gene | sgRNA direction | Sequence (5'-3') | PAM |
|---|---|---|---|
| CD24 | Forward | GAGUACUUCCAACUCUGGGU | TGG |
| | Reverse | ACCCAGAGUUGGAAGUACUC | TGG |

**Table S2 -** Sequences of the 5' and 3' arms for HDR plasmid insertion into the CD24 gene.

| Arm | Sequence (5'-3') |
|---|---|
| 5' | ATTCTAGTTCCATAAGTCTCAGTTACAATATTTAAAAATCCCCATTTTTTTGTGCAACTTTAAAATCTC<br>CCCCTTTCTGGGAAAATAGACCTTATTGCAATGCCATTAAGTCAATATAATCTTGGAAACTGAATTAG<br>TTAACCTGGTCCTTTTTTCCTTTGCAAAGAAATAAAAACAGGAGTTATAGTTATTGAATAACTTTTTTG<br>TTCTTGTTGCCACTTGGCATTTTTGAGGCATCTCTAGGTAGTTTGTTTTGAAACTAAAGAGAATGACC<br>TTGGTGGGTTGAGCAAGAATTCAGAAGTTAATGATGTTGGGTAAGAGAACAATGGTAAGAGAGCAAT<br>CTAAGAATATATCACCTACTTTAATTTTATATGAGAGTACATGGAGGTAGCTGTGATGTGGAAATGTA<br>TCCATGTAACTTTTTATGTATTTTAGATTTATTCCAGTGAAACAACAACTGGAACTT |
| 3' | CAAATCCAACTAATGCCACCACCAAGGCGGCTGGTGGTGCCCTGCAGTCAACAGCCAGTCTCTTCG<br>TGGTCTCACTCTCTCTTCTGCATCTCTACTCTTAAGAGACTCAGGCCAAGAAACGTCTTCTAAATTTC<br>CCCATCTTCTAAACCCAATCCAAATGGCGTCTGGAAGTCCAATGTGGCAAGGAAAAACAGGTCTTCA<br>TCGAATCTACTAATTCCACACCTTTTATTGACACAGAAAATGTTGAGAATCCCAAATTTGATTGATTTG<br>AAGAACATGTGAGAGGTTTGACTAGATGATGGATGCCAATATTAAATCTGCTGGAGTTTCATGTACA<br>AGATGAAGGAGAGGCAACATCCAAAATAGTTAAGACATGATTTCCTTGAATGTGGCTTGAGAAATAT<br>GGACACTTAATACTACCTTGAAAATAAGAATAGAAATAAAGGATGGGATTGTGGAATGGAGATTCAG<br>TTTTCATTTGGTTCATTAATTCTATAAGGCCATAAAACAGGTAATATAAAAAGCTTCCATGATTCTATT<br>TATATGTACATGAGAAGGAACTTCCAGGTGTTACTGTAATTCCTCAA |

**Table S3 –** List of primers used for CD24 sequencing and RT-qPCR.

| Gene | Primer direction | Sequence (5'-3') | Purpose |
|---|---|---|---|
| Puromycin | Forward | GAGGCCGACAAAGAGACCTACG | PCR amplification for Sanger sequencing of edited CD24 KO cell line. |
| CD24 | Reverse | AGAGTGAGACCACGAAGAGAC | |
| CD24 | Forward | GCTCCTACCCACGCAGATTT | CD24 expression quantification by RT-qPCR. |
| | Reverse | GAGACCACGAAGAGACTGGC | |
| EEF1A1 | Forward | ATCCACCTTTGGGTCGCTTT | Housekeeping gene for RT-qPCR. |
| | Reverse | CAGCCTTCTTGTCCACTGCT | |

**Table S4 –** List of antibodies used for Western Blot.

| Antibody | Species | Dilution | Solvent | Brand | Reference |
|---|---|---|---|---|---|
| Anti-CD24 (SN3) | Mouse | 1:500 | 5% milk in TBST | Thermo Fisher Scientific | MA5-11828 |
| Anti-Histone H3 | Rabbit | 1:2000 | 2% BSA in TBST | Cell Signaling | 9715 |
| HRP-conjugated Anti-Mouse IgG | Goat | 1:5000 | 5% milk in TBST | Jackson ImmunoResearch | 115-035-003 |
| HRP-conjugated Anti-Rabbit IgG | Goat | 1:5000 | 5% milk in TBST | Jackson ImmunoResearch | 111-035-003 |

**Table S5 –** List of antibodies used for immunofluorescence of *in vitro* cultures.

| Antibody | Species | Dilution | Brand | Reference |
|---|---|---|---|---|
| Anti-Cleaved Caspase-3 (Asp175) | Mouse | 1:200 | Cell Signalling | 9661 |
| Anti-phospho-Histone H3 (Ser10) | Mouse | 1:200 | Sigma-Aldrich | 05-806 |
| Anti-Phalloidin AlexaFluor™ 488 | - | 1:500 | Thermo Fisher Scientific | A12379 |
| Anti-Mouse IgG Dylight™ 594 | Goat | 1:500 | Thermo Fisher Scientific | 35510 |
| Anti-Rabbit IgG Dylight™ 650 | Goat | 1:500 | Thermo Fisher Scientific | 84546 |
| DAPI | - | 0.00015% in ddH20 | Sigma-Aldrich | 10236276001 |

**Table S6 –** List of antibodies used for immunofluorescence of zebrafish xenografts.

| Antibody | Species | Dilution | Brand | Reference |
|---|---|---|---|---|
| Anti-Cleaved Caspase-3 (Asp175) | Rabbit | 1:100 | Cell Signaling | 9661 |
| Anti-mCherry | Rabbit | 1:100 | Abcam | ab167453 |
| Anti-GFP | Mouse | 1:100 | Roche | ab1218 |
| Anti-Rabbit IgG AlexaFluor™ 488 | Goat | 1:400 | Thermo Fisher Scientific | A11034 |
| Anti-Rabbit IgG DyLight™ 594 | Goat | 1:400 | Thermo Fisher Scientific | 35560 |
| Anti-Rabbit IgG Dylight™ 650 | Goat | 1:400 | Thermo Fisher Scientific | 84546 |
| Anti-Mouse IgG Dylight™ 488 | Goat | 1:400 | Thermo Fisher Scientific | 35502 |
| DAPI | - | 50 µg/ml | Sigma-Aldrich | 10236276001 |

**Table S7 –** Accession numbers and stable identifiers for CD24 proteins and candidate loci. GeneID numbers identify loci and are not nucleotide or protein sequence accessions. LOC symbols correspond to the respective NCBI GeneID numbers. * The reconstructed 261-nt ORF occurs exactly in LOC108716849 transcripts XR_001935769.1 and XR_001935769.2 and encodes the 86-aa GenBank protein OCT80406.1. On CM004474.1, its CDS comprises joined segments 80364850–80364912 and 80366304–80366501. ** The ZFIN records link si:ch211-214p16.1 and si:ch211-214p16.2 to the Ensembl GRCz11 identifiers ENSDARG00000044129 and ENSDARG00000094975, respectively. ENSLOCG00000022258 has the associated transcript ENSLOCT00000027694.1.

| Species | Gene or candidate | Accession or stable identifier | Database and sequence information |
|---|---|---|---|
| Homo sapiens | CD24 isoform a | NP_001278666.1 | NCBI RefSeq protein; 80 aa |
| Homo sapiens | CD24 isoform b | NP_001278668.1 | NCBI RefSeq protein; 129 aa |
| Mus musculus | Cd24a | NP_033976.1 | NCBI RefSeq protein; isoform 2 precursor; 76 aa |
| Gallus gallus | CD24 | NP_001264664.2 | NCBI RefSeq protein; precursor; 87 aa |
| Gallus gallus | LOC124417814 | GeneID: 124417814 | NCBI Gene; ncRNA locus; candidate ORF translated for analysis |
| Xenopus laevis* | LOC108716849 | GeneID: 108716849 | NCBI Gene ncRNA locus; reconstructed ORF matches the GenBank protein exactly; 86 aa |
| Danio rerio** | si:ch211-214p16.1 | ZDB-GENE-030616-606 | ZFIN gene identifier; candidate ORF translated for analysis |
| Danio rerio** | si:ch211-214p16.2 | ZDB-GENE-030616-607 | ZFIN gene identifier; candidate ORF translated for analysis |
| Lepisosteus oculatus | Candidate locus | ENSLOCG00000022258 | Ensembl gene identifier; lincRNA locus; candidate ORF translated for analysis corresponds to Refseq protein accession nr. XP_004082723.1 |
| Oryzias latipes | LOC101171712 | GeneID: 101171712 | NCBI Gene; protein-coding candidate locus |
| Oryzias melastigma | FQA47_024073 | KAF6719451.1 | GenBank protein; hypothetical protein; 66 aa |
| Acipenser ruthenus | LOC117402723 | GeneID: 117402723 | NCBI Gene; ncRNA locus; candidate ORF translated for analysis |
| Callorhinchus milii | LOC121851366 | GeneID: 121851366 | NCBI Gene; ncRNA locus; candidate ORF manually curated and translated |

**Table S8 –** List of putative zebrafish orthologs to human SIGLEC10 and respective identifiers from relevant biological databases: ZFIN for zebrafish genes, Ensembl for transcript information, and UniProt for protein sequences.

| Protein | Organism | Size | ZFIN ID | Ensembl | UniProt |
|---|---|---|---|---|---|
| SIGLEC10 | Human | 697aa | N/A | ENST00000339313.10 | Q96LC7 |
| mag | Zebrafish | 653aa | ZDB-GENE-041217-24 | ENSDART00000165162.2 | F1QSA4 |
| siglec15l | Zebrafish | 347aa | ZDB-GENE-100922-281 | ENSDART00000127243.3 | A7IT78 |
| si:dkey-24p1.7 | Zebrafish | 598aa | ZDB-GENE-121214-215 | ENSDART00000152404.2 | A0A8M2BKP6 |

**Table S9 –** FASTA sequence of each hSIGLEC10 candidate ortholog, extracted from UniProt.

```
>human_siglec10_Q96LC7
MLLPLLLSSLLGGSQAMDGRFWIRVQESVMVPEGLCISVPCSFSYPRQDWTGSTPAYGYWFKAVTETT
KGAPVATNHQSREVEMSTRGRFQLTGDPAKGNCSLVIRDAQMQDESQYFFRVERGSYVRYNFMNDGF
FLKVTALTQKPDVYIPETLEPGQPVTVICVFNWAFEECPPPSFSWTGAALSSQGTKPTTSHFSVLSFTPR
PQDHNTDLTCHVDFSRKGVSAQRTVRLRVAYAPRDLVISISRDNTPALEPQPQGNVPYLEAQKGQFLRL
LCAADSQPPATLSWVLQNRVLSSSHPWGPRPLGLELPGVKAGDSGRYTCRAENRLGSQQRALDLSVQ
YPPENLRVMVSQANRTVLENLGNGTSLPVLEGQSLCLVCVTHSSPPARLSWTQRGQVLSPSQPSDPGV
LELPRVQVEHEGEFTCHARHPLGSQHVSLSLSVHYSPKLLGPSCSWEAEGLHCSCSSQASPAPSLRW
WLGEELLEGNSSQDSFEVTPSSAGPWANSSLSLHGGLSSGLRLRCEAWNVHGAQSGSILQLPDKKGLI
STAFSNGAFLGIGITALLFLCLALIIMKILPKRRTQTETPRPRFSRHSTILDYINVVPTAGPLAQKRNQKATP
NSPRTPLPPGAPSPESKKNQKKQYQLPSFPEPKSSTQAPESQESQEELHYATLNFPGVRPRPEARMPK
GTQADYAEVKFQ
```

```
>zebrafish_mag_F1QSA4
MKGLELLLLPLLLLLKDASAQWNVWMPRDISAMTNSCVVIPCSFTYPSGIRPYRGVHGIWYFGHPYPQLF
PPVVLKSRTDIVHESYKGRTKLLGDLTQKNCTLLISNIGVEHSGKYYFRADLGGANIYTFPDFTKLQVLDQ
PNIDVPEEIVSDQSLDLTCYVPDNCPDMSPEIHWMYTDYLPDPVFTPDQVEEGNTAVLSSTLTFTPKPMH
NGQLLGCRVNFPNTTYVYERLITLDIRYAPRTVWVNVSQEVMEGSSVVLHCDVDSNPAPMITWYFGDKE
LMSETASNSSLSLENLTPEQEGVYTCVGDNGYGNMNTSMYLAVNYPPREPWINESLTVLEGSSVSLQC
TSKGNPMPTLTWLKDGELVGTITAEEGSVLELHEIMPQADGVYRCLAENEHGRASSSLNITVEFAPVLLD
DSKCTIVREGVQCVCIASGNPEPAIEFYLPDLNVTINNSNNRFNYYTHSDGYTSTGVIKLQDKGERGNNG
DTAVHVHCSITNIYGSETIRLELQQEKKYMMAVIVGTIGGVAVIAFIIAAVRYVGQNNKKENGNPGQDVGS
KVENPSMFYSAVKKDKQSLRKKVLKTELLGSKFNSILEEGTGDDSDYQSVGPMAGMERQELNYAALEFL
HGRHREGVFRRADGDGSDYTEIKAK
```

```
>zebrafish_siglec15l_A7IT78
MQGFVFGFLTFICCLKGLTCSDWTINGQTKVNGSLGQNVILPCSFTHPKQDTYTGNIIVMWMQENRKEP
VFQCIMSNKTDSQDGNCSLKPSKRFWLHGNPRKRDISLAITNLQLSDAAKYACRLMLDYDKVQRATELIV
NAPARILILSQYLELTSHALLVKCIAQGNPVPDVKWTLSSGQNASATTERHKYMVMSSVQFSEQDVFTCQ
AVNSLGRAQKTFPPDPRSCFAPLAATGVLAVVLVLGLLLFIWDVIRKRQSEQGAMEHYLGAQQAKSSIYE
ANEMVYANVRTLPPDASAQGKMYSTWPPSEEAKAEVNPRFQSQRRATSKVRPATAATTPTAPVESERY
```

```
>zebrafish_si:dkey-24p1.7_A0A8M2BKP6
MKIISLCTIIFMLVFNVLCEENADNGWNISFSPREVTAVKGLCAHISCTFTYPVAENPIQSVIWQSCVAKDK
CNLIFQNNVEKNKLKKNELGHIKMLEPDLSKNNCSIILKDIKENEKEYTFRIKAKQQYTFNPTVKISIKDEPT
LVVPPLSEKVEVNLTCSAPFPCPETPPVITWWIKTKEENYIKLDNNKITQVHSEHVYYSYLTLIPTSDQHDG
TVGCDVRYGNKKINTNNTLEVKYVQALQIVGENTLMEGDTLNLTCTFRSHPPASNPVWRFNGNTDNLNT
QTSAGTLMIANVGKEHAGIYVCEMTYMNKTLNASITINITEVNILGENRLKKGDTLNLTCSFKNHPQSSSN
PVWSFNGDADKLKNQASAANLIIDNVTKEHAGIYACEMTYMKKTLNASITVDITEHDMGRNNTRAGVLDF
LLSIPNIFTFVAGMAFSALIFSVILCCWVCCHRGKKQKVPTANPDAEINLETVQTDVAQNGTTEQTPLHEQ
PDGETPKTPATPTDGAEEDEALGTEAREVDYASIDYSLLKDKPPEEAETEPTDTDYAEIKKDRTKDWKDT
EDPQDGDEQVETFDQKEMGAEEELYSQVPQ
```

**Table S10 –** Clinical and demographic characteristics of TCGA colorectal cancer patients from the COAD and READ datasets.

| Characteristic | All Patients | CD24 Expression | |
|---|---|---|---|
| | | Low | High |
| **N** | **601** | **299** | **302** |
| Age, median (range), years | 68.0 (31-90) | 67.0 (33-90) | 68.0 (31-90) |
| **Cancer type, n (%)** | | | |
| Colon (COAD) | 444 (73.9) | 227 (75.9) | 217 (71.9) |
| Rectal (READ) | 157 (26.1) | 72 (24.1) | 85 (28.1) |
| **Sex, n (%)** | | | |
| Male | 317 (52.7) | 156 (52.2) | 161 (53.3) |
| Female | 284 (47.3) | 143 (47.8) | 141 (46.7) |
| **Race, n (%)** | | | |
| White | 276 (45.9) | 123 (41.1) | 153 (50.7) |
| Black/African American | 65 (10.8) | 32 (10.7) | 33 (10.9) |
| Asian | 12 (2.0) | 7 (2.3) | 5 (1.7) |
| American Indian/Alaska Native | 1 (0.2) | 1 (0.3) | 0 (0.0) |
| Not reported | 247 (41.1) | 136 (45.5) | 111 (36.8) |
| **Stage, n (%)** | | | |
| Stage I | 105 (17.5) | 52 (17.4) | 53 (17.5) |
| Stage II | 227 (37.8) | 113 (37.8) | 114 (37.7) |
| Stage III | 179 (29.8) | 89 (29.8) | 90 (29.8) |
| Stage IV | 90 (15.0) | 45 (15.1) | 45 (14.9) |
| **Vital status, n (%)** | | | |
| Alive | 481 (80.0) | 237 (79.3) | 244 (80.8) |
| Deceased | 120 (20.0) | 62 (20.7) | 58 (19.2) |
| **Overall Survival** | | | |
| Median follow-up, months | 21.4 | 19.3 | 23.0 |
| Events, n (%) | 120 (20.0) | 62 (20.7) | 58 (19.2) |
| **Progression-Free Interval** | | | |
| Median follow-up, months | 19.0 | 17.6 | 20.0 |
| Events, n (%) | 156 (26.0) | 79 (26.4) | 77 (25.5) |
